# Population-specific causal disease effect sizes in functionally important regions impacted by selection

**DOI:** 10.1101/803452

**Authors:** Huwenbo Shi, Steven Gazal, Masahiro Kanai, Evan M. Koch, Armin P. Schoech, Katherine M. Siewert, Samuel S. Kim, Yang Luo, Tiffany Amariuta, Hailiang Huang, Yukinori Okada, Soumya Raychaudhuri, Shamil R. Sunyaev, Alkes L. Price

## Abstract

Many diseases and complex traits exhibit population-specific causal effect sizes with trans-ethnic genetic correlations significantly less than 1, limiting trans-ethnic polygenic risk prediction. We developed a new method, S-LDXR, for stratifying squared trans-ethnic genetic correlation across genomic annotations, and applied S-LDXR to genome-wide association summary statistics for 31 diseases and complex traits in East Asians (EAS) and Europeans (EUR) (average *N*_EAS_=90K, *N*_EUR_=267K) with an average trans-ethnic genetic correlation of 0.85 (s.e. 0.01). We determined that squared trans-ethnic genetic correlation was 0.82× (s.e. 0.01) smaller than the genome-wide average at SNPs in the top quintile of background selection statistic, implying more population-specific causal effect sizes. Accordingly, causal effect sizes were more population-specific in functionally important regions, including conserved and regulatory regions. In analyses of regions surrounding specifically expressed genes, causal effect sizes were most population-specific for skin and immune genes and least population-specific for brain genes. Our results could potentially be explained by stronger gene-environment interaction at loci impacted by selection, particularly positive selection.

## Introduction

Trans-ethnic genetic correlations are significantly less than 1 for many diseases and complex traits,^1–6^ implying that population-specific causal disease effect sizes contribute to the incomplete portability of genome-wide association study (GWAS) findings and polygenic risk scores to non-European populations.^6–12^ However, current methods for estimating genome-wide trans-ethnic genetic correlations assume the same trans-ethnic genetic correlation for all categories of SNPs,^2,5,13^ providing little insight into why causal disease effect sizes are population-specific. Understanding the biological processes contributing to population-specific causal disease effect sizes can help inform polygenic risk prediction in non-European populations and alleviate health disparities.^6,14,15^

Here, we introduce a new method, S-LDXR, for estimating enrichment of stratified squared trans-ethnic genetic correlation across functional categories of SNPs using GWAS summary statistics and population-matched linkage disequilibrium (LD) reference panels (e.g. the 1000 Genomes Project (1000G)^16^); we stratify the *squared* trans-ethnic genetic correlation across functional categories to robustly handle noisy heritability estimates. S-LDXR analyzes GWAS summary statistics of HapMap3^17^ SNPs with minor allele frequency (MAF) greater than 5% in both East Asian (EAS) and European (EUR) populations (*regression SNPs*) to draw inferences about causal effects of all SNPs with MAF greater than 5% in both populations (*heritability SNPs*). We confirm that S-LDXR yields robust estimates in extensive simulations. We apply S-LDXR to 31 diseases and complex traits with GWAS summary statistics available in both East Asian (EAS) and European (EUR) populations, leveraging recent large studies in East Asian populations from the CONVERGE consortium and Biobank Japan;^18–20^ we analyze a broad set of genomic annotations from the baseline-LD model,^21–23^ as well as tissue-specific annotations based on specifically expressed gene sets.^24^ Most results are meta-analyzed across the 31 traits to maximize power (analogous to ref.^21–23^), as we expect to see similar patterns of enrichment/depletion across traits (even though the underlying biological processes differ across traits). We also investigate trait-specific enrichments/depletions for the tissue-specific annotations (analogous to ref.^24^).

## Results

### Overview of methods

Our method (S-LDXR) for estimating stratified trans-ethnic genetic correlation is conceptually related to stratified LD score regression^21,22^ (S-LDSC), a method for partitioning heritability from GWAS summary statistics while accounting for LD. S-LDXR determines that a category of SNPs is enriched for trans-ethnic genetic covariance if SNPs with high LD to that category have higher product of Z-scores than SNPs with low LD to that category. Unlike S-LDSC, S-LDXR models per-allele effect sizes (accounting for differences in MAF between populations).

In detail, the product of Z-scores of SNP *j* in two populations, *Z*_1*j*_*Z*_2*j*_, has the expectation

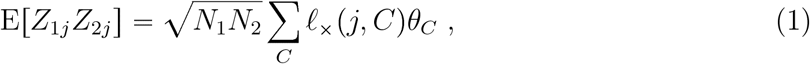

where *N*_*p*_ is the sample size for population *p*; *ℓ*×(*j, C*) = Σ_*k*_ *r*_1*jk*_*r*_2*jk*_σ_1*j*_σ_2*j*_a_*C*_(*k*) is the transethnic LD score of SNP *j* with respect to annotation *C*, whose value for SNP *k*, *a*_*C*_(*k*), can be either binary or continuous; *r*_*pjk*_ is the LD (Pearson correlation) between SNP *j* and *k* in population *p*; *σ*_*pj*_ is the standard deviation of SNP *j* genotypes in population *p*; and *θ*_*C*_ represents the per-SNP contribution to trans-ethnic genetic covariance of the *per-allele* causal disease effect size of annotation *C*. Here, *r*_*pjk*_ and *σ*_*pj*_ can be estimated from population matched reference panels (e.g. 1000 Genomes Project^16^). We estimate *θ*_*C*_ for each annotation *C* using weighted least square regression. Subsequently, we estimate the trans-ethnic genetic covariance of each binary annotation *C* (*ρ*_*g*_(*C*)) by summing trans-ethnic genetic covariance of each SNP in annotation *C* as Σ_*j*∈*C*_ (Σ_*C′*_ *a*_*C′*_ (*j*)*θ*_*C′*_), using coefficients (*θ*_*C′*_) for all binary and continuous-valued annotations *C′* included in the analysis; the heritabilities in each population 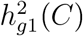 and 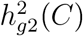 are estimated analogously. We then estimate the stratified *squared* trans-ethnic genetic correlation, defined as

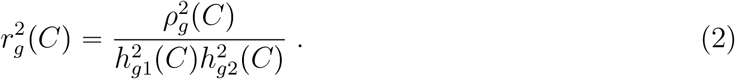

We define the enrichment/depletion of squared trans-ethnic genetic correlation as 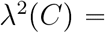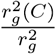, where is 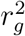 the genome-wide squared trans-ethnic genetic correlation; *λ*^2^ (*C*) can be meta-analyzed across traits with different 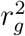. S-LDXR analyzes GWAS summary statistics of common HapMap3^17^ SNPs (*regression SNPs*) to estimate *λ*^2^(*C*) for (causal effects of) all common SNPs (*heritability SNPs*). Further details (quantities estimated, analytical bias correction, shrinkage estimator to reduce standard errors, estimation of standard errors, significance testing, and factors impacting power) of the S-LDXR method are provided in the Methods section; we have publicly released open-source software implementing the method (see URLs).

We apply S-LDXR to 62 annotations (defined in both EAS and EUR populations) from our baseline-LD-X model (Methods, Table S1, Figures S1, S2), primarily derived from the baseline-LD model^21–23^ (v1.1; see URLs). We have publicly released all baseline-LD-X model annotations and LD scores for EAS and EUR populations (see URLs).

### Simulations

We evaluated the accuracy of S-LDXR in simulations using genotypes that we simulated using HAPGEN2^25^ from phased haplotypes of 481 EAS and 489 EUR individuals from the 1000 Genomes Project^16^, preserving population-specific MAF and LD patterns (18,418 simulated EAS-like and 36,836 simulated EUR-like samples, after removing genetically related samples, ratio of sample sizes similar to empirical data; ~2.5 million SNPs on chromosomes 1 – 3) (Methods); we did not have access to individual-level EAS data at sufficient sample size to perform simulations with real genotypes. For each population, we randomly selected a subset of 500 simulated samples to serve as the reference panel for estimating LD scores. We performed both null simulations (heritable trait with functional enrichment but no enrichment/depletion of squared trans-ethnic genetic correlation; *λ*^2^(*C*) = 1) and causal simulations (*λ*^2^(*C*) ≠ 1). In our main simulations, we randomly selected 10% of the SNPs as causal SNPs in both populations, set genome-wide heritability to 0.5 in each population, and adjusted genome-wide genetic covariance to attain a genome-wide *r*_*g*_ of 0.60 (unless otherwise indicated). In the null simulations, we used heritability enrichments from analyses of real traits in EAS samples to specify per-SNP causal effect size variances and covariances. In the causal simulations, we directly specified per-SNP causal effect size variances and covariances to attain *λ*^2^(*C*) ≠ 1 values from analyses of real traits, as these were difficult to attain using the heritability and trans-ethnic genetic covariance enrichments from analyses of real traits.

First, we assessed the accuracy of S-LDXR in estimating genome-wide trans-ethnic genetic correlation (*r*_*g*_); we note that S-LDXR does not use the shrinkage estimator for genome-wide estimates. Across a wide range of simulated *r*_*g*_ values (0.0 to 1.0), S-LDXR yielded approximately unbiased estimates and well-calibrated jackknife standard errors (Table S3, Figure S3).

Second, we assessed the accuracy of S-LDXR in estimating *λ*^2^(*C*) in quintiles of the 8 continuous-valued annotations of the baseline-LD-X model. We performed both null simulations (*λ*^2^(*C*) = 1) and causal simulations (*λ*^2^(*C*) ≠ 1). Results are reported in Figure 1a and Tables S4 – S9. In both null and causal simulations, S-LDXR yielded approximately unbiased estimates of *λ*^2^(*C*) for most annotations, validating our analytical bias correction. As a secondary analysis, we tried varying the S-LDXR shrinkage parameter, *α*, which has a default value of 0.5. In null simulations, results remained approximately unbiased; in causal simulations, reducing *α* led to less precise (but less biased) estimates of *λ*^2^(*C*), whereas increasing *α* biased results towards the null (*λ*^2^(*C*) = 1), demonstrating a bias-variance tradeoff in the choice of *α* (Figure S4, Tables S5, S8). Results were similar at other values of the proportion of causal SNPs (1% and 100%; Tables S4, S6, S7, S9). We also confirmed that S-LDXR produced well-calibrated jackknife standard errors (Tables S4-S9).

**Figure 1:**
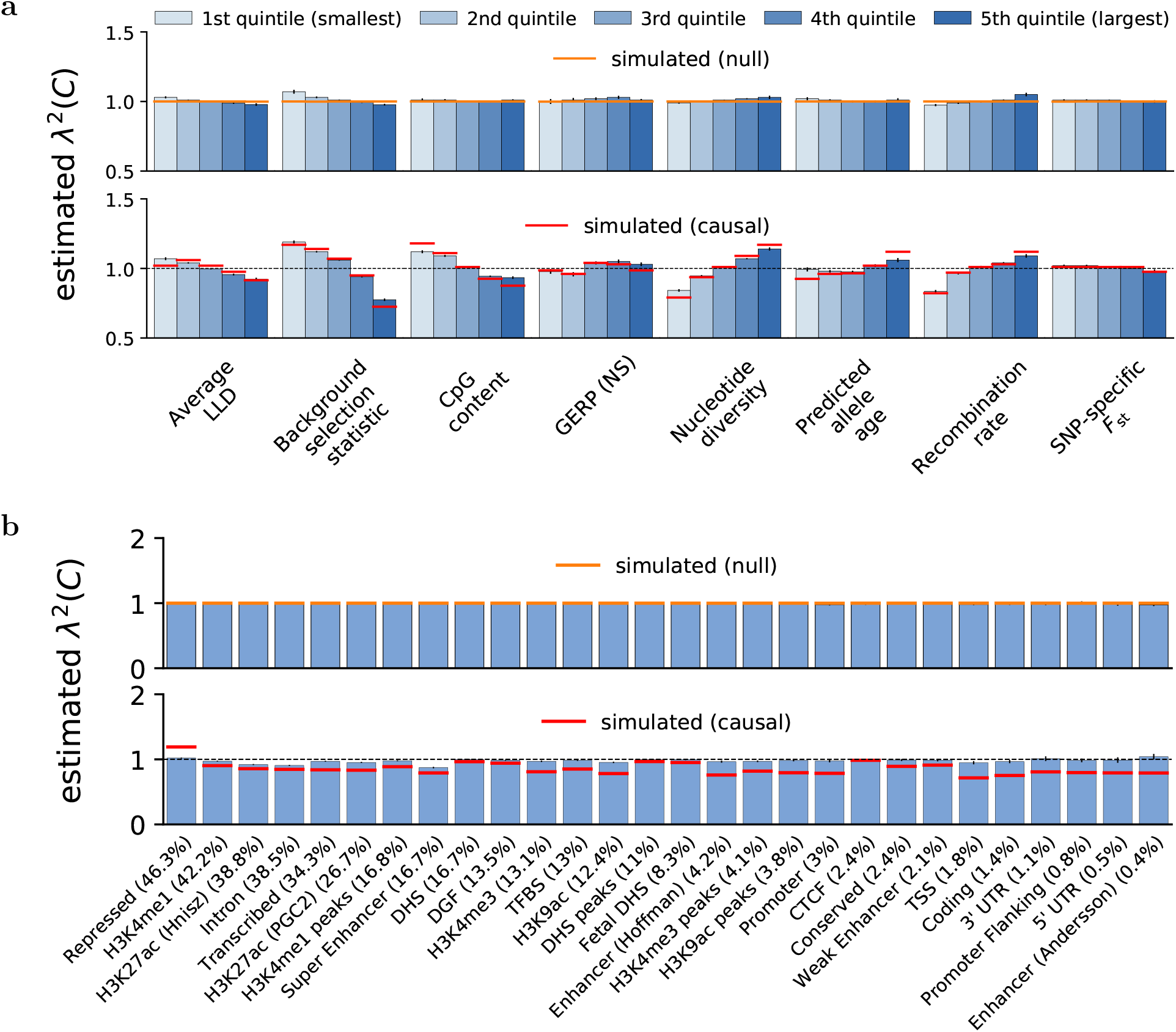
Accuracy of S-LDXR in null and causal simulations. We report estimates of the enrichment/depletion of squared trans-ethnic genetic correlation (*λ*^2^(*C*)) in both null and causal simulations, for (a) quintiles of 8 continuous-valued annotations and (b) 28 main binary annotations (sorted by proportion of SNPs, displayed in parentheses). Results are averaged across 1,000 simulations. Error bars denote ±1.96× standard error. Numerical results are reported in Table S5 and S8.

Third, we assessed the accuracy of S-LDXR in estimating *λ*^2^(*C*) for the 28 main binary annotations of the baseline-LD-X model (inherited from the baseline model of ref.^21^). We discarded *λ*^2^(*C*) estimates with the highest standard errors (top 5%), as estimates with large standard errors (which are particularly common for annotations of small size) are uninformative for evaluating unbiasedness of the estimator (in analyses of real traits, trait-specific estimates with large standard errors are retained, but contribute very little to meta-analysis results, and would be interpreted as inconclusive when assessing trait-specific results). Results are reported in Figure 1b and Tables S5, S8. In null simulations, S-LDXR yielded unbiased estimates of *λ*^2^(*C*), further validating our analytical bias correction. In causal simulations, estimates were biased towards the null (*λ*^2^(*C*) = 1) – particularly for annotations of small size (proportion of SNPs < 1%) – due to our shrinkage estimator; increasing the shrinkage parameter above its default value of 0.5 further biased the estimates towards the null (*λ*^2^(*C*) = 1) in causal simulations (Tables S7, S8, S9). To ensure robust estimates, we focus on the 20 main binary annotations of large size (> 1% of SNPs) in analyses of real traits (see below); although results for these annotations may still be biased towards the null, we emphasize that S-LDXR is unbiased in null data. Results were similar at other values of the proportion of causal SNPs (1% and 100%; Tables S4, S6, S7, S9). We also confirmed that S-LDXR produced well-calibrated jackknife standard errors (Tables S4-S9) and conservative p-values (Figure S7, Table S4-S9).

Fourth, we performed additional null simulations in which causal variants differed across the two populations (Methods). S-LDXR yielded robust estimates of *λ*^2^(*C*), well-calibrated standard errors and conservative p-values in these simulations (Figure S11, Table S12).

Fifth, we performed additional null simulations with annotation-dependent MAF-dependent genetic architectures,^26–28^ defined as architectures in which the level of MAF-dependence is annotation-dependent, to ensure that estimate of *λ*^2^(*C*) remain unbiased. We disproportionately sampled low-frequency causal variants from the top quintile of background selection statistic, and set the variance of per-allele effect sizes of a causal SNP to be inversely proportional to its maximum MAF across both populations (Methods). Results are reported in Figure S8-S10, and Table S10, S11. S-LDXR yielded nearly unbiased estimates of *λ*^2^(*C*) for the 28 binary functional annotations (Figure S8) and nearly unbiased estimates of *λ*^2^(*C*) for most quintiles of continuously valued annotations (Figure S9); estimates were slightly biased estimates in the top and bottom quintile of the average level of LD annotation and the recombination rate annotation, likely due to less accurate reference LD scores at SNPs with extreme levels of LD. We repeated these simulations with 5 MAF bin annotations added to the baseline-LD-X model and obtained similar results (Figure S8a, S9b), supporting our decision not to include MAF bin annotations into the baseline-LD-X model.

Sixth, we performed additional null simulations, in which we increased or decreased the reference panel size from 500 to 250 or 1,000, to assess the impact of reference panel size on the accuracy of S-LDXR (Methods). We simulated GWAS summary statistics based on the baseline-LD-X model as well as the model with annotation-dependent MAF-dependent genetic architectures. We determined that the small systematic biases in null simulations of continuous-valued annotations were on the same order of magnitude as for 500 reference samples (Figures S12, S13 and Table S13 for 250 reference samples; Figures S14, S15 and Table S14 for 1,000 reference samples). We also performed simulations in which we reduced the simulated GWAS sample size by half, from *N*_EAS_=18K, *N*_EUR_=37K to *N*_EAS_=9K, *N*_EUR_=18K (while fixing the reference panel size at 500). We again determined that the small systematic biases were generally on the same order of magnitude as for *N*_EAS_=18K, *N*_EUR_=37K (although estimates were less stable and sometimes subject to larger biases, likely because our analytical bias correction starts to break down when the GWAS has low power) (Figures S16, S17 and Table S15). Although it was not computationally feasible to perform simulations at larger GWAS sample sizes, these analyses do not provide a reason to believe that the small systematic biases that we observed in some of our null simulations of continuously valued annotations would substantially increase at larger GWAS sample sizes.

In summary, S-LDXR produced approximately unbiased estimates of enrichment/depletion of squared trans-ethnic genetic correlation in null simulations, and conservative estimates in causal simulations of both quintiles of continuous-valued annotations and binary annotations.

### Analysis of baseline-LD-X model annotations across 31 diseases and complex traits

We applied S-LDXR to 31 diseases and complex traits with summary statistics in East Asians (average N=90K) and Europeans (average N=267K) available from Biobank Japan, UK Biobank, and other sources (Table S16 and Methods). First, we estimated the trans-ethnic genetic correlation (*r*_*g*_) (as well as population-specific heritabilies) for each trait. Results are reported in Figure S18 and Table S16. The average *r*_*g*_ across 31 traits was 0.85 (s.e. 0.01) (average 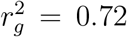 (s.e. 0.02)). 28 traits had *r*_*g*_ < 1, and 11 traits had *r*_*g*_ significantly less than 1 after correcting for 31 traits tested (*P* < 0.05/31); the lowest *r*_*g*_ was 0.34 (s.e. 0.07) for Major Depressive Disorder (MDD), although this may be confounded by different diagnostic criteria in the two populations.^29^ Several other complex traits, including Age at Menopause (*r*_*g*_ = 0.57 (s.e. 0.09)) and LDL (*r*_*g*_ = 0.66 (s.e. 0.11)) also had low trans-ethnic *r*_*g*_, likely due to pervasive gene-environment interaction across the genome. These estimates were consistent with estimates obtained using Popcorn^2^ (Figure S19) and those reported in previous studies.^2,5,6^ We note that our estimates of trans-ethnic genetic correlation for 31 complex traits are higher than those reported for gene expression traits^2^ (average estimate of 0.32, increasing to 0.77 when restricting the analysis to gene expression traits with (cis) heritability greater than 0.2 in both populations), which are expected to have different genetic architectures.

Second, we estimated the enrichment/depletion of squared trans-ethnic genetic correlation (*λ*^2^(*C*)) in quintiles of the 8 continuous-valued annotations of the baseline-LD-X model, meta-analyzing results across traits; these annotations are moderately correlated (Figure 2a and Table S1). We used the default shrinkage parameter (*α* = 0.5) in all analyses. Results are reported in Figure 2b and Table S17. We consistently observed a depletion of 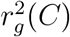 (*λ*^2^(*C*)) < 1, implying more population-specific causal effect sizes) in functionally important regions. For example, we estimated *λ*^2^(*C*) = 0.82 (s.e. 0.01) for SNPs in the top quintile of background selection statistic (defined as 1 - McVicker B statistic / 1000;^30^ see ref.^22^); *λ*^2^(*C*) estimates were less than 1 for 29/31 traits, including 2 traits (Height and EGFR) with two-tailed *p* < 0.05/31. The background selection statistic quantifies the genetic distance of a site to its nearest exon; regions with high background selection statistic have higher per-SNP heritability, consistent with the action of selection, and are enriched for functionally important regions.^22^ We observed the same pattern for CpG content and SNP-specific *F*_st_ (which are positively correlated with background selection statistic; Figure 2a) and the opposite pattern for nucleotide diversity (which is negatively correlated with background selection statistic). We also estimated *λ*^2^(*C*) = 0.87 (s.e. 0.03) for SNPs in the top quintile of average LLD (which is positively correlated with background selection statistic), although these SNPs have *lower* per-SNP heritability due to a competing positive correlation with predicted allele age.^22^ We caution that average LLD was the annotation most susceptible to bias in our simulations; see Simulations. Likewise, we estimated *λ*^2^(*C*) = 0.84 (s.e. 0.02) for SNPs in the *bottom* quintile of recombination rate (which is negatively correlated with background selection statistic), although these SNPs have average per-SNP heritability due to a competing negative correlation with average LLD.^22^ However, *λ*^2^(*C*) < 1 estimates for the bottom quintile of GERP (NS) (which is positively correlated with both background selection statistic and recombination rate) and the middle quintile of predicted allele age are more difficult to interpret. For all annotations analyzed, heritability enrichments did not differ significantly between EAS and EUR, consistent with previous studies.^20,31^ Results were similar at a more stringent shrinkage parameter value (*α* = 1.0; Figure S20), and for a meta-analysis across a subset of 20 approximately independent traits (Methods; Figure S21).

**Figure 2:**
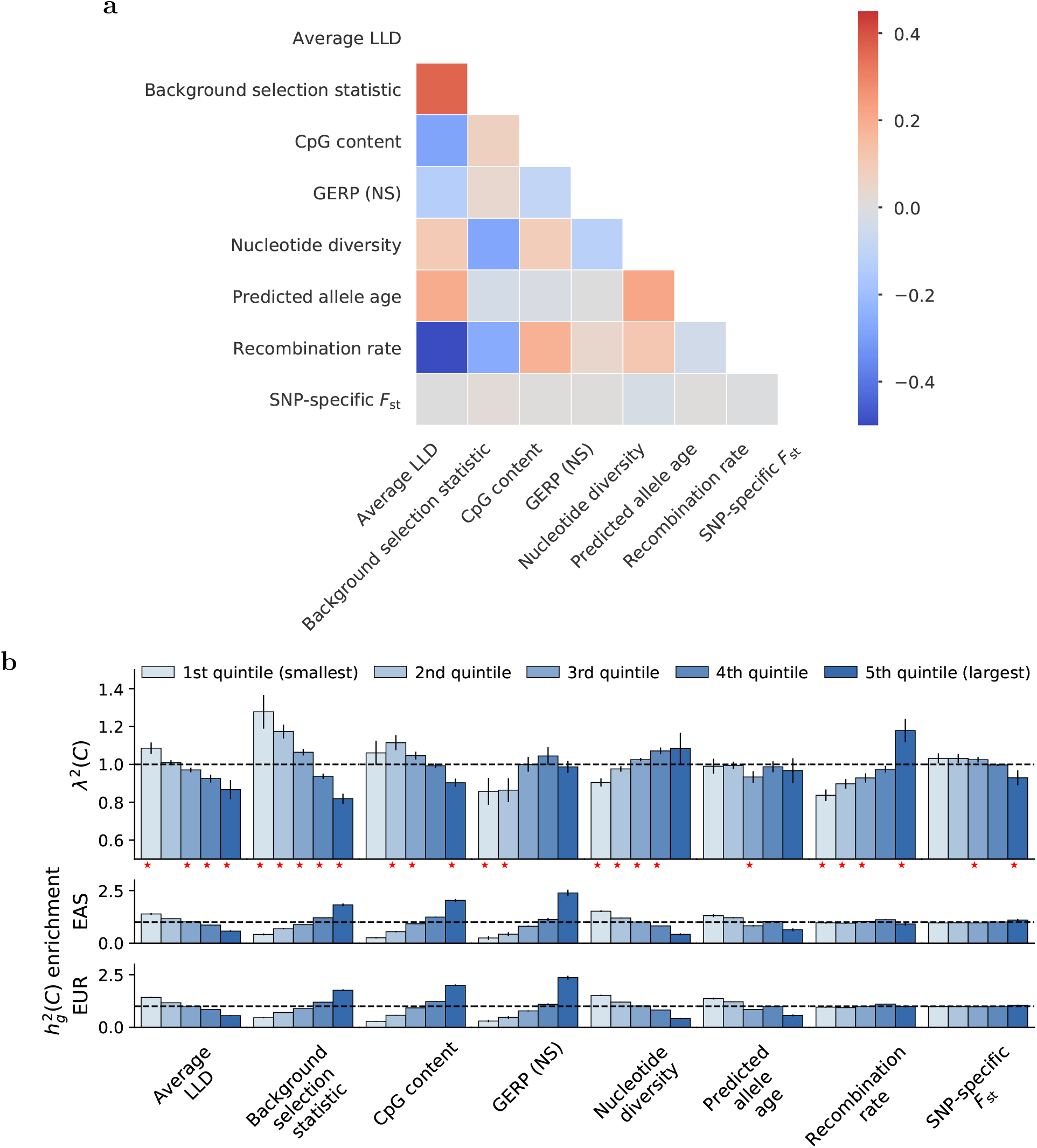
S-LDXR results for quintiles of 8 continuous-valued annotations across 31 diseases and complex traits. (a) We report correlations between each continuous-valued annotation; diagonal entries are not shown. Numerical results are reported in Table S1. (b) We report estimates of the enrichment/depletion of squared trans-ethnic genetic correlation (*λ*^2^ *C*), as well as population-specific estimates of heritability enrichment, for quintiles of each continuous-valued annotation. Results are meta-analyzed across 31 diseases and complex traits. Error bars denote ±1.96× standard error. Red stars (⋆) denote two-tailed p<0.05/40. Numerical results are reported in Table S17.

Finally, we estimated *λ*^2^(*C*) for the 28 main binary annotations of the baseline-LD-X model (Table S1), meta-analyzing results across traits (as we did not observe significant trait-specific enrichment/depletion of squared trans-ethnic genetic correlation for these annotations due to limited power). Results are reported in Figure 3a and Table S18. Our primary focus is on the 20 annotations of large size (> 1% of SNPs), for which our simulations yielded robust estimates; results for remaining annotations are reported in Table S18. We consistently observed a depletion of *λ*^2^(*C*) (implying more population-specific causal effect sizes) within these annotations: 17 annotations had *λ*^2^(*C*) < 1, and 5 annotations had *λ*^2^(*C*) significantly less than 1 after correcting for 20 annotations tested (*P* < 0.05/20). These annotations included Conserved (*λ*^2^(*C*) = 0.93 (s.e. 0.02)), Promoter (*λ*^2^(*C*) = 0.85 (s.e. 0.04)) and Super Enhancer (*λ*^2^(*C*) = 0.93 (s.e. 0.02)), each of which was significantly enriched for per-SNP heritability, consistent with ref.^21^. For all annotations analyzed, heritability enrichments did not differ significantly between EAS and EUR (Figure 3a), consistent with previous studies.^20,31^ Results were similar at a more stringent shrinkage parameter value (*α* = 1.0; Figure S20), and for a meta-analysis across a subset of 20 approximately independent traits (Methods; Figure S22). As a secondary analysis, we also estimated *λ*^2^(*C*) across 10 MAF bin annotations; we did not observe variation in *λ*^2^(*C*) estimates across MAF bins (Table S19), further supporting our decision to not include MAF bin annotations in the baseline-LD-X model.

**Figure 3:**
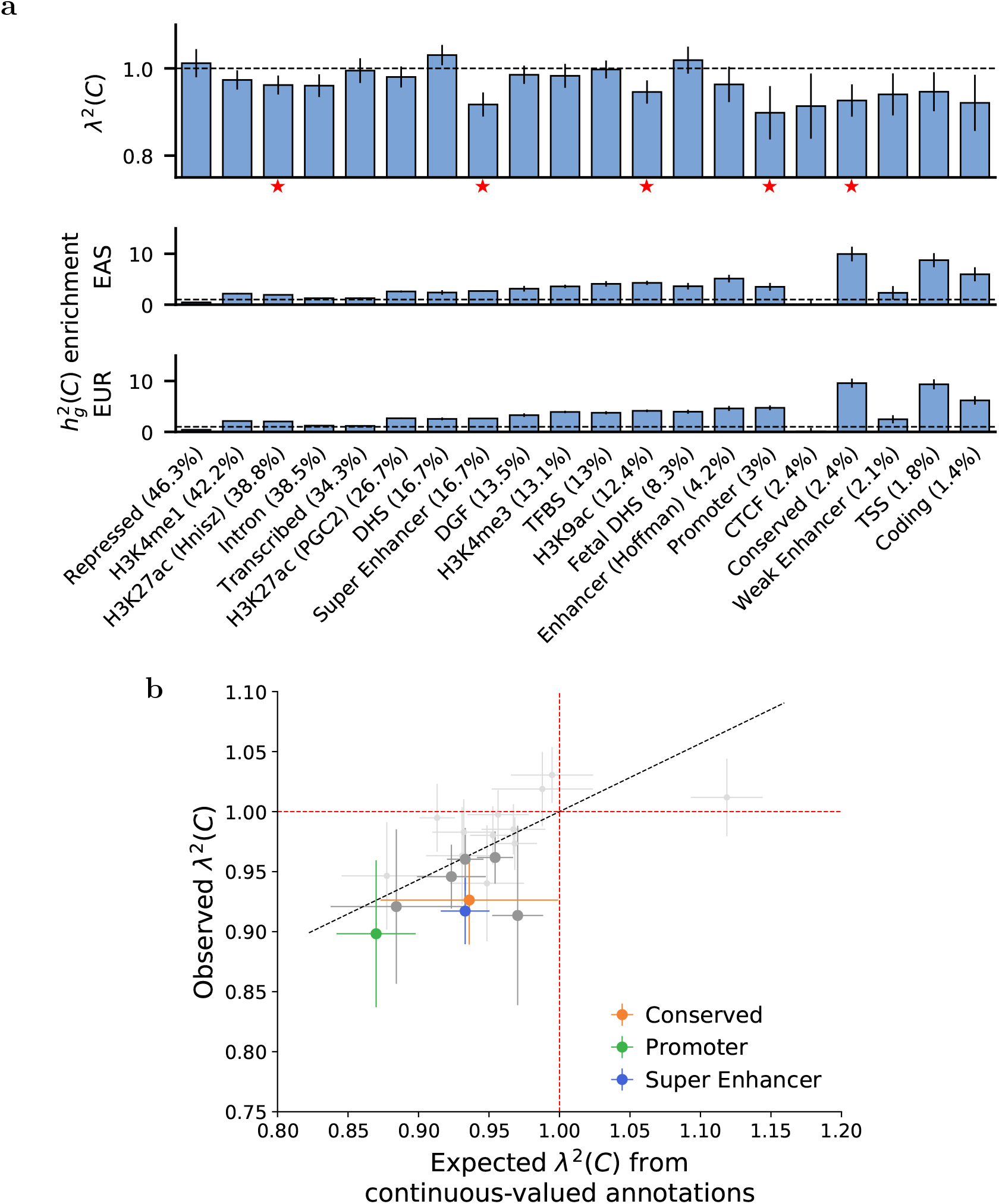
S-LDXR results for 20 binary functional annotations across 31 diseases and complex traits. (a) We report estimates of the enrichment/depletion of squared trans-ethnic genetic correlation (*λ*^2^(*C*)), as well as population-specific estimates of heritability enrichment, for each binary annotation (sorted by proportion of SNPs, displayed in parentheses). Results are meta-analyzed across 31 diseases and complex traits. Error bars denote ±1.96× standard error. Red stars (⋆) denote two-tailed p<0.05/20. Numerical results are reported in Table S18. (b) We report observed *λ*^2^(*C*) vs. expected *λ*^2^(*C*) based on 8 continuous-valued annotations, for each binary annotation. Results are meta-analyzed across 31 diseases and complex traits. Error bars denote 1.96 standard error. Annotations for which *λ*^2^ *C* is significantly different from 1 (p<0.05/20) are denoted in color (see legend) or dark gray. The dashed black line (slope=0.57) denotes a regression of observed *λ*(*C*) 1 vs. expected *λ*(*C*) 1 with intercept constrained to 0. Numerical results are reported in Table S20.

Since the functional annotations are moderately correlated with the 8 continuous-valued annotations (Table S1c, Figure S1), we investigated whether the depletions of squared trans-ethnic genetic correlation (*λ*^2^(*C*) < 1) within the 20 binary annotations could be explained by the 8 continuous-valued annotations. For each binary annotation, we estimated its expected *λ*^2^(*C*) based on values of the 8 continuous-valued annotations for SNPs in the binary annotation (Methods), meta-analyzed this quantity across traits, and compared observed vs. expected *λ*^2^(*C*) (Figure 3b and Table S20). We observed strong concordance, with a slope of 0.57 (correlation of 0.61) across the 20 binary annotations. This implies that the depletions of 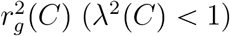 within binary annotations are largely explained by corresponding values of continuous-valued annotations.

In summary, our results show that causal disease effect sizes are more population-specific in functionally important regions impacted by selection. Further interpretation of these findings, including the role of positive and/or negative selection, is provided in the Discussion section.

### Analysis of specifically expressed gene annotations

We analyzed 53 specifically expressed gene (SEG) annotations, defined in ref.^24^ as ±100kb regions surrounding the top 10% of genes specifically expressed in each of 53 GTEx^32^ tissues (Table S2), by applying S-LDXR with the baseline-LD-X model to the 31 diseases and complex traits (Table S16). We note that although SEG annotations were previously used to prioritize disease-relevant tissues based on disease-specific heritability enrichments,^20,24^ enrichment/depletion of squared trans-ethnic genetic correlation (*λ*^2^(*C*)) is standardized with respect to heritability (i.e. increase in heritability in the denominator would lead to increase in trans-ethnic genetic covariance in the numerator (Equation (2))), hence not expected to produce extremely disease-specific signals. Thus, we first assess meta-analyzed *λ*^2^(*C*) estimates across the 31 diseases and complex traits (trait-specific estimates are assessed below).

Results are reported in Figure 4a and Table S21. *λ*^2^(*C*) estimates were less than 1 for all 53 tissues and significantly less than 1 (*p* < 0.05/53) for 37 tissues, with statistically significant heterogeneity across tissues (*p* < 10^−20^; Methods). The strongest depletions of squared trans-ethnic genetic correlation were observed in skin tissues (e.g. *λ*^2^(*C*) = 0.83 (s.e. 0.02) for Skin Sun Exposed (Lower Leg)), Prostate and Ovary (*λ*^2^(*C*) = 0.84 (s.e. 0.02) for Prostate, *λ*^2^(*C*) = 0.86 (s.e. 0.02) for Ovary) and immune-related tissues (e.g. *λ*^2^(*C*) = 0.85 (s.e. 0.02) for Spleen), and the weakest depletions were observed in Testis (*λ*^2^(*C*) = 0.98 (s.e. 0.02); no significant depletion) and brain tissues (e.g. *λ*^2^(*C*) = 0.98 (s.e. 0.02) for Brain Nucleus Accumbens (Basal Ganglia); no significant depletion). Results were similar at less stringent and more stringent shrinkage parameter values (*α* = 0.0 and *α* = 1.0; Figures S23, S24 and Table S21). A comparison of 14 blood-related traits and 16 other traits yielded highly consistent *λ*^2^(*C*) estimates (*R* = 0.82; Figure S25, Table S22), confirming that these findings were not extremely disease-specific.

**Figure 4:**
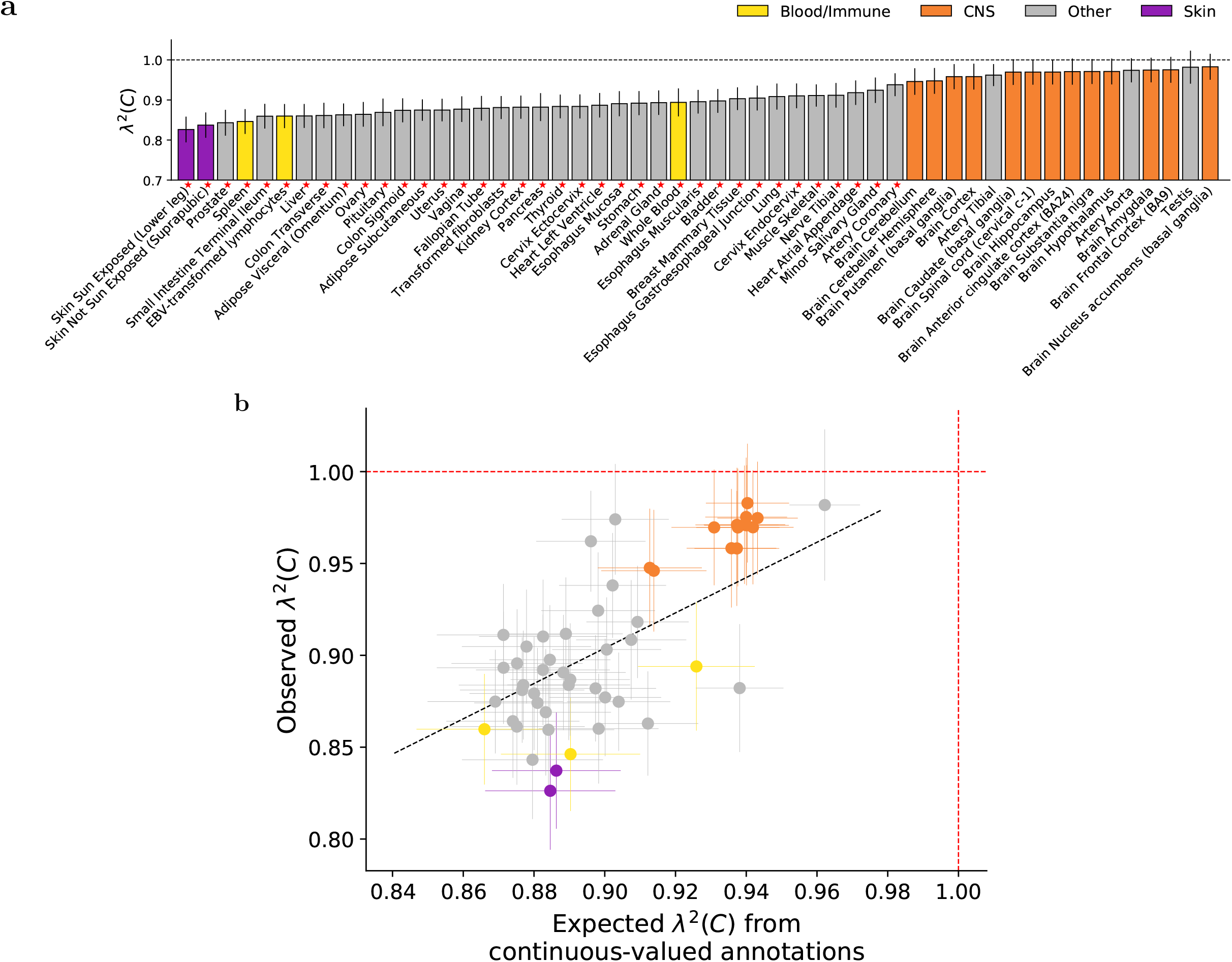
S-LDXR results for 53 specifically expressed gene (SEG) annotations across 31 diseases and complex traits. (a) We report estimates of the enrichment/depletion of squared trans-ethnic genetic correlation (*λ*^2^(*C*)) for each SEG annotation (sorted by *λ*^2^(*C*)). Results are meta-analyzed across 31 diseases and complex traits. Error bars denote±1.96× standard error. Red stars (⋆) denote two-tailed p<0.05/53. Numerical results are reported in Table S21. (b) We report observed *λ*^2^ *C* vs. expected *λ*^2^(*C*) based on 8 continuous-valued annotations, for each SEG annotation. Results are meta-analyzed across 31 diseases and complex traits. Error bars denote ±1.96× standard error. Annotations are color-coded as in (a). The dashed black line (slope=0.96) denotes a regression of observed *λ*(*C*) - 1 vs. expected *λ*(*C*) - 1 with intercept constrained to 0. Numerical results and population-specific heritability enrichment estimates are reported in Table S23.

These *λ*^2^(*C*) results were consistent with the higher background selection statistic^30^ in Skin Sun Exposed (Lower Leg) (*R* = 0.17), Prostate (*R* = 0.16) and Spleen (*R* = 0.14) as compared to Testis (*R* = 0.02) and Brain Nucleus Accumbens (Basal Ganglia) (*R* = 0.08) (Figure S26, Table S2), and similarly for CpG content (Figure S27, Table S2). Although these results could in principle be confounded by gene size,^33^ the low correlation between gene size and background selection statistic (*R* = 0.06) or CpG content (*R* = −0.20) (in ±100kb regions) implies limited confounding. We note the well-documented action of recent positive selection on genes impacting skin pigmentation,^34–38^ the immune system,^34–37,39^ and Ovary;^40^ we are not currently aware of any evidence of positive selection impacting Prostate. We further note the well-documented action of negative selection on fecundity- and brain-related traits,^26,28,41^ but it is possible that recent positive selection may more closely track differences in causal disease effect sizes across human populations, which have split relatively recently^42^ (see Discussion).

More generally, since SEG annotations are moderately correlated with the 8 continuous-valued annotations (Figure S28, Table S2), we investigated whether these *λ*^2^(*C*) results could be explained by the 8 continuous-valued annotations (analogous to Figure 3b). Results are reported in Figure 4b and Table S23. We observed strong concordance, with a slope of 0.96 (correlation of 0.76) across the 53 SEG annotations. This implies that the depletions of *λ*^2^(*C*) within SEG annotations are explained by corresponding values of continuous-valued annotations.

The strong depletion of squared trans-ethnic genetic correlation in tissues impacted by positive selection (as opposed to negative selection) suggests a possible connection between positive selection and population-specific causal effect sizes. To further assess this, we estimated the enrichment/depletion of squared trans-ethnic genetic correlation in SNPs with high integrated haplotype score (iHS),^43,44^ which quantifies the action of positive selection (Methods). We observed a significant depletion (*λ*^2^(*C*) = 0.88 (s.e. 0.03)), further implicating positive selection (however, it is difficult to assess whether the iHS annotation contains unique information about *λ*^2^(*C*) conditional on other annotations; see Discussion). In addition, we observed a high genome-wide trans-ethnic genetic correlation for schizophrenia (*r*_*g*_ = 0.95 (s.e. 0.04) vs. average of 0.85 (s.e. 0.01) across traits), a psychiatric disorder hypothesized to be strongly impacted by negative selection,^45,46^ suggesting that negative selection may play a limited role in population-specific causal effect sizes. As noted above, these estimates pertain to parameters that were defined based on common variants (see Overview of methods); we note that although negative selection has the strongest impact on low-frequency variants,^26^ common variants are also impacted by negative selection and can inform inferences about negative selection.^22^ The role of positive selection (as opposed to negative selection) in population-specific causal effect sizes is discussed further in the Discussion section.

We investigated the enrichment/depletion of *λ*^2^(*C*) in the 53 specifically expressed gene annotations for each individual trait (Table S24). We identified 6 significantly depleted (vs. 0 significantly enriched) trait-tissue pairs at per-trait *p* < 0.05/53. The limited number of statistically significant results was expected, due to the reduced power of trait-specific analyses; however, *λ*^2^(*C*) estimates were generally consistent across traits. Results for BMI and height, two widely studied anthropometric traits, are reported in Figure 5. For BMI, we observed significant depletion of squared trans-ethnic genetic correlation (*λ*^2^(*C*) = 0.84 (s.e. 0.05)) in Pituitary. Previous studies have highlighted the role of Pituitary in obesity;^47–49^ our results suggest that this tissue-specific mechanism is population-specific. For height, we observed significant depletion of squared trans-ethnic genetic correlation for Transformed fibroblasts (*λ*^2^(*C*) = 0.87 (s.e. 0.03)), a connective tissue linked to human developmental disorders;^50^ again, our results suggest that this tissue-specific mechanism is population-specific. Although Pituitary was significantly depleted for BMI but not height, and Transformed fibroblasts was significantly depleted for height but not BMI, we caution that for both tissues our *λ*^2^(*C*) estimates did not differ significantly between BMI and height.

In summary, our results show that causal disease effect sizes are more population-specific in regions surrounding specifically expressed genes.

**Figure 5:**
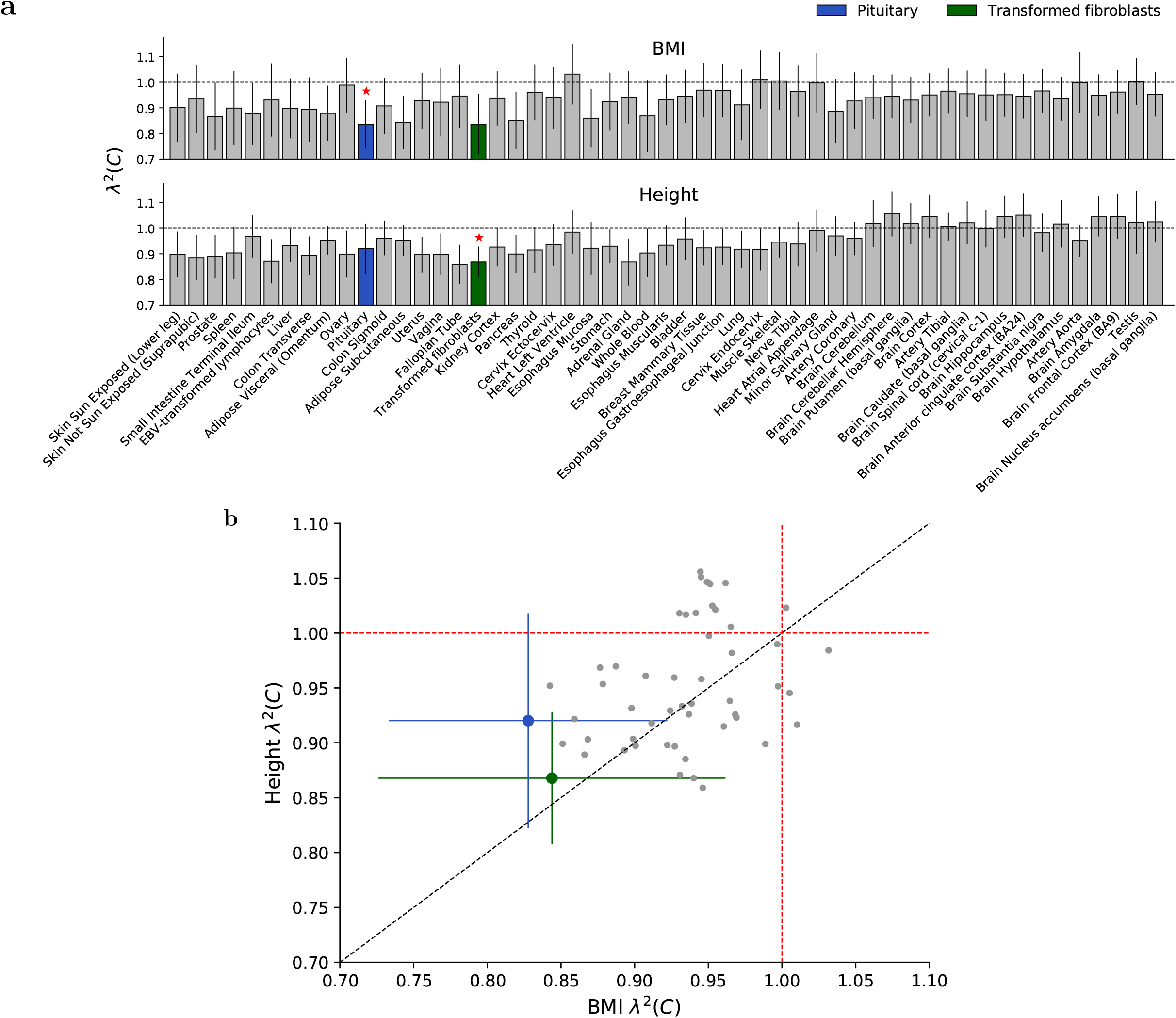
S-LDXR results for 53 specifically expressed gene (SEG) annotations for BMI and height. (a) We report estimates of the enrichment/depletion of squared trans-ethnic genetic correlation (*λ*^2^ *C*) for each SEG annotation for BMI and height. SEG annotations are ordered as in Figure 4. Error bars denote ±1.96× standard error. Red stars (⋆) denote two-tailed *p* < 0.05/53 for each respective trait. (b) We report *λ*^2^ *C* estimates for height vs. BMI for each SEG annotation. These estimates were moderately correlated (*R* = 0.35). Annotations are color-coded as in (a). For Pituitary, height *λ*^2^(*C*) is left-shifted by 0.008 and BMI *λ*^2^ *C* is right-shifted by 0.008 for easier visualization of standard errors. Error bars denote ±1.96× standard error. The dashed black line denotes the y vs x line. Numerical results for all 31 diseases and complex traits are reported in Table S24.

## Discussion

We developed a new method (S-LDXR) for stratifying squared trans-ethnic genetic correlation across functional categories of SNPs that yields approximately unbiased estimates in extensive simulations. By applying S-LDXR to East Asian and European summary statistics across 31 diseases and complex traits, we determined that SNPs with high background selection statistic^30^ have substantially depleted squared trans-ethnic genetic correlation (vs. the genome-wide average), implying that causal effect sizes are more population-specific. Accordingly, squared trans-ethnic genetic correlations were substantially depleted for SNPs in many functional categories and enriched in less functionally important regions (although the power of S-LDXR to detect enrichment of squared trans-ethnic genetic correlation is limited due to depletion of heritability in less functionally important regions). In analyses of specifically expressed gene annotations, we observed substantial depletion of squared trans-ethnic genetic correlation for SNPs near skin and immune-related genes, which are strongly impacted by recent positive selection, but not for SNPs near brain genes. We also observed trait-specific depletions of squared-trans-ethnic genetic correlation for specifically expressed gene annotations, which indicate population-specific disease mechanisms.

Reductions in trans-ethnic genetic correlation have several possible underlying explanations, including gene-environment (G×E) interaction, gene-gene (G×G) interaction, and dominance variation (but not differences in heritability across populations, which would not affect trans-ethnic genetic correlation and were not observed in our study). Given the increasing evidence of the role of G×E interaction in complex trait architectures,^51^ and evidence that G×G interaction and dominance variation explain limited heritability,^52–54^ we hypothesize that depletion of squared trans-ethnic genetic correlation in the top quintile of background selection statistic and in functionally important regions may be primarily attributable to stronger GxE interaction in these regions. Interestingly, a recent study on plasticity in Arabidopsis observed a similar phenomenon: lines with more extreme phenotypes exhibited stronger G×E interaction.^55^ Although depletion of squared trans-ethnic genetic correlation is often observed in regions with higher per-SNP heritability, which may often be subject to stronger GxE, depletion may also occur in regions with lower per-SNP heritability that are subject to stronger GxE; we hypothesize that this is the case for SNPs in the top quintile of average LLD and the bottom quintile of GERP (NS) (Figure 2).

Distinguishing between stronger GxE interaction in regions impacted by selection and stronger G×E interaction in functionally important regions as possible explanations for our findings is a challenge, because functionally important regions are more strongly impacted by selection. To this end, we constructed an annotation that is similar to the background selection statistic but does not make use of recombination rate, instead relying solely on a SNP’s physical distance to the nearest exon (Methods). Applying S-LDXR to the 31 diseases and complex traits using a joint model incorporating baseline-LD-X model annotations and the nearest exon annotation, the background selection statistic remained highly conditionally informative for trans-ethnic genetic correlation, whereas the nearest exon annotation was not conditionally informative (Table S25). This result implicates stronger G×E interaction in regions with reduced effective population size that are impacted by selection, and not just proximity to functional regions, in explaining depletions of squared trans-ethnic genetic correlation; however, we emphasize that selection acts on allele frequencies rather than causal effect sizes, and could help explain our findings only in conjunction with other explanations such as G×E interaction. Our results on specifically expressed genes implicate stronger GxE interaction near skin, immune, and ovary genes and weaker GxE interaction near brain genes, potentially implicating positive selection (as opposed to negative selection). This conclusion is further supported by the significant depletion of squared trans-ethnic genetic correlation in the integrated haplotype score (iHS) annotation that specifically reflects positive selection, high genome-wide trans-ethnic genetic correlation for schizophrenia (Table S16), and lack of variation in squared trans-ethnic genetic correlation across genes in different deciles of probability of loss-of-function intolerance^56^ (Methods, Figure S29, S30, Table S26). We conclude that depletions of squared trans-ethnic genetic correlation could potentially be explained by stronger GxE interaction at loci impacted by positive selection. We caution that other explanations are also possible; in particular, evolutionary modeling using an extension of the Eyre-Walker model^57^ to two populations suggests that our results for the background selection statistic could also be consistent with negative selection (Supplementary Note, Figure S31, S32, Table S27). Additional information, such as genomic annotations that better distinguish different types of selection or data from additional diverse populations, may help elucidate the relationship between selection and population-specific causal effect sizes.

Our study has several implications. First, polygenic risk scores (PRS) in non-European populations that make use of European training data^6,9,11^ may be improved by reweighting SNPs based on the expected enrichment/depletion of squared trans-ethnic genetic correlation, helping to alleviate health disparities.^6,14,15^ For example, when applying LD-pruning + p-value thresholding methods,^58,59^ both the strength of association and trans-ethnic genetic correlation should be accounted for when prioritizing SNPs for trans-ethnic PRS, as our results suggest that trans-ethnic genetic correlation is likely depleted near functional SNPs with significant p-values (due to stronger G×E). In particular, when multiple SNPs have similar level of significance, the SNPs enriched for trans-ethnic genetic correlation should be prioritized. Analogously, when applying more recent methods that estimate posterior mean causal effect sizes^60–66^ (including functionally informed methods^62,66^), these estimates should subsequently be weighted according to the expected enrichment/depletion for squared trans-ethnic genetic correlation based on their functional annotations. Second, modeling population-specific genetic architectures may improve trans-ethnic fine-mapping. Our results suggest that causal effect sizes and/or causal variants are likely to differ across different populations, contrary to standard assumptions.^31,67^ Thus, incorporating information about trans-ethnic genetic correlations in trans-ethnic fine-mapping may lead to more accurate identification of both population-specific and shared causal variants.^68^ Third, modeling population-specific genetic architectures may also increase power in trans-ethnic meta-analysis,^69^ e.g. by adapting MTAG^70^ to two populations (instead of two traits), leveraging trans-ethnic (instead of cross-trait) genetic correlation between pairs of populations to improve estimation of SNP effect sizes in both populations. Fourth, it may be of interest to stratify GxE interaction effects^51^ across genomic annotations. Fifth, modeling and incorporating environmental variables, where available, may provide additional insights into population-specific causal effect sizes. In our simulations, we did not explicitly simulate GxE. However, GxE would induce population-specific causal effect sizes, which we did explicitly simulate. Sixth, the S-LDXR method could potentially be extended to stratify squared *cross-trait* genetic correlations^71^ across genomic annotations.^72^

We note several limitations of this study that pertain to the S-LDXR method. First, S-LDXR is designed for populations of homogeneous continental ancestry (e.g. East Asians and Europeans) and is not currently suitable for analysis of admixed populations^73^ (e.g. African Americans or admixed Africans from UK Biobank^74^), analogous to LDSC and its published extensions.^21,71,75^ However, a recently proposed extension of LDSC to admixed populations^76^ could be incorporated into S-LDXR, enabling its application to the growing set of large studies in admixed populations.^10^ Second, S-LDXR estimates of enrichment of stratified squared trans-ethnic genetic correlation (*λ*^2^(*C*)) are slightly downward biased in null simulations of the top quintile of the background selection statistic and average LLD annotations, especially in simulations involving annotation-dependent MAF-dependent genetic architectures. However, these biases are small compared to the depletions of *λ*^2^(*C*) observed in analysis of real traits. We further note that our estimates are unbiased in null simulations of binary annotations, implying that our results on real traits for binary annotations are extremely robust. Third, since S-LDXR applies shrinkage to reduce standard error in estimating stratified squared trans-ethnic genetic correlation and its enrichment, estimates are conservative – true depletions of squared trans-ethnic genetic correlation in functionally important regions may be stronger than the estimated depletions. However, we emphasize that S-LDXR is approximately unbiased in null data. Fourth, the optimal value of the shrinkage parameter *α* may be specific to the pair of populations analyzed. In our simulations, we determined that *α* = 0.5 provides a satisfactory bias-variance tradeoff across a wide range of values of polygenicity and power. Thus, *α* = 0.5 may also be satisfactory for other pairs of populations. However, we recommend that one should ideally perform simulations on the pair of populations being analyzed to selection the optimal value of *α*. Fifth, it is difficult to assess whether a focal annotation contains unique information about *λ*^2^(*C*) conditional on other annotations, as squared trans-ethnic genetic correlation is a non-linear quantity defined by the quotient of squared trans-ethnic genetic covariance and the product of heritabilities in each population.

We also note several limitations of this study that pertain to our analysis of real traits. First, we focused on comparisons of East Asians and Europeans, due to limited availability of very large GWAS in other populations. For other pairs of continental populations, if differences in environment are similar, then we would expect similar genome-wide trans-ethnic genetic correlation and similar enrichment/depletion of squared trans-ethnic genetic correlation, based on our hypothesis that imperfect trans-ethnic genetic correlation is primarily attributable to G×E. We also note that different set of SNPs, with different MAF and LD patterns, would be analyzed for different pairs of populations. However, we expect that these differences would not contribute to differences in trans-ethnic genetic correlation, if G×E is the fundamental factor impacting trans-ethnic genetic correlation. Second, the specifically expressed gene (SEG) annotations analyzed in this study are defined predominantly (but not exclusively) based on gene expression measurements of Europeans.^24^ We hypothesize that results based on SEG annotations defined in East Asian populations would likely be similar, as heritability enrichment of functional annotations (predominantly defined in Europeans) are consistent across continental populations,^20,31^ despite the fact that gene expression patterns and genetic architectures of gene expression differ across diverse populations.^12,77,78^ Thus, SEG annotations derived from gene expression data from diverse populations may provide additional insights into population-specific causal effect sizes. Third, we restricted our analyses to SNPs that were relatively common (MAF>5%) in both populations (estimating parameters that were defined based on common SNPs), due to the lack of a large LD reference panel for East Asians. Extending our analyses to lower-frequency SNPs may provide further insights into the role of negative selection in shaping population-specific genetic architectures, as negative selection has the strongest impact on variants with low frequency.^26,27^ Fourth, we did not consider population-specific variants in our analyses, due to the difficulty in defining trans-ethnic genetic correlation for population-specific variants^2,5^, a more fundamental challenge than analyzing low-frequency SNPs; a recent study^79^ has reported that population-specific variants substantially limit trans-ethnic genetic risk prediction accuracy. Fifth, estimates of genome-wide trans-ethnic genetic correlation may be confounded by different trait definitions or diagnostic criteria in the two populations, particularly for major depressive disorder. However, this would not impact estimates of enrichment/depletion of squared trans-ethnic genetic correlation (*λ*^2^(*C*)), which is defined relative to genome-wide values. Sixth, we have not pinpointed the exact underlying phenomena (e.g. environmental heterogeneity coupled with gene-environment interaction) that lead to population-specific causal disease effect sizes at functionally important regions. Despite these limitations, our study provides an improved understanding of the underlying biology that contribute to population-specific causal effect sizes, and highlights the need for increasing diversity in genetic studies.

## URLs

- S-LDXR software: https://github.com/huwenboshi/s-ldxr/
- Python code for simulating GWAS summary statistics: https://github.com/huwenboshi/s-ldxr-sim/
- baseline-LD-X model annotations and LD scores: https://data.broadinstitute.org/alkesgroup/S-LDXR/
- Distance to nearest exon annotation and LD scores: https://data.broadinstitute.org/alkesgroup/S-LDXR/
- baseline-LD model annotations: https://alkesgroup.broadinstitute.org/LDSCORE/
- 1000 Genomes Project: https://www.internationalgenome.org/
- PLINK2: https://www.cog-genomics.org/plink/2.0/
- HAPGEN2: https://mathgen.stats.ox.ac.uk/genetics_software/hapgen/hapgen2.html
- UCSC Genome Browser: https://genome.ucsc.edu/
- Exome Aggregation Consortium (ExAC): https://exac.broadinstitute.org/
- Integrated haplotype scores (iHS): http://coruscant.itmat.upenn.edu/data/JohnsonEA_iHSscores.tar.gz

## Methods

### Definition of stratified squared trans-ethnic genetic correlation

We model a complex phenotype in two populations using linear models, ****Y****_1_ = ***X***_1_***β***_1_ +***ϵ***_1_ and ***Y***_2_ = ***X***_2_***β***_2_ +***ϵ***_2_, where ***Y***_1_ and ***Y***_2_ are vectors of phenotype measurements of population 1 and population 2 with sample size *N*_1_ and *N*_2_, respectively; ***X***_1_ and ***X***_2_ are mean-centered *but not normalized* genotype matrices at *M* SNPs in the two populations; ***β***_1_ and ***β***_2_ are *per-allele causal* effect sizes of the *M* SNPs; and ***ϵ***_1_ and ***ϵ***_2_ are environmental effects in the two populations. We assume that in each population, genotypes, causal effect sizes, and environmental effects are independent from each other. We assume that the per-allele effect size of SNP *j* in the two populations has variance and covariance,

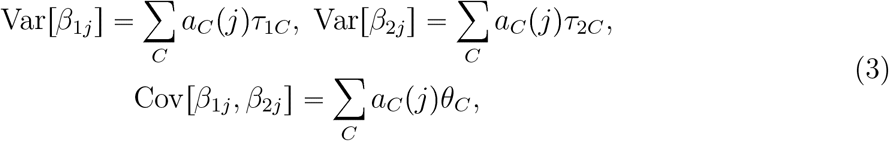

where *a*_*C*_(*j*) is the value of SNP *j* for annotation *C*, which can be binary or continuous-valued; *τ*_1*C*_ and *τ*_2*C*_ are the net contribution of annotation *C* to the variance of *β*_1*j*_ and *β*_2*j*_, respectively; and *θ*_*C*_ is the net contribution of annotation *C* to the covariance of *β*_1*j*_ and *β*_2*j*_.

We define stratified trans-ethnic genetic correlation of a binary annotation *C* (e.g. functional annotations^21^ or quintiles of continuous-valued annotations^22^) as,

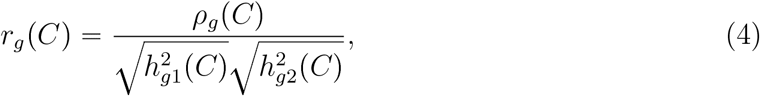

where *ρ_g_*(*C*) = Σ_*j*∈*C*_ Cov[*β*_1*j*_, *β*_2*j*_] = Σ_*j*∈*C*_ Σ_*C′*_ *a*_*C′*_ (*j*)*θ*_*C′*_ is the trans-ethnic genetic covariance of annotation *C*; and 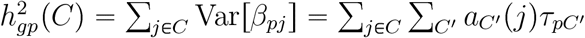 is the “allelic-scale heritability” (sum of per-SNP variance of per-allele causal effect sizes; different from heritability on the standardized scale) of annotation *C* in population *p*. Here, *C′* includes all binary and continuous-valued annotations included in the analysis. Since estimates of 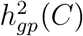 can be noisy (possibly negative), we estimate *squared* stratified trans-ethnic genetic correlation,

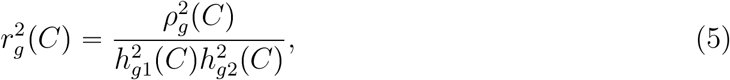

to avoid bias or undefined values in the square root. In this work, we only estimate 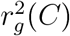 for SNPs with minor allele frequency (MAF) greater than 5% in both populations. To assess whether causal effect sizes are more or less correlated for SNPs in annotation *C* compared with the genome-wide average 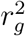, we define the enrichment/depletion of stratified squared trans-ethnic genetic correlation as

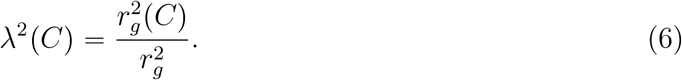

We meta-analyze *λ*^2^(*C*) instead of 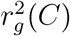 across diseases and complex traits. For continuous-valued annotations, defining 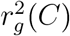 and *λ*^2^(*C*) is challenging, as squared correlation is a non-linear term involving a quotient of squared covariance and a product of variances; we elected to instead estimate *λ*^2^(*C*) for quintiles of continuous-valued annotations (analogous to ref.^22^). We note that the average value of *λ*^2^(*C*) across quintiles of continuous-valued annotations is not necessarily equal to 1, as squared trans-ethnic genetic correlation is a non-linear quantity.

### S-LDXR method

S-LDXR is conceptually related to stratified LD score regression^21,22^ (S-LDSC), a method for stratifying heritability from GWAS summary statistics, to two populations. The S-LDSC method determines that a category of SNPs is enriched for heritability if SNPs with high LD to that category have higher expected *χ*^2^ statistic than SNPs with low LD to that category. Analogously, the S-LDXR method determines that a category of SNPs is enriched for trans-ethnic genetic covariance if SNPs with high LD to that category have higher expected product of Z-scores than SNPs with low LD to that category.

S-LDXR relies on the regression equation

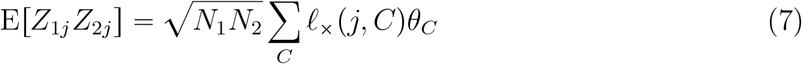

to estimate *θ*_*C*_, where *Z*_*pj*_ is the Z-score of SNP *j* in population *p*; *ℓ*×(*j, C*) = Σ_*k*_ *r*_1*jk*_*r*_2*jk*_σ_1*j*_σ_2*j*_*a*_*C*_(*k*) is the trans-ethnic LD score of SNP *j* with respect to annotation *C*, whose value for SNP *k*, *a_C_*(*k*), can be either binary or continuous; *r*_*pjk*_ is the LD between SNP *j* and *k* in population *p*; and *σ*_*pj*_ is the standard deviation of SNP *j* in population *p*. We obtain unbiased estimates of *ℓ*×(*j, C*) using genotype data of 481 East Asian and 489 European samples in the 1000 Genomes Project.^16^ To account for heteroscedasticity and increase statistical efficiency, we use weighted least square regression to estimate *θ*_*C*_. We use regression equations analogous to those described in ref.^21^ to estimate *τ*_1*C*_ and *τ*_2*C*_. We include only well-imputed (imputation INFO>0.9) and common (MAF>5% in both populations) SNPs that are present in HapMap 3^17^ (irrespective of GWAS significance level) in the regressions (*regression SNPs*), analogous to our previous work.^21,71,75^ We use all SNPs present in either population in 1000 Genomes^16^ to estimate trans-ethnic LD scores *ℓ*×(*j, C*) (*reference SNPs*; analogous to S-LDSC^21^), so that the resulting coefficients *θ*_*C*_ also pertain to these SNPs. However, we estimate 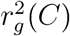 and *λ*^2^(*C*) (see below; defined as a function of causal effect sizes) for all SNPs with MAF>5% in both populations (*heritability SNPs*), accounting for tagging effects (analogous to S-LDSC^21^).

Let 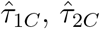, and 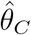 be the estimates of *τ*_1*C*_, *τ*_2*C*_, and *θ*_*C*_, respectively. For each binary annotation *C*, we estimate the stratified heritability of annotation *C* in each population, 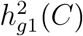 and 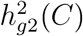, and trans-ethnic genetic covariance, *ρ_g_*(*C*), as

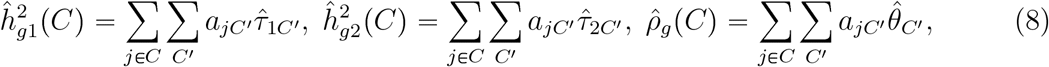

respectively, restricting to causal effects of SNPs with MAF>5% in both populations (*heritability SNPs*), using coefficients (*τ*_1*C′*_, *τ*_2*C′*_, and *θ*_*C′*_) of both binary and continuous-valued annotations. We estimate genome-wide trans-ethnic genetic correlation as 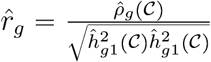, where 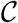 represents the set of all SNPs with MAF>5% in both populations. We then estimate 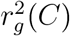 as

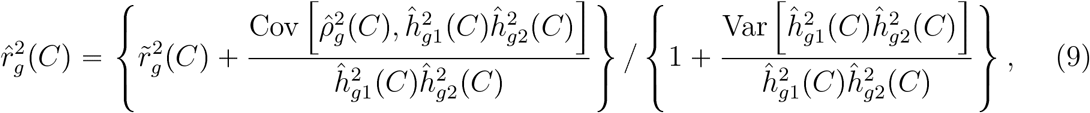

where 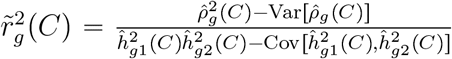. The correction to 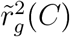 in Equation (9) is necessary for obtaining an unbiased estimate of 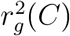, as computing quotients of two random variables introduces bias (Supplementary Note). (We do not constrain the estimate of 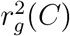 to its plausible range of [−1, 1], as this would introduce bias.) Subsequently, we estimate enrichment of stratified squared trans-ethnic genetic correlation as

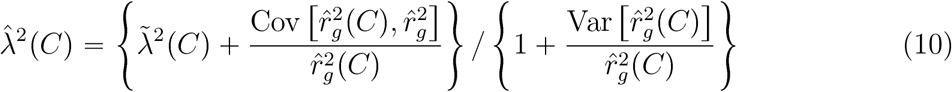

where 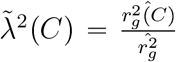, the ratio between estimated stratified 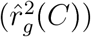 and genome-wide 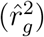 squared trans-ethnic genetic correlation. We use block jackknife over 200 non-overlapping and equally sized blocks to obtain standard error of all estimates. The standard error of *λ*^2^(*C*) primarily depends on the total allelic-scale heritability of SNPs in the annotation (sum of per-SNP variances of causal per-allele effect sizes), which appears as the denominator 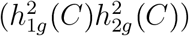 in the estimation of stratified squared trans-ethnic genetic correlation 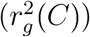; if this denominator is small, estimation of 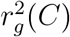 becomes noisy. The standard error of *λ*^2^(*C*) indirectly depends on the size of the annotation, because larger annotations tend to have larger total heritability. However, estimates of *λ*^2^(*C*) for a large annotation may have large standard error if the annotation is depleted for heritability.

To assess the informativeness of each annotation in explaining disease heritability and trans-ethnic genetic covariance, we define standardized annotation effect size on heritability and trans-ethnic genetic covariance for each annotation *C* analogous to ref.^22^,

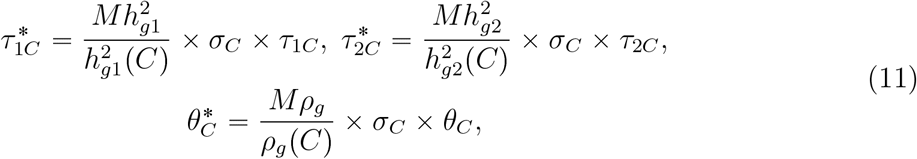

where 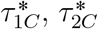, and 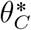 represent proportionate change in per-SNP heritability in population 1 and 2 and trans-ethnic genetic covariance, respectively, per standard deviation increase in annotation *C*; *τ*_1*C*_, *τ*_2*C*_, and *θ*_*C*_ are the corresponding unstandardized effect sizes, defined in Equation (3); and *σ_C_* is the standard deviation of annotation *C*.

We provide a more detailed description of the method, including derivations of the regression equation and unbiased estimators of the LD scores, in the **Supplementary Note**.

### S-LDXR shrinkage estimator

Estimates of 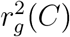 can be imprecise with large standard errors if the denominator, 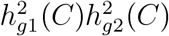, is close to zero and noisily estimated. This is especially the case for annotations of small size (< 1% SNPs). We introduce a shrinkage estimator to reduce the standard error in estimating 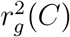.

Briefly, we shrink the estimated per-SNP heritability and trans-ethnic genetic covariance of annotation *C* towards the genome-wide averages, which are usually estimated with smaller standard errors, prior to estimating 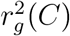. In detail, let *M_C_* be the number of SNPs in annotation *C*, we shrink 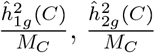, and 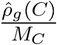, towards 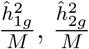 and 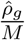, respectively, where 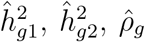 are the genome-wide estimates, and *M* the total number of SNPs. We obtain the shrinkage as follows. Let 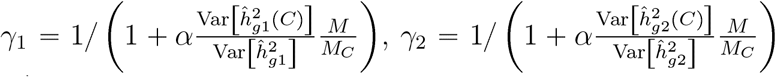, and 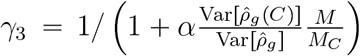 be the shrinkage obtained separately for 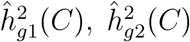 and 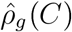, respectively, where *α* ∈ [0, 1] is the shrinkage parameter adjusting magnitude of shrinkage. We then choose the most stringent shrinkage, *γ* = min {*γ*_1_*, γ*_2_*, γ*_3_}, as the final shared shrinkage for both heritability and trans-ethnic genetic covariance.

We shrink heritability and trans-ethnic genetic covariance of annotation *C* using *γ* as,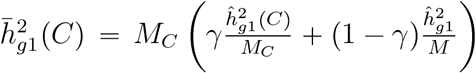,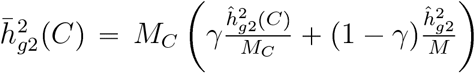 and 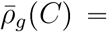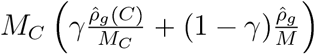, where 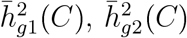, and 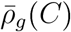 are the shrunk counterparts of 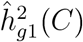, 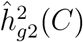, and 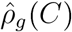, respectively. We shrink 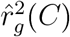 by substituting 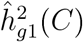, 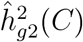, and 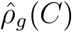 with 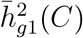, 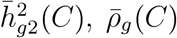, respectively, in Equation (9), to obtain its shrunk counterpart, 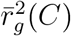. Finally, we shrink 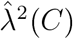, by plugging in 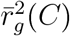 in Equation (10) to obtain its shrunk counterpart, 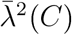. We recommend *α* = 0.5 as the default shrinkage parameter value, as this value provides robust estimates of *λ*^2^(*C*) in simulations. We note that S-LDXR does not use the shrinkage estimator when estimating genome-wide *r*_*g*_ and 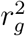.

### Significance testing

To assess whether an annotation *C* is enriched or depleted of squared trans-ethnic genetic correlation for a trait, we test the null hypothesis 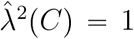. Since 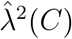 is not normally distributed,^80^ we instead test the equivalent null hypothesis 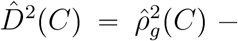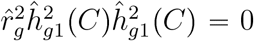, where 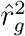 is the genome-wide squared trans-ethnic genetic correlation. We obtain test statistic as 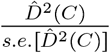, and obtain p-value under t-distribution with *B* - 1 degrees of freedom, where *B* is the number of jackknife blocks. Since the 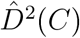 statistic does not involve division by 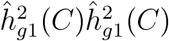, we do not apply any shrinkage to 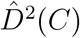.

### Baseline-LD-X model

We include a total of 54 binary functional annotations in the baseline-LD-X model. These include 53 annotations introduced in ref.,^21^ which consists of 28 main annotations including conserved annotations (e.g. Coding, Conserved) and epigenomic annotations (e.g. H3K27ac, DHS, Enhancer) derived from ENCODE^81^ and Roadmap,^82^ 24 500-base-pair-extended main annotations, and 1 annotation containing all SNPs. We note that although chromatin accessibility can be population-specific, the fraction of such regions is small.^83^ Following ref,^22^ we created an additional annotation for all genomic positions with number of rejected substitutions^84^ greater than 4. Further information for all functional annotations included in the baseline-LD-X model is provided in Table S1a.

We also include a total of 8 continuous-valued annotations in the baseline-LD-X model. First, we include 5 continuous-valued annotations introduced in ref.^22^ (see URLs), without modification: background selection statistic,^30^ CpG content (within a ±50 kb window), GERP (number of substitutation) score,^84^ nucleotide diversity (within a ±10 kb window), and Oxford map recombination rate (within a ±10 kb window).^85^ Second, we include 2 minor allele frequency (MAF) adjusted annotations introduced in ref.,^22^ with modification: level of LD (LLD) and predicted allele age. We created analogous annotations applicable to both East Asian and European populations. To create an analogous LLD annotation, we estimated LD scores for each population using LDSC,^75^ took the average across populations, and then quantile-normalized the average LD scores using 10 average MAF bins. We call this annotation “average level of LD”. To create analogous predicted allele age annotation, we quantile-normalized allele age estimated by ARGweaver^86^ across 54 multi-ethnic genomes using 10 average MAF bins. Finally, we include 1 continuous-valued annotation based on *F*_ST_ estimated by PLINK2,^87^ which implements the Weir & Cockerham estimator of *F*_ST_.^88^ Further information for all continuous-valued annotations included in the baseline-LD-X model is provided in Table S1b.

### Code and data availability

Python code implementing S-LDXR is available at https://github.com/huwenboshi/s-ldxr. Python code for simulating GWAS summary statistics under the baseline-LD-X model is available at https://github.com/huwenboshi/s-ldxr-sim. baseline-LD-X model annotations and LD scores are available at https://data.broadinstitute.org/alkesgroup/S-LDXR/.

### Simulations

We used simulated East Asian (EAS) and European (EUR) genotype data to assess the performance our method, as we did not have access to real EAS genotype data at sufficient sample size to perform simulations with real genotypes. We simulated genotype data for 100,000 East-Asian-like and 100,000 European-like individuals using HAPGEN2^25^ (see URLs), which preserves population-specific MAF and LD patterns, starting from phased haplotypes of 481 East Asians and 489 Europeans individuals available in the 1000 Genomes Project^16^ (see URLs), restricting to −2.5 million SNPs on chromosome 1 – 3 with minor allele count greater than 5 in either population. Since direct output of HAPGEN2 includes substantial relatedness,^2^ we used PLINK2^87^ (see URLs) to remove simulated individuals with genetic relatedness greater than 0.05, resulting in 35,378 EAS-like and 36,836 EUR-like individuals. From the filtered set of individuals, we randomly selected 500 individuals in each simulated population to serve as reference panels. We used 18,418 EAS-like and 36,836 EUR-like individuals to simulate GWAS summary statistics, capturing the imbalance in sample size between EAS and EUR GWAS in analysis of real traits. In our secondary simulations, we also decreased or increased the reference panel size or decreased the GWAS sample size, to evaluate the robustness of our method with respect to reference panel size and GWAS sample size.

We performed both null simulations, where enrichment of squared trans-ethnic genetic correlation, *λ*^2^(*C*), is 1 across all functional annotations, and causal simulations, where *λ*^2^(*C*) varies across annotations, under various degrees of polygenicity (1%, 10%, and 100% causal SNPs). In the null simulations, we set *τ*_1*C*_, *τ*_2*C*_, *θ*_*C*_ to be the meta-analyzed *τ_C_* in real-data analyses of EAS GWASs, and followed Equation (3) to obtain variance, Var [*β*_1*j*_] and Var [*β*_2*j*_], and covariance, Cov [*β*_1*j*_, β_2*j*_], of per-SNP causal effect sizes *β*_1*j*_, *β*_2*j*_, setting all negative per-SNP variance and covariance to 0. In the causal simulations, we directly specified per-SNP causal effect size variances and covariances using self-devised *τ*_1*C*_, *τ*_2*C*_, and *θ*_*C*_ coefficients, to attain *λ*^2^(*C*) ≠ 1, as these were difficult to attain using the coefficients from analyses of real traits.

We randomly selected a subset of SNPs to be causal for both populations, and set Var [*β*_1*j*_], Var [*β*_2*j*_], and Cov [*β*_1*j*_, *β*_2*j*_] to be 0 for all remaining non-causal SNPs. We scaled the trans-ethnic genetic covariance to attain a desired genome-wide *r*_*g*_. Next, we drew causal effect sizes of each causal SNP *j* in the two populations from the bi-variate Gaussian distribution,

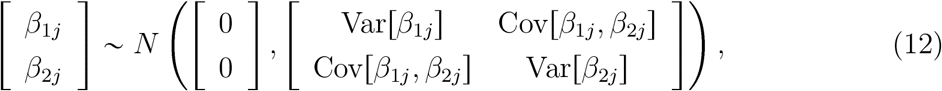

and scaled the drawn effect sizes to match the desired total heritability and trans-ethnic genetic covariance. We also performed null simulations in which imperfect genome-wide trans-ethnic genetic correlation is due to population-specific causal variants. In these simulations, we randomly selected 10% of the SNPs to be causal in each population, with 80% of causal variants in each population shared with the other population, and sampled perfectly correlated causal effect sizes for shared causal variants using Equation (12). We simulated genetic component of the phenotype in population *p* as ***X***_*p*_***β***_*p*_, where ***X***_***p***_ is column-centered genotype matrix, and drew environmental effects, ***ϵ***_***p***_, from the Gaussian distribution, *N* (0, 1 - Var [***X***_*p*_***β***_*p*_]), such that the total phenotypic variance in each population is 1. Finally, we simulated GWAS summary association statistics for population *p*, ***Z***_***p***_, as 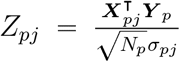, where *σ*_*pj*_ is the standard deviation of SNP *j* in population *p*. We have publicly released Python code for simulating GWAS summary statistics for 2 populations (see URLs). Fifth, we performed additional null simulations with annotation-dependent MAF-dependent genetic architectures,^26–28^ defined as architectures in which the level of MAF-dependence is annotation-dependent.

We also performed null simulations with annotation-dependent MAF-dependent genetic architectures,^26–28^ defined as architectures in which the level of MAF-dependence is annotation-dependent, to assess the impact on estimates of enrichment of stratified squared trans-ethnic genetic correlation, (*λ*^2^(*C*)). In these simulations, we set the variance of causal effect size of each SNP *j* in both populations to be proportional to [*p*_*j*,*max*_(1 - *p*_*j*,*max*_)]^*α*^, where *p*_*j*,*max*_ is the maximum MAF of SNP *j* in the two populations. (We elected to use maximum MAF because a SNP that is rare in one population but common in the other is less likely to be impacted by negative selection.) We set *α* to −0.38, as previously estimated for 25 UK Biobank diseases and complex traits in ref.^28^. We sampled causal effect sizes using Equation (12), with Var [*β*_1*j*_], Var [*β*_2*j*_], and Cov [*β*_1*j*_, β_2*j*_] scaled to attain a desired genome-wide heritability and trans-ethnic genetic correlation. We randomly selected 10% of SNPs to be causal in both populations. Additionally, in the top quintile of background selection statistic, we selected 1.8x more low-frequency causal variants (*p*_*j*,*max*_ < 0.05) than common variants (*p*_*j*,*max*_ ≥ 0.05), capturing the action of negative selection across low-frequency and common variants.^27^

### Summary statistics for 31 diseases and complex traits

We analyzed GWAS summary statistics of 31 diseases and complex traits, primarily from UK Biobank,^74^ Biobank Japan,^20^ and CONVERGE.^18^ All summary statistics were based on genotyping arrays with imputation to an appropriate LD reference panel (e.g. Haplotype Reference Consortium^89^ and UK10K^90^ for UK Biobank,^74^ the 1000 Genomes Project^16^ for Biobank Japan^20^), except those of the MDD GWAS in the East Asian population, which was based on low-coverage whole genome sequencing data.^18^ These include: atrial fibrillation (AF),^91,92^ age at menarche(AMN),^93,94^ age at menopause (AMP),^93,94^ basophil count(BASO),^20,95^ body mass index (BMI),^20,96^ blood sugar(BS),^20,96^ diastolic blood pressure (DBP),^20,96^ eosinophil count(EO),^20,96^ estimated glomerular filtration rate (EGFR),^20,97^ hemoglobin A1c(HBA1C),^20,96^ height (HEIGHT),^96,98^ high density lipoprotein (HDL),^20,96^ hemoglobin (HGB),^20,95^ hematocrit (HTC),^20,95^ low density lipoprotein (LDL),^20,96^ lymphocyte count(LYMPH),^20,96^ mean corpuscular hemoglobin (MCH),^20,96^ mean corpuscular hemoglobin concentration (MCHC),^20,95^ mean corpuscular volume (MCV),^20,95^ major depressive disorder (MDD),^18,99^ monocyte count (MONO),^20,96^ neutrophil count(NEUT),^20,95^ platelet count (PLT),^20,96^ rheumatoid arthritis(RA),^100^ red blood cell count (RBC),^20,96^ systolic blood pressure (SBP),^20,96^ schizophrenia (SCZ)^101^, type 2 diabetes (T2D),^102,103^ total cholesterol (TC),^20,96^ triglyceride (TG),^20,96^ and white blood cell count (WBC).^20,96^ Further information for the GWAS summary statistics analyzed is provided in Table S16. In our main analyses, we performed random-effect meta-analysis to aggregate results across all 31 diseases and complex traits. To test if the meta-analyzed 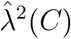 is significantly different from 1, we computed a test statistic as 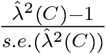, where *s.e.* 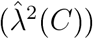 is the standard error of meta-analyzed 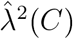, and obtained a p-value under the normal distribution. We also defined a set of 20 approximately independent diseases and complex traits with cross-trait 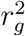 (estimated using cross-trait LDSC^71^) less than 0.25 in both populations: AF, AMN, AMP, BASO, BMI, EGFR, EO, HBA1C, HEIGHT, HTC, LYMPH, MCHC, MCV, MDD, NEUT, PLT, RA, SBP, TC, TG.

### Expected enrichment of stratified squared trans-ethnic genetic correlation from 8 continuous-valued annotations

To obtain expected enrichment of squared trans-ethnic genetic correlation of a binary annotation *C*, *λ*^2^(*C*), from 8 continuous-valued annotations, we first fit the S-LDXR model using these 8 annotations together with the base annotation for all SNPs, yielding coefficients, *τ*_1*C′*_, *τ*_2*C′*_, and *θ*_*C′*_, for a total of 9 annotations. We then use Equation (3) to obtain per-SNP variance and covariance of causal effect sizes, *β*_1*j*_ and *β*_1*j*_, substituting *τ*_1*C*_, *τ*_2*C*_, *θ*_*C*_ with *τ*_1*C′*_, *τ*_2*C′*_, and *θ*_*C′*_, respectively. We apply shrinkage with default parameter setting (*α* = 0.5), and use Equation (9) and (10) to obtain expected stratified squared trans-ethnic genetic correlation, 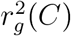, and subsequently *λ*^2^(*C*).

### Analysis of specifically expressed gene annotations

We obtained 53 specifically expressed gene (SEG) annotations, defined in ref.^24^ as ±100k-base-pair regions surrounding genes specifically expressed in each of 53 GTEx^32^ tissues. A list of the SEG annotations is provided in Table S2. Correlations between SEG annotations and the 8 continuous-valued annotations are reported in Figure S28 and Table S2. Most SEG annotations are moderately correlated with the background selection statistic and CpG content annotations.

To test whether there is heterogeneity in enrichment of squared trans-ethnic genetic correlation, *λ*^2^(*C*), across the 53 SEG annotations, we first computed the average *λ*^2^(*C*) across the 53 annotations, 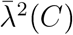, using fixed-effect meta-analysis. We then computed the test statistic 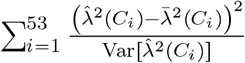, where *C*_*i*_ is the *i*-th SEG annotation, and 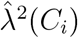 the estimated *λ*^2^(*C*). We computed a p-value for this test statistic based on a *χ*^2^ distribution with 53 degrees of freedom.

### Analysis of distance to nearest exon annotation

We created a continuous-valued annotation, named “distance to nearest exon annotation”, based on a SNP’s physical distance (number of base pairs) to its nearest exon, using 233,254 exons defined on the UCSC genome browser^104^ (see URLs). This annotation is moderately correlated with the background selection statistic annotation^22^ (*R* = −0.21), defined as (1 - McVicker B statistic / 1000), where the McVicker B statistic quantifies a site’s genetic distance to its nearest exon.^30^ We have publicly released this annotation (see URLs).

To assess the informativeness of functionally important regions versus regions impacted by selection in explaining the depletions of squared trans-ethnic genetic correlation, we applied S-LDXR on the distance to nearest exon annotation together with the baseline-LD-X model annotations. We used both enrichment of squared trans-ethnic genetic correlation (*λ*^2^(*C*)) and standardized annotation effect size (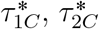 and 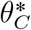) to assess informativeness.

### Analysis of probability of loss-of-function intolerance decile gene annotations

We created 10 annotations based on genes in deciles of probability of being loss-of-function intolerant (pLI) (see URLs), defined as the probability of assigning a gene into haplosufficient regions, where protein-truncating variants are depleted.^56^ Genes with high pLI (e.g. > 0.9) have higly constrained functionality, and therefore mutations in these genes are subject to negative selection. We included SNPs within a 100kb-base-pair window around each gene, following ref.^24^ A correlation heat map between pLI decile gene annotations and the 8 continuous-valued annotations is provided in Figure S29. All pLI decile gene annotations are moderately correlated with the background selection statistic and CpG content annotations.

### Analysis of the integrated haplotype score annotation

We created a binary annotation (proportion of SNPs: 6.3%) that includes all SNPs whose maximum absolute value of the integrated haplotype score (iHS)^43,44^ (see URLs) across all 1000 Genomes Project EAS and EUR sub-populations are greater than 2.0, the recommended threshold to detect positive selection in ref.^43^. This annotation is positively correlated with the top quintile of the background selection statistic annotation (*R* = 0.077). We note that although the iHS is a recombination-rate-adjusted quantity to detect the action of recent positive selection, it may also capture actions of negative selection.^43,44^

## Acknowledgements

We are grateful to L. O’Connor, H. Finucane, D. Kassler, S. Mallick, N. Patterson, B. Neale, R. Walters, A. Martin, B. Brown, F. Hormozdiari, M. Hujoel, K. Burch, and B. Pasaniuc for helpful discussions. This research was conducted using the UK Biobank Resource under Application 16549 and was funded by NIH grants R01 HG006399, U01 HG009379, R37 MH107649, R01 MH101244, and R01 CA222147.

