## Supplementary Notes for "Population-specific causal disease effect sizes in functionally important regions impacted by selection"

#### Causal and joint-fit effect size

Following ref,[1] we define causal effect size of a SNP as the underlying *true* effect size of the SNP on phenotype; we define joint-fit effect size of a SNP as the *inferred* effect size of the SNP. Causal effect size of a SNP is unique, whereas joint-fit effect size is subjected to the set of SNPs included in fitting the model for inferring the causal effect. Previous work[2] estimates trans-ethnic genetic correlation of joint-fit effect size – the set of SNPs for model fitting is the set of SNPs with minor allele frequency greater than 5% in both populations.[1,2] In this work, we focus on estimating trans-ethnic genetic correlation of causal effect size.

#### Per-allele and standardized causal effect size

Per-allele causal effect size of a SNP is the change in phenotype resulted from having an additional allele at that SNP. Standardized causal effect size of a SNP is the change in phenotype per standard deviation increase in normalized genotype of that SNP. Per-allele and standardized causal effect size of a SNP are related to each other through

$$\beta_{\text{standardized}} = \sqrt{2p(1-p)}\beta_{\text{per-allele}}, \tag{1}$$

where $p$ is the minor allele frequency (MAF) of the SNP in a population. Comparing standardized causal effect size of a SNP across populations is less informative due to differences in MAF. Thus, we focus on comparing per-allele causal effect size across populations in this work.

#### Defining stratified squared trans-ethnic genetic correlation of per-allele causal effect size

We model a complex phenotype in two populations using linear models,

$$\begin{aligned} \boldsymbol{Y}_1 &= \boldsymbol{X}_1\boldsymbol{\beta}_1 + \boldsymbol{\epsilon}_1, \\ \boldsymbol{Y}_2 &= \boldsymbol{X}_2\boldsymbol{\beta}_2 + \boldsymbol{\epsilon}_2, \end{aligned} \tag{2}$$

where $\boldsymbol{Y}_1 \in \mathbb{R}^{N_1}$ and $\boldsymbol{Y}_2 \in \mathbb{R}^{N_2}$ are vectors of standardized phenotype measurements with 0 mean and unit variance in the two populations, with sample size $N_1$ and $N_2$, respectively; $\boldsymbol{X}_1 \in \mathbb{R}^{N_1 \times M}$ and $\boldsymbol{X}_2 \in \mathbb{R}^{N_2 \times M}$ are mean centered (but *not normalized*) genotype matrices of the two populations across $M$ SNPs, respectively; $\boldsymbol{\beta}_1 \in \mathbb{R}^M$ and $\boldsymbol{\beta}_2 \in \mathbb{R}^M$ are the *per-allele* *causal* effect size vectors of the $M$ SNPs on phenotypes in the two populations, respectively; and $\boldsymbol{\epsilon}_1 \in \mathbb{R}^{N_1}$ and $\boldsymbol{\epsilon}_2 \in \mathbb{R}^{N_2}$ are the environmental effects in the two populations, respectively. We model per-allele causal effect sizes, instead of standardized effect sizes as is modeled in LDSC, to account for differences in minor allele frequency across different populations.

We assume both $\boldsymbol{X}_1$ and $\boldsymbol{X}_2$ to be random. We assume a random effect model for the per-allele causal effect sizes of SNP $j$ in the two populations, $\beta_{1j}$ and $\beta_{2j}$, respectively, with mean, variance, and covariance,

$$\begin{aligned} \mathrm{E}[\beta_{1j}] = 0, \mathrm{Var}[\beta_{1j}] &= \sum_C a_j(C)\tau_{1C}, \\ \mathrm{E}[\beta_{2j}] = 0, \mathrm{Var}[\beta_{2j}] &= \sum_C a_j(C)\tau_{2C}, \\ \mathrm{Cov}[\beta_{1j}, \beta_{2j}] &= \sum_C a_j(C)\theta_C, \end{aligned} \tag{3}$$

where $a_j(C)$ is SNP $j$'s value with respect to annotation $C$; $\tau_{1C}$ and $\tau_{2C}$ are the net contribution of annotation $C$ to the variance of per-allele causal effect size of SNP $j$ in the two populations; $\theta_C$ the net contribution of annotation $C$ to the co-variance of per-allele causal effect size of SNP $j$ in the two populations.

We define stratified trans-ethnic genetic co-variance of a binary annotation $C$ (e.g. functional annotations or quintiles of continuous-valued annotations) as the sum of per-SNP genetic covariance of SNPs that are a member of annotation $C$,

$$\rho_g(C) = \sum_{j \in C} \mathrm{Cov}[\beta_{1j}, \beta_{2j}] = \sum_{j \in C} \sum_{C'} a_j(C')\theta_{C'}. \tag{4}$$

Here, $C$ is a binary annotation, but $C'$ can be either binary or continuous-valued. Similarly, we define stratified heritability (of *per-allele* causal effect sizes) of a binary annotation $C$ in the two populations as,

$$\begin{aligned} h_{g1}^2(C) &= \sum_{j \in C} \mathrm{Var}[\beta_{1j}] = \sum_{j \in C} \sum_{C'} a_j(C')\tau_{1C'}, \\ h_{g2}^2(C) &= \sum_{j \in C} \mathrm{Var}[\beta_{2j}] = \sum_{j \in C} \sum_{C'} a_j(C')\tau_{2C'}. \end{aligned} \tag{5}$$

We define stratified trans-ethnic genetic correlation as

$$r_g(C) = \frac{\rho_g(C)}{\sqrt{h_{g1}^2(C) h_{g2}^2(C)}}. \tag{6}$$

Since estimates of $h^2_{g1}(C)$ and $h^2_{g2}(C)$ can be noisy and possibly negative, rendering the square roots undefined, we estimate stratified squared trans-ethnic genetic correlation instead, which is defined as,

$$r_g^2(C) = \frac{\rho_g^2(C)}{h^2_{g1}(C)h^2_{g2}(C)}. \tag{7}$$

Another advantage of estimating $r_g^2(C)$ over $r_g(C)$ is that taking square root of a random variable creates downward bias, which is difficult to correct for – estimating $r_g^2(C)$ resolves this issues. In this work, we only estimate $r_g^2(C)$ for SNPs with minor allele frequency (MAF) greater than 5% in both populations. Additionally, we define enrichment of stratified squared trans-ethnic genetic correlation,

$$\lambda^2(C) = \frac{r_g^2(C)}{r_g^2}, \tag{8}$$

as the ratio between stratified squared trans-ethnic genetic correlation of annotation $C$ and squared genome-wide trans-ethnic genetic correlation; we meta-analyze $\lambda^2(C)$ across different traits.

### Estimating stratified squared trans-ethnic genetic correlation

#### Regression equations to estimate $\theta_C$ and $\tau_C$

We estimate the net contributions of annotation to per-SNP trans-ethnic genetic covariance and per-allele heritability, $\theta_C$, $\tau_{1C}$ and $\tau_{2C}$, respectively, from GWAS summary association statistics using methods of moments.

In genome-wide association studies (GWAS) across two populations, Z-scores testing association between SNP $j$ and the trait are calculated as,

$$\begin{aligned} Z_{1j} &= \frac{1}{\sigma_{1j}\sqrt{N_1}}\boldsymbol{X}_{1j}^\intercal \boldsymbol{Y}_1, \\ Z_{2j} &= \frac{1}{\sigma_{2j}\sqrt{N_2}}\boldsymbol{X}_{2j}^\intercal \boldsymbol{Y}_2. \end{aligned} \tag{9}$$

where $Z_{1j}$ and $Z_{2j}$ are Z-scores for SNP $j$ in the two populations, respectively; $\sigma_{1j}$ and $\sigma_{2j}$ are the standard deviation of SNP $j$ in the two population.

Substituting the linear phenotype model from Equation (2), it can be shown that

$$
\begin{aligned}
\mathrm{E}[Z_{1j}Z_{2j}] &= \frac{1}{\sigma_{1j}\sigma_{2j}\sqrt{N_1N_2}} \mathrm{E}\left[(\boldsymbol{X}_{1j}^\intercal \boldsymbol{X}_1\boldsymbol{\beta}_1 + \boldsymbol{X}_{1j}^\intercal \boldsymbol{\epsilon}_1)(\boldsymbol{X}_{2j}^\intercal \boldsymbol{X}_2\boldsymbol{\beta}_2 + \boldsymbol{X}_{2j}^\intercal \boldsymbol{\epsilon}_2)\right] \\
&= \frac{1}{\sigma_{1j}\sigma_{2j}\sqrt{N_1N_2}} \mathrm{E}\left[\left(\boldsymbol{X}_{1j}^\intercal\left(\sum_{k=1}^M \boldsymbol{X}_{1k}\beta_{1k}\right)\right)\left(\boldsymbol{X}_{2j}^\intercal\left(\sum_{k=1}^M \boldsymbol{X}_{2k}\beta_{2k}\right)\right)\right] \\
&= \frac{1}{\sigma_{1j}\sigma_{2j}\sqrt{N_1N_2}} \mathrm{E}\left[\left(\sum_{k=1}^M \beta_{1k}\boldsymbol{X}_{1j}^\intercal\boldsymbol{X}_{1k}\right)\left(\sum_{k=1}^M \beta_{2k}\boldsymbol{X}_{2j}^\intercal\boldsymbol{X}_{2k}\right)\right] \\
&= \frac{1}{\sigma_{1j}\sigma_{2j}\sqrt{N_1N_2}} \mathrm{E}\left[\sum_{k=1}^M \beta_{1k}\beta_{2k}\left(\boldsymbol{X}_{1j}^\intercal\boldsymbol{X}_{1k}\right)\left(\boldsymbol{X}_{2j}^\intercal\boldsymbol{X}_{2k}\right)\right] \\
&= \frac{1}{\sigma_{1j}\sigma_{2j}\sqrt{N_1N_2}} \sum_{k=1}^M \mathrm{Cov}[\beta_{1k},\beta_{2k}]\,\mathrm{E}[\boldsymbol{X}_{1j}^\intercal\boldsymbol{X}_{1k}]\,\mathrm{E}[\boldsymbol{X}_{2j}^\intercal\boldsymbol{X}_{2k}] \\
&= \frac{1}{\sigma_{1j}\sigma_{2j}\sqrt{N_1N_2}} \sum_{k=1}^M \sum_C \theta_C a_k(C) N_1\rho_{1jk}N_2\rho_{2jk} \\
&= \sqrt{N_1N_2}\sum_C \left(\sum_{k=1}^M \frac{\rho_{1jk}\rho_{2jk}}{\sigma_{1j}\sigma_{2j}} a_k(C)\right)\theta_C,
\end{aligned}
\tag{10}
$$

where $\rho_{1jk}$ and $\rho_{2jk}$ are the covariances between SNP $j$ and $k$ in population 1 and population 2, respectively. Let

$$
\ell_\times(j,C) = \sum_{k=1}^M \frac{\rho_{1jk}\rho_{2jk}}{\sigma_{1j}\sigma_{2j}} a_k(C) \tag{11}
$$

be the trans-ethnic LD score of SNP $j$ with respect to annotation $C$, we arrive at the regression equation for estimating $\theta_C$,

$$
\mathrm{E}[Z_{1j}Z_{2j}|\ell_\times(j,C)] = \sqrt{N_1N_2}\sum_C \ell_\times(j,C)\theta_C. \tag{12}
$$

Following ref.[3], regression equations for estimating $\tau_{1C}$ and $\tau_{2C}$, contribution of annotation $C$ to per-SNP heritability, can be derived similarly,

$$
\begin{aligned}
\mathrm{E}[\chi_{1j}^2|\ell_1(j,C)] &= N_1\sum_C \ell_1(j,C)\tau_{1C} + N_1a_1 + 1, \\
\mathrm{E}[\chi_{2j}^2|\ell_2(j,C)] &= N_2\sum_C \ell_2(j,C)\tau_{2C} + N_2a_2 + 1,
\end{aligned}
\tag{13}
$$

where

$$
\ell_p(j,C) = \sum_{k=1}^M \frac{\rho_{pjk}^2}{\sigma_{pj}^2} a_k(C) \tag{14}
$$

is the LD score of SNP $j$ with respect to annotation $C$ in population $p$; and $a_p$ is the intercept term capturing population stratification in population $p$. An intercept term is not necessary in the regression in Equation (12), as GWAS from different populations are not expected to share samples or shared population stratification.

#### Estimating LD scores

We estimate trans-ethnic and population-specific LD scores using publicly available reference genotypes of 481 East Asian and 489 European individuals from the 1000 Genomes Project.[4]

Let $\boldsymbol{X}_1 \in \mathbb{R}^{N_1 \times M}$ and $\boldsymbol{X}_2 \in \mathbb{R}^{N_2 \times M}$ be the mean centered (but *not normalized*) reference genotype matrices for $M$ SNPs in the two populations, with reference sample size $N_1$ and $N_2$, respectively, we obtain unbiased estimates of trans-ethnic LD score of SNP $j$ with respect to annotation $C$, $\ell_\times(j, C)$ as

$$\hat{\ell}_\times(j, C) = \frac{1}{\hat{\sigma}_{1j}\hat{\sigma}_{2j}} \sum_{k=1}^{M} \hat{\rho}_{1jk}\hat{\rho}_{2jk}, \tag{15}$$

where

$$\hat{\rho}_{pjk} = \frac{\boldsymbol{X}_{pk}^\intercal \boldsymbol{X}_{pj}}{N_p - 1},\ \hat{\sigma}_{pj}^2 = \frac{\boldsymbol{X}_{pj}^\intercal \boldsymbol{X}_{pj}}{N_p - 1}. \tag{16}$$

At sample size of $N_1 = 481$ and $N_2 = 489$, both standard deviation estimation and ratio estimation are nearly unbiased. Thus, to show $\mathrm{E}[\hat{\ell}_\times(j, C)] = \ell_\times(j, C)$, it suffices to show $\mathrm{E}[\hat{\rho}_{1jk}\hat{\rho}_{2jk}] = \rho_{1jk}\rho_{2jk}$. Indeed,

$$\begin{aligned}
\mathrm{E}[\hat{\rho}_{1jk}\hat{\rho}_{2jk}] &= \mathrm{E}\left[\left(\frac{\boldsymbol{X}_{1k}^\intercal \boldsymbol{X}_{1j}}{N_1 - 1}\right)\left(\frac{\boldsymbol{X}_{2k}^\intercal \boldsymbol{X}_{2j}}{N_2 - 1}\right)\right] \\
&= \frac{1}{(N_1 - 1)(N_2 - 1)} \mathrm{E}\left[\sum_{i=1}^{N_1} \boldsymbol{X}_{1ik}\boldsymbol{X}_{1ij} \sum_{i'=1}^{N_2} \boldsymbol{X}_{2i'k}\boldsymbol{X}_{2i'j}\right] \\
&= \frac{1}{(N_1 - 1)(N_2 - 1)} \sum_{i=1}^{N_1}\sum_{i'=1}^{N_2} \mathrm{E}\left[\boldsymbol{X}_{1ik}\boldsymbol{X}_{1ij}\boldsymbol{X}_{2i'k}\boldsymbol{X}_{2i'j}\right] \\
&= \frac{(N_1 - 1)\rho_{1jk}(N_2 - 1)\rho_{2jk}}{(N_1 - 1)(N_2 - 1)} \\
&= \rho_{1jk}\rho_{2jk},
\end{aligned} \tag{17}$$

where the equality on the fourth line follows from Isserlis' theorem[5] and the fact that unadjusted sample covariance is biased by a factor of $\frac{N-1}{N}$. When estimating the trans-ethnic LD scores, we restrict to SNPs that are present in both populations. Effectively, we assume that only SNPs present in both populations contribute to genetic covariance. Since LD is small outside a 1 centimorgan window, we only include SNPs within a 1 centimorgan window in the summation in Equation (17), similar to previous works.[3,6,7]

Similarly, we obtain unbiased estimates of population-specific LD score, $\ell_p(j, C)$, as

$$\hat{\ell}_p(j, C) = \frac{1}{\hat{\sigma}^2_{pj}} \sum_{k=1}^{M} \frac{N_p}{N_p - 1} \left( \hat{\rho}^2_{pjk} - \frac{\hat{\sigma}^2_{pj}\hat{\sigma}^2_{pk}}{N_p - 1} \right). \tag{18}$$

For sample size of $N_1 = 481$ and $N_2 = 489$, the bias introduced in ratio estimation is negligible. Thus, to show $\hat{\ell}_p(j, C)$ is unbiased, it suffices to show $\mathrm{E}\left[\hat{\rho}^2_{pjk} - \frac{\hat{\sigma}^2_{pj}\hat{\sigma}^2_{pk}}{N_p}\right] = \frac{N_p-1}{N_p}\rho^2_{pjk}$. Indeed,

$$\begin{aligned}
&\mathrm{E}\left[\hat{\rho}^2_{pjk} - \frac{\hat{\sigma}^2_{pj}\hat{\sigma}^2_{pk}}{N_p}\right] = \mathrm{E}\left[\left(\frac{\boldsymbol{X}^\intercal_{pk}\boldsymbol{X}_{pj}}{N_p - 1}\right)^2 - \frac{\hat{\sigma}^2_{pj}\hat{\sigma}^2_{pk}}{N_p - 1}\right] \\
&= \left(\frac{1}{N_p - 1}\right)^2 \mathrm{E}\left[\sum_{i=1}^{N_p}\sum_{i'=1}^{N_p} \boldsymbol{X}_{pik}\boldsymbol{X}_{pij}\boldsymbol{X}_{pi'k}\boldsymbol{X}_{pi'j}\right] - \frac{\sigma^2_{pj}\sigma^2_{pk}}{N_p - 1} \\
&= \left(\frac{1}{N_p - 1}\right)^2 \sum_{i=1}^{N_p}\sum_{i'=1}^{N_p} \mathrm{E}\left[\boldsymbol{X}_{pik}\boldsymbol{X}_{pij}\boldsymbol{X}_{pi'k}\boldsymbol{X}_{pi'j}\right] - \frac{\sigma^2_{pj}\sigma^2_{pk}}{N_p - 1} \\
&= \left(\frac{1}{N_p - 1}\right)^2 \left[(N_p - 1)^2\rho^2_{pjk} + (N_p - 1)\sigma^2_j\sigma^2_k + (N_p - 1)\rho^2_{pjk}\right] - \frac{\sigma^2_{pj}\sigma^2_{pk}}{N_p - 1} \\
&= \frac{N_p - 1}{N_p}\rho^2_{jk}\,.
\end{aligned} \tag{19}$$

When estimating the trans-ethnic LD scores, we restrict to SNPs that are present in population $p$. Effectively, we assume that SNPs present in population $p$ contribute to heritability. Since LD is small outside a 1 centimorgan window, we only include SNPs within a 1 centimorgan window in the summation for estimating LD scores.

#### Regression SNPs and regression weights

To mitigate potential confounding due to imputation quality, we include only well-imputed SNPs (INFO>0.9) in the regression. We further restrict to HapMap 3[8] SNPs with minor allele frequency (estimated using 1000 Genomes Project[4] data) greater than 5% in both populations, which is a set of SNPs that are well imputed in diverse populations and has been used in previous studies.[3,7]

We use weighted least square regression to obtain estimates of $\tau_{1C}$, $\tau_{2C}$, and $\theta_C$. For estimating $\tau_{pC}$, we use weights similar to those described in Finucane et al 2015. In detail, the weights for each regression SNP $j$ in population $p$ is

$$w_{pj} = \frac{1}{\ell_p(j, \text{HapMap3}) \left(N_p \sum_C \ell_p(j, C)\tau_{pC} + 1\right)^2}. \tag{20}$$

For estimating $\theta_C$, we use the following weights

$$v_{pj} = \frac{1}{\sqrt{\prod_{p=1}^{2} \ell_p(j, \text{HapMap3})} \left[\prod_{p=1}^{2} \left(N_p \sum_C \ell_p(j, C)\tau_{pC} + 1\right) + N_p \sum_C \ell_\times(j, C)\theta_C\right]}. \tag{21}$$

#### Estimating stratified squared trans-ethnic genetic correlation

Let $\hat{\tau}_{1C}$, $\hat{\tau}_{2C}$, and $\hat{\theta}_C$, be the estimates of $\tau_{1C}$, $\tau_{2C}$, and $\theta_C$, respectively. First, we obtain estimates of stratified trans-ethnic genetic covariance and heritability of a binary annotation $C$ as,

$$\begin{aligned} \hat{\rho}_g(C) &= \sum_{j \in C} \sum_{C'} a_{C'}(j)\hat{\theta}_{C'}, \\ \hat{h}^2_{g1}(C) &= \sum_{j \in C} \sum_{C'} a_{C'}(j)\hat{\tau}_{1C'}, \\ \hat{h}^2_{g2}(C) &= \sum_{j \in C} \sum_{C'} a_{C'}(j)\hat{\tau}_{2C'}. \end{aligned} \tag{22}$$

We jackknife over 200 continuous and disjoint blocks of SNPs to obtain standard error of each estimates. As, an example, we estimate standard error of $\hat{\rho}_g(C)$ as

$$\text{S.E.}[\hat{\rho}_g(C)] = \sqrt{\frac{B-1}{B} \sum_{b=1}^{B} \left[\hat{\rho}_g(C) - \hat{\rho}_g^{(b)}(C)\right]^2}, \tag{23}$$

where $B$ is the total number of jackknife samples, and $\hat{\rho}_g^{(b)}(C)$ denotes the estimate with SNPs in the $b$-th block removed.

Next, we obtain an initial estimate of stratified squared trans-ethnic genetic correlation, $r_g^2(C)$, as

$$\tilde{r}_g^2(C) = \frac{\hat{\rho}_g^2(C) - (\text{S.E.}[\hat{\rho}_g(C)])^2}{\hat{h}^2_{g1}(C)\hat{h}^2_{g2}(C) - \text{Cov}[\hat{h}^2_{g1}(C), \hat{h}^2_{g2}(C)]}, \tag{24}$$

where $\text{Cov}[\hat{h}^2_{g1}(C), \hat{h}^2_{g2}(C)]$ is estimated using jackknife over 200 continuous and disjoint blocks of SNPs,

$$\mathrm{Cov}[\hat{h}^2_{g1}(C), \hat{h}^2_{g2}(C)] = \frac{B-1}{B} \sum_{b=1}^{B} \left[\hat{h}^2_{g1}(C) - \hat{h}^{2(b)}_{g1}(C)\right] \left[\hat{h}^2_{g2}(C) - \hat{h}^{2(b)}_{g2}(C)\right]. \quad (25)$$

The initial estimator, $\tilde{r}^2_g(C)$, however, is a biased estimator of $r^2_g(C)$.[9] We estimate and correct for the bias using jackknife samples of $\tilde{r}^2_g(C)$.[9] We obtain final bias-corrected estimate of $r^2_g(C)$ as,

$$\hat{r}^2_g(C) = \left\{ \tilde{r}^2_g(C) + \frac{\mathrm{Cov}\left[\hat{\rho}^2_g(C), \hat{h}^2_{g1}(C)\hat{h}^2_{g2}(C)\right]}{\hat{h}^2_{g1}(C)\hat{h}^2_{g2}(C)} \right\} \Big/ \left\{ 1 + \frac{\mathrm{Var}\left[\hat{h}^2_{g1}(C)\hat{h}^2_{g2}(C)\right]}{\hat{h}^2_{g1}(C)\hat{h}^2_{g2}(C)} \right\}, \quad (26)$$

and obtain its standard error using block jackknife.

We obtain an initial estimate enrichment of stratified squared trans-ethnic genetic correlation

$$\tilde{\lambda}^2(C) = \frac{\hat{r}^2_g(C)}{\hat{r}^2_g}, \quad (27)$$

and obtain bias corrected enrichment as

$$\hat{\lambda}^2(C) = \left\{ \tilde{\lambda}^2(C) + \frac{\mathrm{Cov}\left[\hat{r}^2_g(C), \hat{r}^2_g\right]}{\hat{r}^2_g(C)} \right\} \Big/ \left\{ 1 + \frac{\mathrm{Var}\left[\hat{r}^2_g(C)\right]}{\hat{r}^2_g(C)} \right\} \quad (28)$$

#### Shrinkage estimator

Estimates of $r^2_g(C)$ can be imprecise and unreliable if the denominator, $h^2_{g1}(C)h^2_{g2}(C)$, is noisy and close to 0. This is especially true for small annotations. To mitigate this issue, we introduce a shrinkage estimator to "regularize" the estimates of $r^2_g(C)$.

We apply the shrinkage to estimates of stratified per-SNP genetic covariance and heritability, so that the per-SNP estimates are shrunk towards genome-wide average. Inspired by Bayesian shrinkage, we derive a shrinkage factor for per-SNP genetic covariance and heritability as follows. Let

$$
\begin{aligned}
\gamma_1 &= 1/\left(1+\alpha\frac{\mathrm{Var}\left[\hat{\rho}_g(C)\right]}{\mathrm{Var}\left[\hat{\rho}_g\right]}\frac{M}{M_C}\right),\\
\gamma_2 &= 1/\left(1+\alpha\frac{\mathrm{Var}\left[\hat{h}_{g1}(C)\right]}{\mathrm{Var}\left[\hat{h}^2_{g1}\right]}\frac{M}{M_C}\right),\\
\gamma_3 &= 1/\left(1+\alpha\frac{\mathrm{Var}\left[\hat{h}_{g2}(C)\right]}{\mathrm{Var}\left[\hat{h}^2_{g2}\right]}\frac{M}{M_C}\right),
\end{aligned}
\tag{29}
$$

where $M_C$ is the number of SNPs in annotation $C$, and $\alpha \in [0,1]$ is a user-controlled tuning parameter that governs the magnitude of shrinkage. We define the shared shrinkage factor as

$$
\gamma = \min\{\gamma_1, \gamma_2, \gamma_3\}. \tag{30}
$$

We use shared shrinkage factor instead of separate shrinkage factors for convenience of characterizing the behavior of the estimator. When $\alpha$ is set to 0, no shrinkage is applied; when $\alpha$ is set to 1, the entire Bayesian shrinkage is applied.

We apply the shrinkage to stratified genetic covariance and heritability as follows,

$$
\begin{aligned}
\bar{\rho}_g(C) &= M_C\left(\gamma\frac{\hat{\rho}_g(C)}{M_C}+(1-\gamma)\frac{\hat{\rho}_g}{M}\right)\\
\bar{h}^2_{g1}(C) &= M_C\left(\gamma\frac{\hat{h}^2_{g1}(C)}{M_C}+(1-\gamma)\frac{\hat{h}^2_{g1}}{M}\right)\\
\bar{h}^2_{g2}(C) &= M_C\left(\gamma\frac{\hat{h}^2_{g2}(C)}{M_C}+(1-\gamma)\frac{\hat{h}^2_{g2}}{M}\right),
\end{aligned}
\tag{31}
$$

and obtain standard errors of the shrunk estimates using block jackknife. Intuitively, if stratified heritability and trans-ethnic genetic covariance are estimated with low variance, the amount of shrinkage needed will be small, and shrinkage estimator will preserve the unshrunk estimates. On the other hand, if stratified heritability and genetic covariance are estimated with large variance (i.e. noisy), the shrinkage estimator will shrink the estimates towards genome-wide average.

Finally, we obtain shrunk $r_g^2(C)$ and $\lambda^2(C)$, $\bar{r}_g^2(C)$ and $\bar{\lambda}^2(C)$, by plugging in $\bar{\rho}_g(C)$, $\bar{h}^2_{g1}(C)$, and $\bar{h}^2_{g2}(C)$ into the procedures described in previous section. We found that when $\alpha = 0.5$, the shrinkage estimator yields robust results across a wide range of polygenicity.

### Two-population Eyre-Walker model

The Eyre-Walker model[10] couples fitness effect (selection coefficient) with causal disease effect size, $\beta$, through the equation

$$\beta = \delta S^{\tau}(1+\epsilon), \tag{32}$$

where $\delta = \pm 1$ with equal probabilities governs the sign of $\beta$; $S = 4sN_e$ ($s$ is the fitness effect, $N_e$ effective sample size of the population); $\tau$ is the parameter coupling selection and $\beta$; and $\epsilon$ is normally distributed with mean 0 and variance $\sigma_e^2$. Since the scaling factor $4N_e$ does not affect trans-ethnic genetic correlation (and subsequently enrichment of stratified squared trans-ethnic genetic correlation, $\lambda^2(C)$), we use the simplified equation instead,

$$\beta \propto \delta s^{\tau}(1+\epsilon). \tag{33}$$

We use negative $s$ to denote deleteriousness, following convention of previous works.[11,12] However, we emphasize that positive $s$ (i.e. beneficial mutations) is also plausible.

We extend the Eyre-Walker model to two populations to model causal disease effect sizes of SNP $j$, $\beta_{1j}$ and $\beta_{2j}$, in population 1 and population 2, respectively,

$$\begin{aligned} \beta_{1j} &\propto \delta s_{1j}^{\tau}(1+\epsilon_1), \\ \beta_{2j} &\propto \delta s_{2j}^{\tau}(1+\epsilon_2), \end{aligned} \tag{34}$$

where $s_{1j}$ and $s_{2j}$ are the fitness effects of SNP $j$ in the two populations; $\epsilon_1$ and $\epsilon_2$ independently follow normal distributions with mean 0 and variance $\sigma_1^2$ and $\sigma_2^2$. Assuming $\tau$ is a constant, $\beta_{1j}$ and $\beta_{2j}$ has covariance,

$$\text{Cov}[\beta_{1j}, \beta_{2j}] \propto \text{E}[\delta s_{1j}^{\tau}(1+\epsilon_1)\delta s_{2j}^{\tau}(1+\epsilon_2)] = E[(s_{1j}s_{2j})^{\tau}], \tag{35}$$

and variance,

$$\begin{aligned} \text{Var}[\beta_{1j}] &\propto \text{E}[(\delta s_{1j}^{\tau}(1+\epsilon_1))^2] = \text{E}[s_{1j}^{2\tau}](1+\sigma_1^2), \\ \text{Var}[\beta_{2j}] &\propto \text{E}[(\delta s_{2j}^{\tau}(1+\epsilon_2))^2] = \text{E}[s_{2j}^{2\tau}](1+\sigma_2^2). \end{aligned} \tag{36}$$

The squared genome-wide trans-ethnic genetic correlation is then

$$\begin{aligned} r_g^2 &= \frac{(\sum_j \mathrm{E}[(s_{1j}s_{2j})^\tau])^2}{(\sum_j \mathrm{E}[s_{1j}^{2\tau}](1+\sigma_e^2)\,\mathrm{E}[s_{2j}^{2\tau}](1+\sigma_e^2))} \\ &= \frac{1}{(1+\sigma_1^2)(1+\sigma_2^2)} \frac{(\sum_j \mathrm{E}[(s_{1j}s_{2j})^\tau])^2}{\sum_j \mathrm{E}[s_{1j}^{2\tau}]\,\mathrm{E}[s_{2j}^{2\tau}]}. \end{aligned} \tag{37}$$

And the stratified squared trans-ethnic genetic correlation of a binary annotation $C$ is

$$\begin{aligned} r_g^2(C) &= \frac{(\sum_{j\in C} \mathrm{E}[(s_{1j}s_{2j})^\tau])^2}{(\sum_{j\in C} \mathrm{E}[s_{1j}^{2\tau}](1+\sigma_e^2)\,\mathrm{E}[s_{2j}^{2\tau}](1+\sigma_e^2))} \\ &= \frac{1}{(1+\sigma_1^2)(1+\sigma_2^2)} \frac{(\sum_{j\in C} \mathrm{E}[(s_{1j}s_{2j})^\tau])^2}{\sum_{j\in C} \mathrm{E}[s_{1j}^{2\tau}]\,\mathrm{E}[s_{2j}^{2\tau}]}. \end{aligned} \tag{38}$$

The enrichment of squared trans-ethnic genetic correlation, $\lambda^2(C)$, only depends on $s_{1j}$ and $s_{2j}$,

$$\lambda^2(C) = \frac{r_g^2(C)}{r_g^2} = \frac{(\sum_{j\in C} \mathrm{E}[(s_{1j}s_{2j})^\tau])^2}{(\sum_{j\in C} \mathrm{E}[s_{1j}^{2\tau}]\,\mathrm{E}[s_{2j}^{2\tau}])} \frac{(\sum_j \mathrm{E}[s_{1j}^{2\tau}]\,\mathrm{E}[s_{2j}^{2\tau}])}{(\sum_j \mathrm{E}[(s_{1j}s_{2j})^\tau])^2}. \tag{39}$$

Therefore, although $r_g^2$ can be less than 1 as long as $\sigma_1^2$ or $\sigma_2^2$ is greater than 0, differential fitness effects in annotation $C$ compared with genome-wide average is necessary for $\lambda^2(C)$ to be different from 1.

To introduce population-specific fitness effects, we assume

$$\begin{aligned} s_1 &= s_0(1+\Delta_1), \\ s_2 &= s_0(1+\Delta_2), \end{aligned} \tag{40}$$

where $s_0$ represents the fitness effect prior to the split of population 1 and population 2, and $\Delta_1$ and $\Delta_2$ represent the relative change in fitness effects since the split, and are independently sampled from $N(0, \sigma_\Delta^2)$ (and truncated so that $(1+\Delta_1)$ and $(1+\Delta_2)$ are non-negative). We further assume that $\sigma_\Delta^2$ is small (close to zero) at weakly deleterious or effectively neutral SNPs (i.e. $s_1 \approx s_2$), and large at more strongly deleterious SNPs (i.e. $s_1 \neq s_2$) (Figure S31a). Under our model, fitness effects in the two populations have the same mean, but higher variance at SNPs with large fitness effect (i.e. strongly deleterious SNPs) and lower variance at SNPs with small fitness effect (i.e. weakly deleterious SNPs). Since both populations have the same mean fitness effect, we expect the relationship between effect size and MAF to be the same in the two populations for strongly and weakly deleterious SNPs. We have publicly released Python code implementing the 2-population Eyre-Walker model (see URLs).

We used Equation (40) to sample population-specific fitness effects ($s_1$ and $s_2$) and subsequently used Equation (34) to sample causal disease effect sizes ($\beta_1$ and $\beta_2$) for 50,000 simulated unlinked SNPs, setting 90% of the SNPs to be weakly deleterious ($s_0 = -10^{-5}$) and 10% of the SNPs to be more strongly deleterious ($s_0 = -10^{-4}$) (Methods). We then used the sampled causal effect sizes to compute the enrichment/depletion of squared trans-ethnic genetic correlation ($\lambda^2(C)$) for SNPs in each of these two categories. When $\tau = 0.2$, $\sigma_1^2 = \sigma_2^2 = 1.0$, and $\sigma_\Delta^2 = 0.0$ for both weakly and more strongly deleterious SNPs (i.e. same fitness effects across populations), $\lambda^2(C)$ was equal to 1.00 (s.e. 0.00) for both categories (Figure S31b). However, when $\sigma_\Delta^2 = 0.0$ for weakly deleterious SNPs but $\sigma_\Delta^2 = 0.7$ for more strongly deleterious SNPs (leaving all other parameters unchanged), $\lambda^2(C)$ for more strongly deleterious SNPs decreased to 0.79 (s.e. 0.01) (Figure S31c) due to more population-specific causal disease effect sizes, roughly matching results for SNPs in the top quintile of background selection statistic in real data analyses (Figure 2). Analyses at other values of $\tau$ produced similar results, yielding lower values of $\lambda^2(C)$ for more strongly deleterious SNPs at higher values of $\sigma_\Delta^2$ (Table S27). We also performed a secondary analysis with $\sigma_\Delta^2 = 0.7$ for both weakly and more strongly deleterious SNPs (leaving all other parameters unchanged). We observed no depletion of $\lambda^2(C)$ at more strongly deleterious SNPs (Figure S32). Thus, we concluded that, under the Eyre-Walker evolutionary model, a lower $\sigma_\Delta^2$ at weakly deleterious SNPs and a higher $\sigma_\Delta^2$ at more strongly deleterious SNPs is necessary to explain the results observed in analyses of real traits.

Here, we did not consider demographic histories in our evolutionary modeling, which may lead to increased proportions of population-specific variants, decreasing trans-ethnic polygenic risk score accuracy.[13] We also note that other evolutionary models[14,15] exist, and could also be explored.[14,15]

### Supplementary tables

Table S1: **List of baseline-LD-X model annotations.** **a**) Summary information for functional annotations. **b**) Summary information for continuous-valued annotations. **c**) Correlation between functional and continuous-valued annotations. **d**) Correlation between continuous-valued annotations.

See attached Excel file.

Table S2: **List of the specifically expressed gene (SEG) annotations.** **a)** Mean background selection statistic of SEG annotations. **b)** Correlation between SEG annotation and continuous-valued annotations.

See attached Excel file.

| **simulated $r_g$** | **estimated $r_g$** | **s.e. mean** | **mean jackknife s.e.** |
|---|---|---|---|
| 0 | 0 | 0.0016 | 0.0016 |
| 0.2 | 0.21 | 0.0015 | 0.0016 |
| 0.4 | 0.41 | 0.0016 | 0.0017 |
| 0.6 | 0.62 | 0.0017 | 0.0017 |
| 0.8 | 0.82 | 0.0018 | 0.0019 |
| 1 | 1.03 | 0.002 | 0.0021 |

Table S3: **Numerical results of S-LDXR in estimating genome-wide trans-ethnic genetic correlation.** Mean and standard errors are based on 1,000 simulations.

Table S4: **Numerical results of S-LDXR in null simulations with 1% causal SNPs.** The shrinkage parameter, $\alpha$, is set to 0.0 in **a**, 0.25 in **b**, 0.5 in **c**, 0.75 in **d**, and 1.0 in **e**. Positive rate is defined as the proportion of tests with p-value less than 0.05. Standard error of the positive rate is obtained using jackknife.

See attached Excel file.

Table S5: **Numerical results of S-LDXR in null simulations with 10% causal SNPs.** The shrinkage parameter, $\alpha$, is set to 0.0 in **a**, 0.25 in **b**, 0.5 in **c**, 0.75 in **d**, and 1.0 in **e**. Positive rate is defined as the proportion of tests with p-value less than 0.05. Standard error of the positive rate is obtained using jackknife.

See attached Excel file.

Table S6: **Numerical results of S-LDXR in null simulations with 100% causal SNPs.** The shrinkage parameter, $\alpha$, is set to 0.0 in **a**, 0.25 in **b**, 0.5 in **c**, 0.75 in **d**, and 1.0 in **e**. Positive rate is defined as the proportion of tests with p-value less than 0.05. Standard error of the positive rate is obtained using jackknife.

See attached Excel file.

Table S7: **Numerical results of S-LDXR in causal simulations with 1% causal SNPs.** The shrinkage parameter, $\alpha$, is set to 0.0 in **a**, 0.25 in **b**, 0.5 in **c**, 0.75 in **d**, and 1.0 in **e**. Positive rate is defined as the proportion of tests with p-value less than 0.05. Standard error of the positive rate is obtained using jackknife.

See attached Excel file.

Table S8: **Numerical results of S-LDXR in causal simulations with 10% causal SNPs.** The shrinkage parameter, $\alpha$, is set to 0.0 in **a**, 0.25 in **b**, 0.5 in **c**, 0.75 in **d**, and 1.0 in **e**. Positive rate is defined as the proportion of tests with p-value less than 0.05. Standard error of the positive rate is obtained using jackknife.

See attached Excel file.

Table S9: **Numerical results of S-LDXR in causal simulations with 100% causal SNPs.** The shrinkage parameter, $\alpha$, is set to 0.0 in **a**, 0.25 in **b**, 0.5 in **c**, 0.75 in **d**, and 1.0 in **e**. Positive rate is defined as the proportion of tests with p-value less than 0.05. Standard error of the positive rate is obtained using jackknife.

See attached Excel file.

Table S10: **Numerical results of S-LDXR in null simulations with annotation-dependent MAF-dependent genetic architectures.** 10% of SNPs were randomly selected to be causal. The shrinkage parameter, $\alpha$, is set to 0.0 in **a**, 0.25 in **b**, 0.5 in **c**, 0.75 in **d**, and 1.0 in **e**. Positive rate is defined as the proportion of tests with p-value less than 0.05. Standard error of the positive rate is obtained using jackknife.

See attached Excel file.

Table S11: **Numerical results of S-LDXR in null simulations with annotation-dependent MAF-dependent genetic architectures and 5 MAF bins added to the baseline-LD-X model.** 10% of SNPs were randomly selected to be causal. The shrinkage parameter, $\alpha$, is set to 0.0 in **a**, 0.25 in **b**, 0.5 in **c**, 0.75 in **d**, and 1.0 in **e**. Positive rate is defined as the proportion of tests with p-value less than 0.05. Standard error of the positive rate is obtained using jackknife.

See attached Excel file.

Table S12: **Numerical results of S-LDXR in null simulations where causal variants differ across the two populations.** Here, 10% of SNPs were randomly selected to be causal in each population, with 80% of the causal variants in each population shared with the other population. The shrinkage parameter, $\alpha$, is set to 0.0 in **a**, 0.25 in **b**, 0.5 in **c**, 0.75 in **d**, and 1.0 in **e**. Positive rate is defined as the proportion of tests with p-value less than 0.05. Standard error of the positive rate is obtained using jackknife.

See attached Excel file.

Table S13: **Numerical results of S-LDXR in null simulations using all simulated GWAS samples and half (250) the default reference sample size.** Here, 10% of SNPs were randomly selected to be causal. Shrinkage level, $\alpha$, was set to 0.5. **a**) Estimates of $\lambda^2(C)$ for binary annotations and quintiles of continuously valued annotation in simulations under the baseline-LD-X model. **b**) Estimates of $\lambda^2(C)$ for binary annotations and quintiles of continuously valued annotation in simulations under the model involving annotation-dependent and MAF-dependent genetic architectures. Mean and standard errors were obtained across 1,000 simulations.

See attached Excel file.

Table S14: **Numerical results of S-LDXR in null simulations using all simulated GWAS samples and twice (1,000) the default reference sample size.** Here, 10% of SNPs were randomly selected to be causal. Shrinkage level, $\alpha$, was set to 0.5. **a**) Estimates of $\lambda^2(C)$ for binary annotations and quintiles of continuously valued annotation in simulations under the baseline-LD-X model. **b**) Estimates of $\lambda^2(C)$ for binary annotations and quintiles of continuously valued annotation in simulations under the model involving annotation-dependent and MAF-dependent genetic architectures. Mean and standard errors were obtained across 1,000 simulations.

See attached Excel file.

Table S15: **Numerical results of S-LDXR in null simulations using half of the simulated GWAS samples and default (500) reference samples.** Here, 10% of SNPs were randomly selected to be causal. Shrinkage level, $\alpha$, was set to 0.5. **a**) Estimates of $\lambda^2(C)$ for binary annotations and quintiles of continuously valued annotation in simulations under the baseline-LD-X model. **b**) Estimates of $\lambda^2(C)$ for binary annotations and quintiles of continuously valued annotation in simulations under the model involving annotation-dependent and MAF-dependent genetic architectures. Mean and standard errors were obtained across 1,000 simulations.

See attached Excel file.

| trait (abbrev.) | $N_{EAS}$ | $N_{EUR}$ | $h^2_{g,EAS}$ | $h^2_{g,EUR}$ | $r_g$ |
|---|---|---|---|---|---|
| *Atrial Fibrillation (AF) | 36792[16] | 1030836[17] | 0.110 (0.026) | 0.021 (0.002) | 0.817 (0.193) |
| Age at Menarche (AMN) | 67029[18] | 252514[19] | 0.074 (0.013) | 0.128 (0.010) | 0.878 (0.057) |
| Age at Menopause (AMP) | 43861[18] | 69360[19] | 0.092 (0.021) | 0.190 (0.016) | 0.567 (0.091) |
| Basophil Count (BASO) | 62076[20] | 131860[21] | 0.107 (0.018) | 0.088 (0.011) | 0.427 (0.061) |
| Body Mass Index (BMI) | 158284[20] | 337539[22] | 0.161 (0.010) | 0.207 (0.007) | 0.804 (0.021) |
| Blood Sugar (BS) | 93146[20] | 337539[22] | 0.057 (0.011) | 0.036 (0.004) | 0.829 (0.087) |
| Diastolic Blood Pressure (DBP) | 136615[20] | 337539[22] | 0.052 (0.008) | 0.146 (0.007) | 0.862 (0.059) |
| Estimated Glomerular Filtration Rate (EGFR) | 143658[20] | 100125[23] | 0.074 (0.008) | 0.058 (0.007) | 1.053 (0.063) |
| Eosinophil Count (EO) | 62076[20] | 337539[22] | 0.076 (0.016) | 0.154 (0.010) | 0.950 (0.092) |
| Hemoglobin A1c (HBA1C) | 42790[20] | 337539[22] | 0.109 (0.022) | 0.082 (0.006) | 0.875 (0.083) |
| High Density Lipoprotein (HDL) | 70657[20] | 337539[22] | 0.109 (0.016) | 0.140 (0.010) | 0.892 (0.056) |
| Height (HEIGHT) | 151569[24] | 337539[22] | 0.371 (0.017) | 0.366 (0.018) | 0.897 (0.018) |
| Hemoglobin (HGB) | 108769[20] | 132596[21] | 0.070 (0.010) | 0.166 (0.012) | 0.911 (0.058) |
| Hematocrit (HTC) | 108757[20] | 132699[21] | 0.078 (0.009) | 0.161 (0.012) | 0.870 (0.054) |
| Low Density Lipoprotein (LDL) | 72866[20] | 337539[22] | 0.047 (0.015) | 0.076 (0.009) | 0.662 (0.105) |
| Lymphocyte Count (LYMPH) | 62076[20] | 337539[22] | 0.121 (0.015) | 0.165 (0.011) | 0.903 (0.059) |
| Mean Corpuscular Hemoglobin (MCH) | 108054[20] | 337539[22] | 0.130 (0.014) | 0.144 (0.010) | 0.884 (0.049) |
| MCH Concentration (MCHC) | 108728[20] | 132586[21] | 0.069 (0.010) | 0.089 (0.010) | 0.887 (0.077) |
| Mean Corpuscular Volume (MCV) | 108256[20] | 132353[21] | 0.146 (0.015) | 0.200 (0.015) | 0.891 (0.048) |
| *Major Depressive Disorder (MDD) | 10640[25] | 62984[26] | 0.354 (0.078) | 0.202 (0.014) | 0.342 (0.074) |
| Monocyte Count (MONO) | 62076[20] | 337539[22] | 0.123 (0.015) | 0.156 (0.012) | 0.811 (0.048) |
| Neutrophil Count (NEUT) | 62076[20] | 131564[21] | 0.123 (0.016) | 0.163 (0.011) | 0.766 (0.059) |
| Platelet Count (PLT) | 108208[20] | 337539[22] | 0.157 (0.015) | 0.214 (0.013) | 0.879 (0.035) |
| *Rheumatoid Arthritis (RA) | 22343[27] | 37598[27] | 0.219 (0.041) | 0.191 (0.021) | 0.872 (0.098) |
| Red Blood Cell Count (RBC) | 108794[20] | 337539[22] | 0.105 (0.011) | 0.167 (0.009) | 0.924 (0.052) |
| Systolic Blood Pressure (SBP) | 136597[20] | 337539[22] | 0.064 (0.008) | 0.149 (0.007) | 0.807 (0.043) |
| *Schizophrenia (SCZ) | 13761[28] | 35737[28] | 0.908 (0.064) | 0.868 (0.040) | 0.945 (0.036) |
| *Type 2 Diabetes (T2D) | 190559[29] | 141364[30] | 0.099 (0.007) | 0.046 (0.006) | 0.927 (0.048) |
| Total Cholesterol (TC) | 128305[20] | 337539[22] | 0.057 (0.013) | 0.087 (0.010) | 0.910 (0.073) |
| Triglyceride (TG) | 105597[20] | 337539[22] | 0.061 (0.010) | 0.100 (0.009) | 0.932 (0.066) |
| White Blood Cell Count (WBC) | 107964[20] | 337539[22] | 0.103 (0.010) | 0.156 (0.007) | 0.848 (0.037) |

Table S16: **Details of 31 diseases and complex traits analyzed.** We report genome-wide heritability of the traits estimated using S-LDSC[3,11] conditioned on baseline-LD-v2.2 model annotations in each population, and trans-ethnic genetic correlation estimated using S-LDXR conditioned on baseline-LD-X model annotations. Heritability estimates for binary traits denote observed-scale heritability (* denotes binary traits). Standard errors of the estimates are shown in parentheses. The prevalence of MDD is 2.2% and 7.3%[31] in UK Biobank[32] EAS (Chinese) and EUR population, respectively. The prevalence of schizophrenia (SCZ) is 0.33% and 0.52% in Asia and Europe, respectively.[33] The prevalence of type 2 diabetes (T2D) is 2.7% and 4.2%[31] in UK Biobank EAS (Chinese) and EUR populations.

Table S17: **Numerical S-LDXR results for quintiles of 8 continuous-valued annotations across 31 diseases and complex traits.** The shrinkage parameter, $\alpha$, was set to 0.0 in **a**, 0.5 in **b**, and 1.0 in **c**. **d**) Here, results were meta-analyzed across a subset of 20 approximately independent traits with default shrinkage parameter ($\alpha = 0.5$).

See attached Excel file.

Table S18: **Numerical S-LDXR results for 28 binary functional annotations across 31 diseases and complex traits.** The shrinkage parameter, $\alpha$, was set to 0.0 in **a**, 0.5 in **b**, and 1.0 in **c**. **d**) Here, results were meta-analyzed across a subset of 20 approximately independent traits with default shrinkage parameter ($\alpha = 0.5$).

See attached Excel file.

| **decile** | $h^2_{g,EAS}(C)$ **enrch.** | $h^2_{g,EUR}(C)$ **enrch.** | $\lambda^2(C)$ **(s.e.)** |
|---|---|---|---|
| 1st | NA | NA | NA |
| 2nd | NA | NA | NA |
| 3rd | NA | NA | NA |
| 4th | NA | NA | NA |
| 5th | 1.04 (0.15) | 1.17 (0.08) | 0.92 (0.09) |
| 6th | 1.02 (0.08) | 1.05 (0.04) | 0.91 (0.04) |
| 7th | 1.01 (0.05) | 0.99 (0.03) | 1.04 (0.03) |
| 8th | 1.04 (0.04) | 1.07 (0.03) | 0.97 (0.02) |
| 9th | 0.99 (0.04) | 0.99 (0.02) | 0.97 (0.02) |
| 10th | 0.91 (0.03) | 0.90 (0.02) | 0.95 (0.03) |

Table S19: **Enrichment of heritability and stratified squared trans-ethnic genetic correlation across 10 MAF bin annotations.** MAF bins containing no SNP with MAF $> 5\%$ in either East Asian (EAS) or European (EUR) populations are reported as "NA". Standard errors are reported in parentheses.

Table S20: **Numerical S-LDXR results of observed $\lambda^2(C)$ vs. expected $\lambda^2(C)$ based on 8 continuous-valued annotations for 20 binary annotations across 31 diseases and complex traits.** The shrinkage parameter, $\alpha$, was set to 0.0 in **a**, 0.5 in **b**, and 1.0 in **c**.

See attached Excel file.

Table S21: **Numerical S-LDXR results for 53 specifically expressed gene (SEG) annotations across 31 diseases and complex traits.** While heritability enrichment may be impacted by choice of diseases and traits, $\lambda^2(C)$ is not expected to be disease specific. The shrinkage parameter, $\alpha$, was set to 0.0 in **a**, 0.5 in **b**, and 1.0 in **c**.

See attached Excel file.

Table S22: **Numerical S-LDXR results for 53 specifically expressed gene (SEG) annotations for 14 blood-related traits vs. 17 other traits.** The list of 14 blood phenotypes is: BASO, EO, HBA1C, HGB, HTC, LYMPH, MCH, MCHC, MCV, MONO, NEUT, PLT, RBC, WBC. The list of 17 non-blood phenotype is: AF, AMN, AMP, BMI, BS, DBP, EGFR, HEIGHT, HDL, LDL, MDD, RA, SBP, SCZ, TC, TG, T2D. Full name of the abbreviations can be found in Table S16. Here, the shrinkage parameter $\lambda^2(C)$ was set to the default of 0.5.

Table S23: **Numerical S-LDXR results of observed $\lambda^2(C)$ vs. expected $\lambda^2(C)$ based on 8 continuous-valued annotations for 53 specifically expressed gene (SEG) annotations across 31 diseases and complex traits.** While heritability enrichment may be impacted by choice of diseases and traits, $\lambda^2(C)$ is not expected to be disease specific. The shrinkage parameter, $\alpha$, was set to 0.0 in **a**, 0.5 in **b**, and 1.0 in **c**.

See attached Excel file.

Table S24: **Numerical S-LDXR results of $\lambda^2(C)$ for 53 specifically expressed gene (SEG) annotations for each of the 31 diseases and complex traits.** The shrinkage parameter, $\alpha$, was set to the default value of 0.5.

See attached Excel file.

| quintile | $-1\times$ **distance to nearest exon** | | | **background selection statistic** | | |
|---|---|---|---|---|---|---|
| | $h^2_{g,EAS}(C)$ **enrch.** | $h^2_{g,EUR}(C)$ **enrch.** | $\lambda^2(C)$ | $h^2_{g,EAS}(C)$ **enrch.** | $h^2_{g,EUr}(C)$ **enrch.** | $\lambda^2(C)$ |
| 1st | 0.24 (0.030) | 0.23 (0.023) | 1.07 (0.050) | 0.44 (0.026) | 0.43 (0.020) | 1.23 (0.052) |
| 2nd | 0.59 (0.019) | 0.65 (0.013) | 1.16 (0.028) | 0.68 (0.014) | 0.68 (0.011) | 1.19 (0.025) |
| 3rd | 0.89 (0.020) | 0.92 (0.014) | 1.03 (0.022) | 0.88 (0.010) | 0.89 (0.0073) | 1.06 (0.011) |
| 4th | 1.15 (0.028) | 1.15 (0.019) | 0.96 (0.021) | 1.20 (0.012) | 1.21 (0.0087) | 0.91 (0.010) |
| 5th | 2.13 (0.062) | 2.04 (0.042) | 0.87 (0.018) | 1.82 (0.037) | 1.81 (0.027) | 0.79 (0.016) |

(a)

| **annotation** | $\tau^*_{EAS}(C)$ (s.e.) | $\tau^*_{EUR}(C)$ (s.e.) | $\theta^*(C)$ (s.e.) |
|---|---|---|---|
| distance to nearest exon | 0.020 (0.021) | -0.03 (0.018) | -0.015 (0.020) |
| background selection statistic | 0.28 (0.027) | 0.25 (0.020) | 0.19 (0.021) |

(b)

Table S25: **Numerical S-LDXR results for distance to nearest exon annotation. a**) Heritability enrichment and enrichment of squared trans-ethnic genetic correlation ($\lambda^2(C)$) of the reversed distance to nearest exon annotation and the background selection statistic annotation. Standard errors are displayed in parentheses. **b**) Standardized annotation effect sizes of the distance to nearest exon annotation and the background selection statistic annotation.

| **pLI decile** | $h^2_{g,EAS}(C)$ **enrch.** | $h^2_{g,EUR}(C)$ **enrch.** | $\lambda^2(C)$ |
|---|---|---|---|
| 1st | 1.39 (0.035) | 1.38 (0.023) | 0.851 (0.018) |
| 2nd | 1.62 (0.045) | 1.56 (0.03) | 0.863 (0.018) |
| 3rd | 1.68 (0.045) | 1.57 (0.029) | 0.9 (0.018) |
| 4th | 1.62 (0.045) | 1.55 (0.03) | 0.887 (0.019) |
| 5th | 1.68 (0.048) | 1.65 (0.03) | 0.871 (0.018) |
| 6th | 1.62 (0.047) | 1.64 (0.031) | 0.866 (0.019) |
| 7th | 1.55 (0.044) | 1.55 (0.029) | 0.922 (0.02) |
| 8th | 1.83 (0.044) | 1.8 (0.03) | 0.873 (0.018) |
| 9th | 1.95 (0.044) | 1.92 (0.029) | 0.888 (0.016) |
| 10th | 1.91 (0.04) | 1.85 (0.023) | 0.895 (0.015) |

Table S26: **Numerical S-LDXR results for deciles of probability of loss-of-function intolerance (pLI) annotations.**

| $\tau$ | $\sigma^2_\Delta$ | $h^2_{g1}(A)$ **enrch.** | $h^2_{g1}(B)$ **enrch.** | $h^2_{g2}(A)$ **enrch.** | $h^2_{g2}(B)$ **enrch.** | $\lambda^2(A)$ | $\lambda^2(B)$ |
|---|---|---|---|---|---|---|---|
| 0 | 0 | 1.0 (0.002) | 1.0 (0.02) | 1.0 (0.002) | 1.0 (0.02) | 1.0 (0.0) | 1.0 (0.0) |
| 0 | 0.2 | 1.0 (0.002) | 1.0 (0.02) | 1.0 (0.002) | 1.0 (0.02) | 1.0 (0.0) | 1.0 (0.0) |
| 0 | 0.4 | 1.0 (0.002) | 1.0 (0.02) | 1.0 (0.002) | 1.0 (0.02) | 1.0 (0.0) | 1.0 (0.0) |
| 0 | 0.6 | 1.0 (0.002) | 1.0 (0.02) | 1.0 (0.002) | 1.0 (0.02) | 1.0 (0.0) | 1.0 (0.0) |
| 0 | 0.8 | 1.0 (0.002) | 1.0 (0.02) | 1.0 (0.002) | 1.0 (0.02) | 1.0 (0.0) | 1.0 (0.0) |
| 0 | 1 | 1.0 (0.002) | 1.0 (0.02) | 1.0 (0.002) | 1.0 (0.02) | 1.0 (0.0) | 1.0 (0.0) |
| 0.2 | 0 | 0.87 (0.004) | 2.2 (0.03) | 0.87 (0.004) | 2.2 (0.03) | 1.0 (0.0) | 1.0 (0.0) |
| 0.2 | 0.2 | 0.88 (0.004) | 2.1 (0.03) | 0.88 (0.004) | 2.1 (0.03) | 1.01 (0.0008) | 0.96 (0.003) |
| 0.2 | 0.4 | 0.89 (0.004) | 2.1 (0.03) | 0.89 (0.004) | 2.1 (0.03) | 1.03 (0.002) | 0.88 (0.007) |
| 0.2 | 0.6 | 0.89 (0.004) | 2.01 (0.03) | 0.89 (0.004) | 2.01 (0.03) | 1.05 (0.002) | 0.82 (0.008) |
| 0.2 | 0.8 | 0.89 (0.004) | 2.0 (0.04) | 0.89 (0.004) | 2.0 (0.04) | 1.06 (0.003) | 0.76 (0.009) |
| 0.2 | 1 | 0.89 (0.004) | 2.0 (0.04) | 0.89 (0.004) | 2.0 (0.04) | 1.07 (0.003) | 0.72 (0.01) |
| 0.4 | 0 | 0.65 (0.006) | 4.12 (0.05) | 0.65 (0.006) | 4.12 (0.05) | 1.0 (0.0) | 1.0 (0.0) |
| 0.4 | 0.2 | 0.66 (0.007) | 4.08 (0.05) | 0.66 (0.007) | 4.08 (0.05) | 1.04 (0.002) | 0.94 (0.003) |
| 0.4 | 0.4 | 0.66 (0.007) | 4.07 (0.06) | 0.66 (0.007) | 4.07 (0.06) | 1.10 (0.004) | 0.87 (0.006) |
| 0.4 | 0.6 | 0.66 (0.007) | 4.07 (0.06) | 0.66 (0.007) | 4.07 (0.06) | 1.14 (0.006) | 0.81 (0.008) |
| 0.4 | 0.8 | 0.66 (0.007) | 4.09 (0.06) | 0.66 (0.007) | 4.09 (0.06) | 1.18 (0.007) | 0.76 (0.009) |
| 0.4 | 1 | 0.65 (0.007) | 4.11 (0.06) | 0.65 (0.007) | 4.11 (0.06) | 1.21 (0.008) | 0.73 (0.01) |
| 0.6 | 0 | 0.40 (0.006) | 6.38 (0.07) | 0.40 (0.006) | 6.38 (0.07) | 1.0 (0.0) | 1.0 (0.0) |
| 0.6 | 0.2 | 0.40 (0.007) | 6.44 (0.07) | 0.40 (0.007) | 6.44 (0.07) | 1.12 (0.009) | 0.94 (0.003) |
| 0.6 | 0.4 | 0.39 (0.007) | 6.52 (0.07) | 0.39 (0.007) | 6.52 (0.07) | 1.24 (0.009) | 0.88 (0.005) |
| 0.6 | 0.6 | 0.38 (0.007) | 6.61 (0.08) | 0.38 (0.007) | 6.61 (0.08) | 1.35 (0.01) | 0.84 (0.006) |
| 0.6 | 0.8 | 0.37 (0.007) | 6.70 (0.08) | 0.37 (0.007) | 6.70 (0.08) | 1.44 (0.02) | 0.81 (0.008) |
| 0.6 | 1 | 0.36 (0.007) | 6.77 (0.09) | 0.36 (0.007) | 6.77 (0.09) | 1.52 (0.02) | 0.79 (0.009) |
| 0.8 | 0 | 0.21 (0.004) | 8.16 (0.09) | 0.21 (0.004) | 8.16 (0.09) | 1.0 (0.0) | 1.0 (0.0) |
| 0.8 | 0.2 | 0.19 (0.004) | 8.30 (0.09) | 0.19 (0.004) | 8.30 (0.09) | 1.23 (0.009) | 0.95 (0.002) |
| 0.8 | 0.4 | 0.18 (0.004) | 8.42 (0.10) | 0.18 (0.004) | 8.42 (0.10) | 1.47 (0.02) | 0.92 (0.003) |
| 0.8 | 0.6 | 0.16 (0.004) | 8.52 (0.10) | 0.16 (0.004) | 8.52 (0.10) | 1.67 (0.03) | 0.90 (0.004) |
| 0.8 | 0.8 | 0.15 (0.004) | 8.61 (0.10) | 0.15 (0.004) | 8.61 (0.10) | 1.84 (0.04) | 0.89 (0.005) |
| 0.8 | 1 | 0.15 (0.004) | 8.69 (0.11) | 0.15 (0.004) | 8.69 (0.11) | 1.99 (0.04) | 0.88 (0.005) |
| 1 | 0 | 0.092 (0.002) | 9.18 (0.11) | 0.092 (0.002) | 9.18 (0.11) | 1.0 (0.0) | 1.0 (0.0) |
| 1 | 0.2 | 0.078 (0.002) | 9.30 (0.12) | 0.078 (0.002) | 9.30 (0.12) | 1.39 (0.02) | 0.97 (0.002) |
| 1 | 0.4 | 0.067 (0.002) | 9.39 (0.12) | 0.067 (0.002) | 9.39 (0.12) | 1.75 (0.03) | 0.96 (0.002) |
| 1 | 0.6 | 0.060 (0.002) | 9.46 (0.13) | 0.060 (0.002) | 9.46 (0.13) | 2.06 (0.05) | 0.95 (0.002) |
| 1 | 0.8 | 0.054 (0.002) | 9.51 (0.12) | 0.054 (0.002) | 9.51 (0.12) | 2.32 (0.07) | 0.95 (0.003) |
| 1 | 1 | 0.050 (0.002) | 9.55 (0.12) | 0.050 (0.002) | 9.55 (0.12) | 2.54 (0.08) | 0.95 (0.003) |

Table S27: **Numerical evolutionary modeling results using 2-population extension of Eyre-Walker model.** Standard errors of the mean are reported in parenthesis. Here, $A$ refers to the set of SNPs with $\bar{s} = -10^{-5}$, and $B$ the set of SNPs with $\bar{s} = -10^{-4}$. We use negative $s$ to denote deleteriousness, following convention of previous works.[11,12] However, positive $s$ (i.e. beneficial mutations) may also be plausible.

216 # Supplementary figures

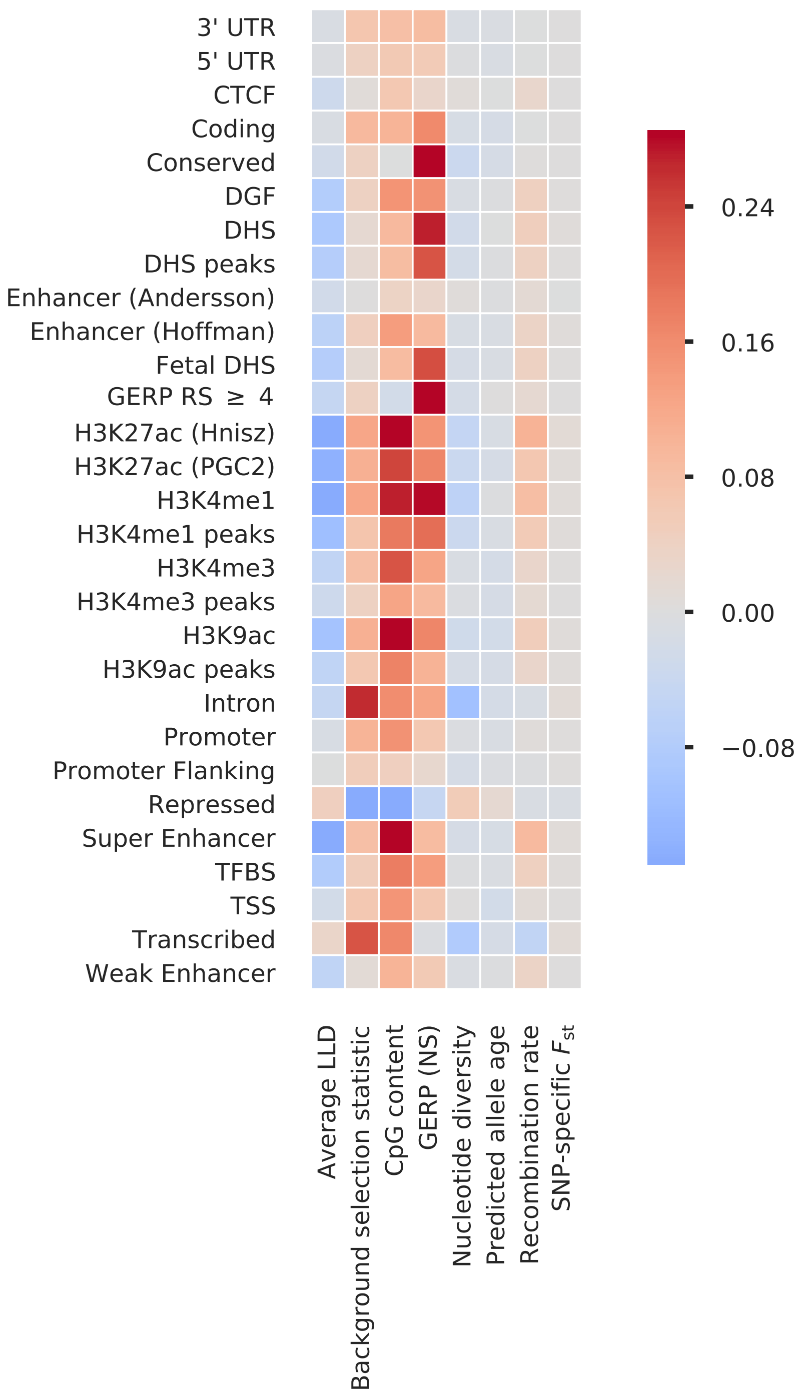


Figure S1: **Correlation between functional annotations and continuous-valued annotations.** The correlations were computed across SNPs with minor allele frequency > 5% in both populations.

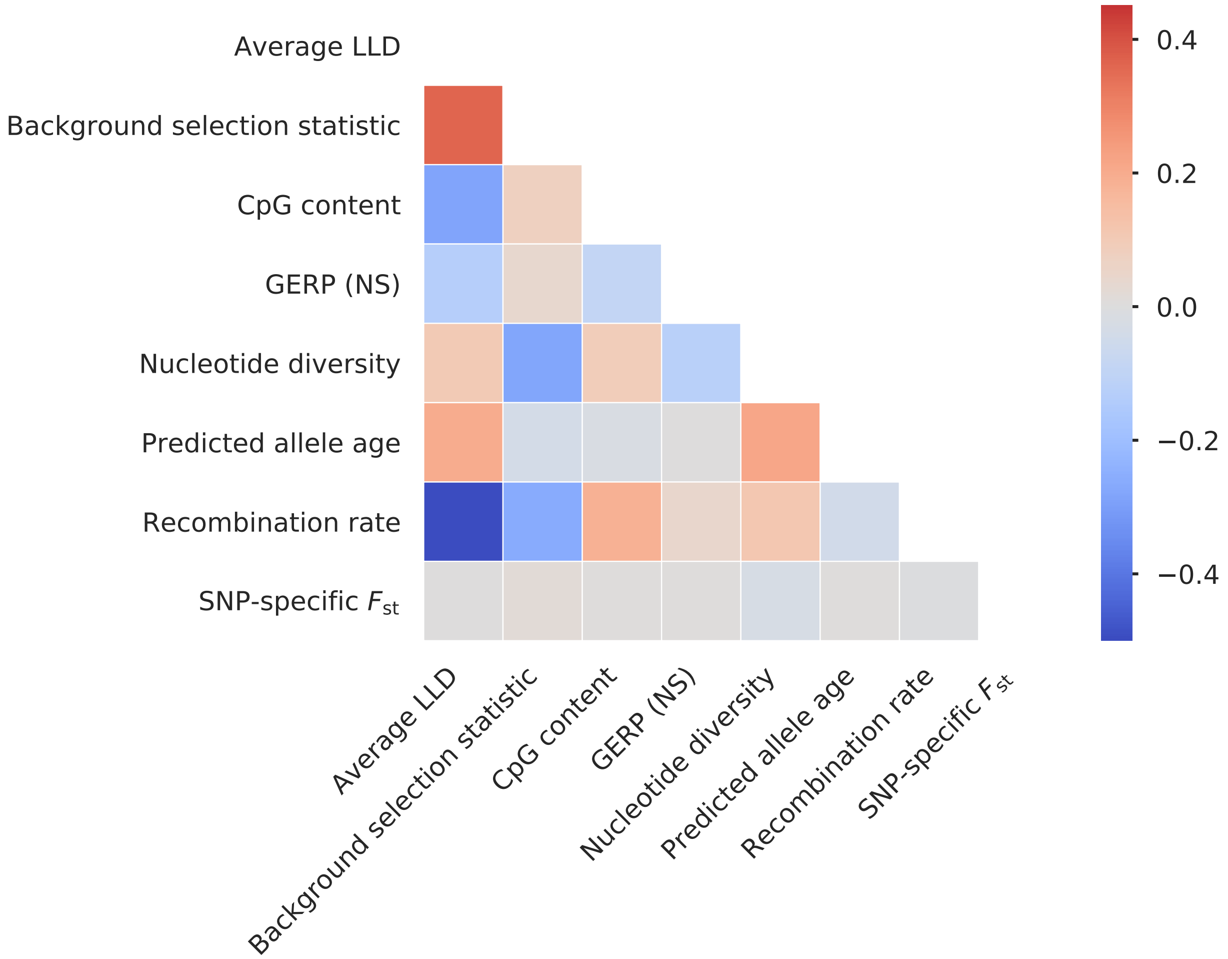


Figure S2: **Correlation between continuous-valued annotations.** The correlations were computed across SNPs with minor allele frequency > 5% in both populations.

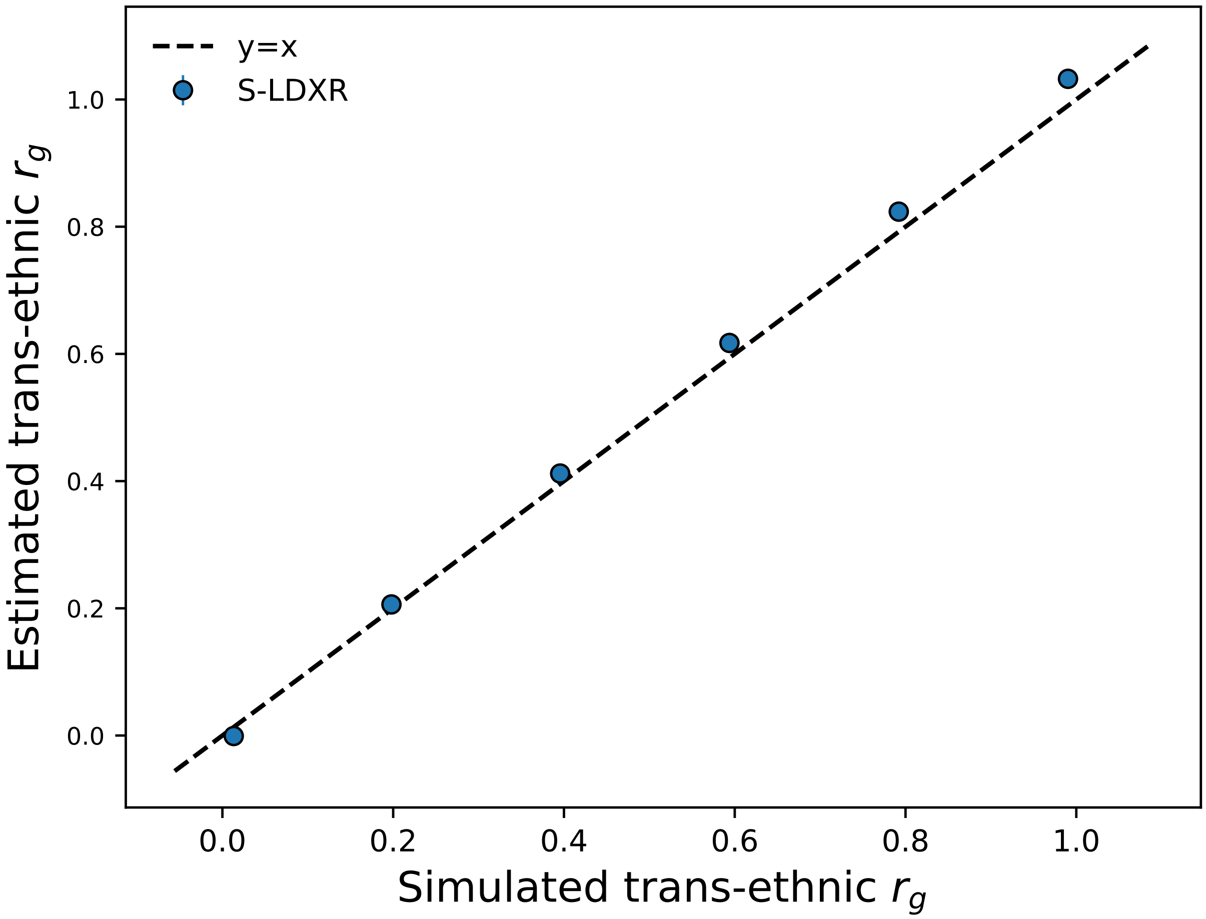


Figure S3: **Accuracy of S-LDXR in estimating genome-wide trans-ethnic genetic correlation.** Here, 10% of SNPs are randomly selected to be causal. Mean and standard errors were obtained across 1,000 simulations. Error bars represent 1.96 times the standard error on both sides.

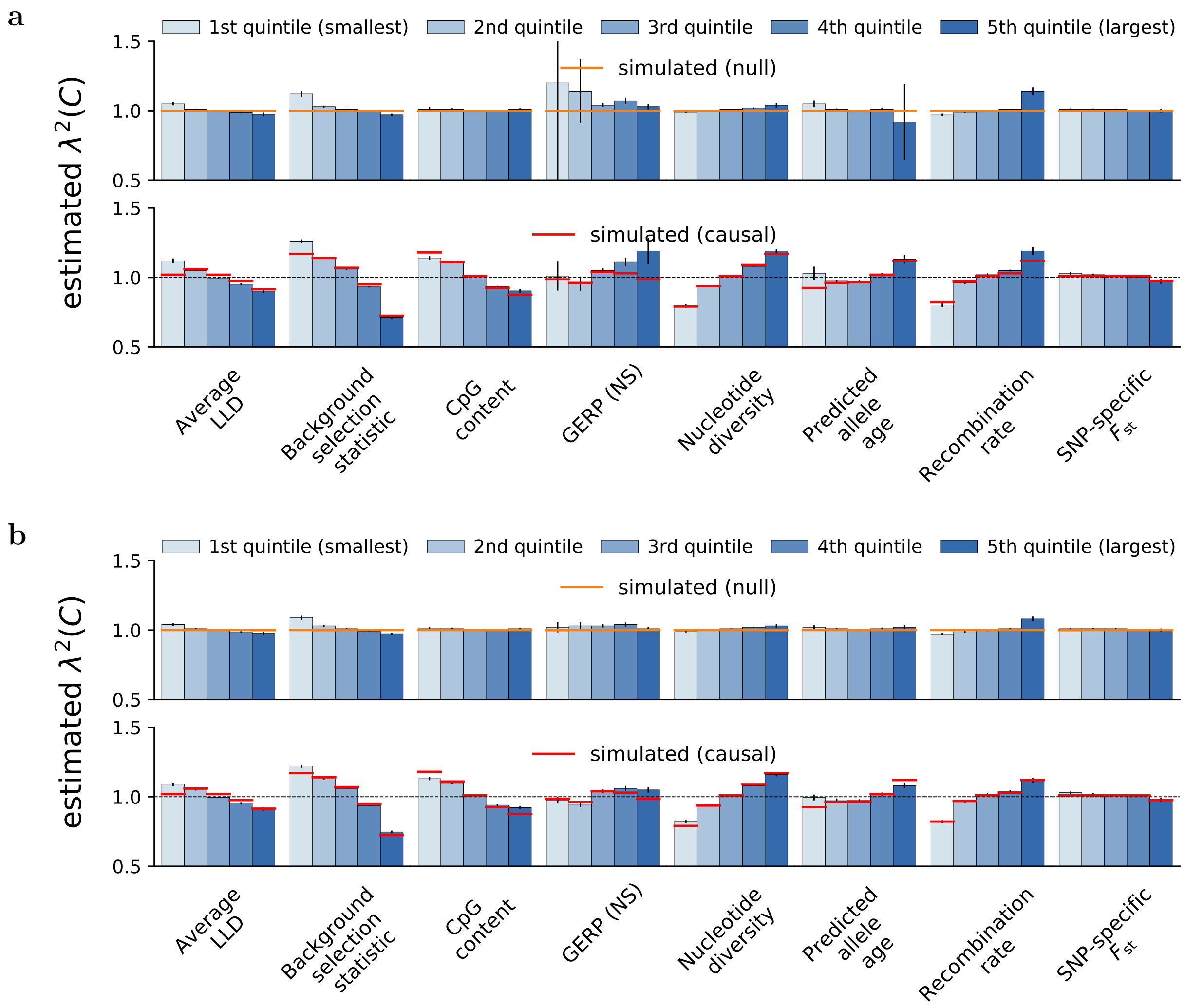


Figure S4: (continued on next page)

**c**

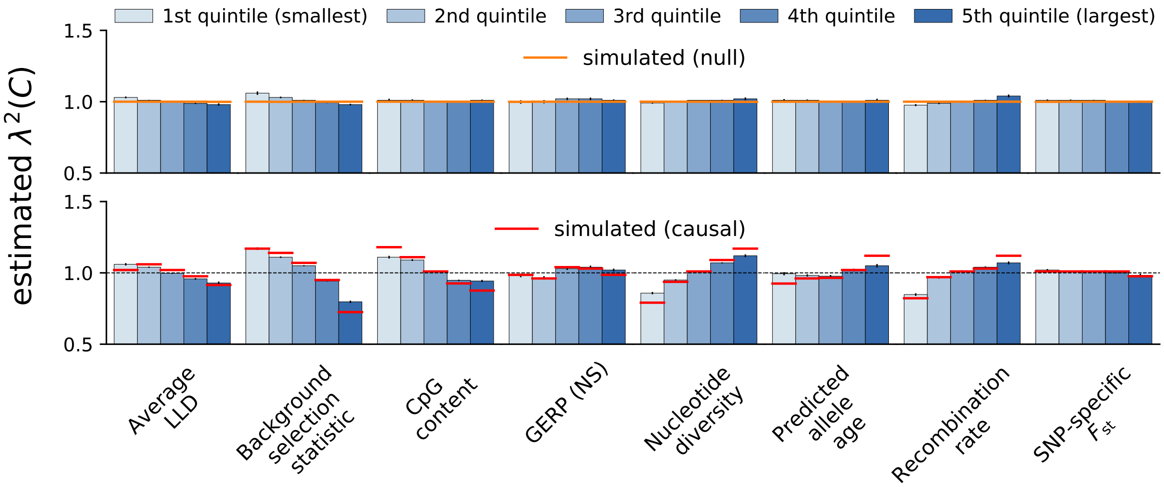


**d**

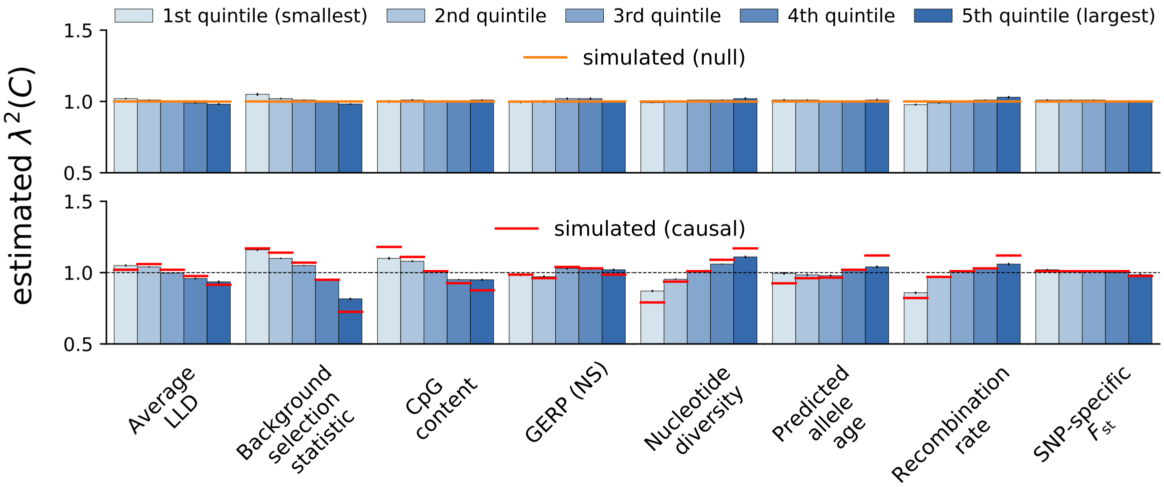


Figure S4: **Accuracy of S-LDXR in estimating enrichment of stratified squared trans-ethnic genetic correlation, $\lambda^2(C)$, of quintiles of continuous-valued annotations.** Here, 10% of SNPs were randomly selected to be causal. Shrinkage level, $\alpha$, was set to 0.0 in **a**, 0.25 in **b**, 0.75 in **c**, and 1.0 in **d**. Mean and standard errors were obtained across 1,000 simulations. Error bars represent 1.96 times the standard error on both sides.

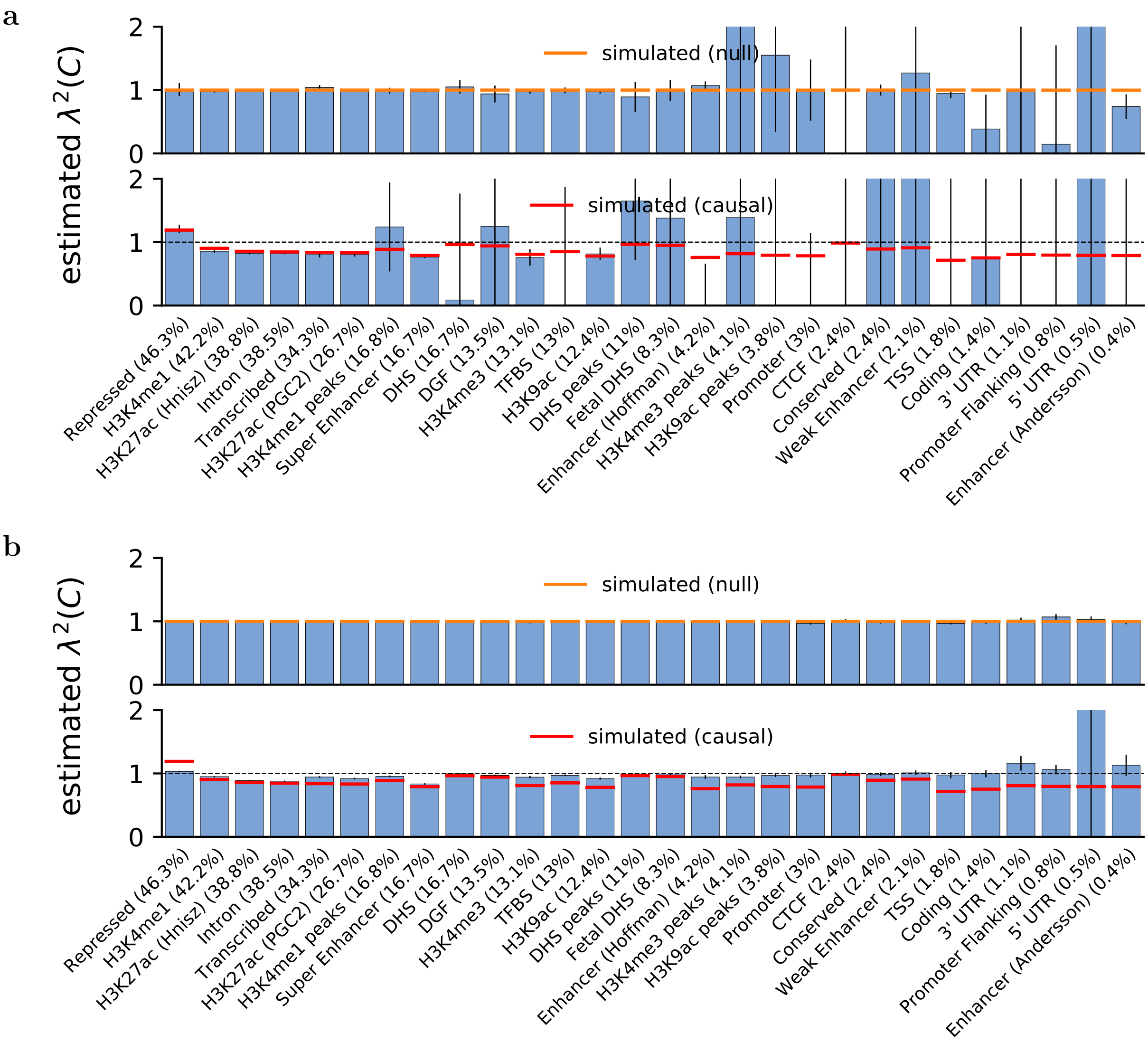


Figure S5: (continued on next page)

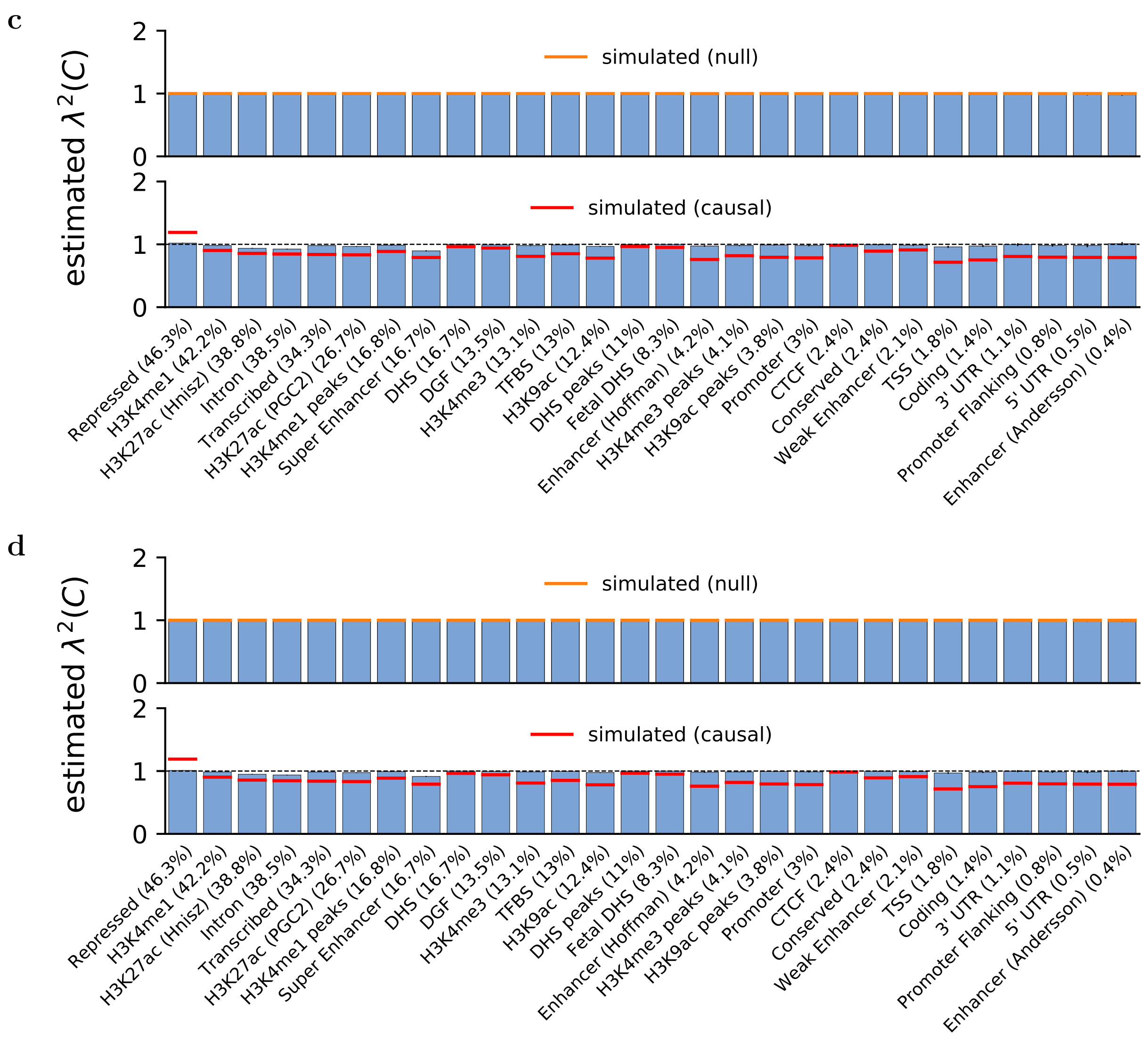


Figure S5: **Accuracy of S-LDXR in estimating enrichment of stratified squared trans-ethnic genetic correlation, $\lambda^2(C)$, of functional annotations.** Here, 10% of SNPs were randomly selected to be causal. Shrinkage level, $\alpha$, was set to 0.0 in **a**, 0.25 in **b**, 0.75 in **c**, and 1.0 in **d**. Mean and standard errors were obtained across 1,000 simulations. Error bars represent 1.96 times the standard error on both sides.

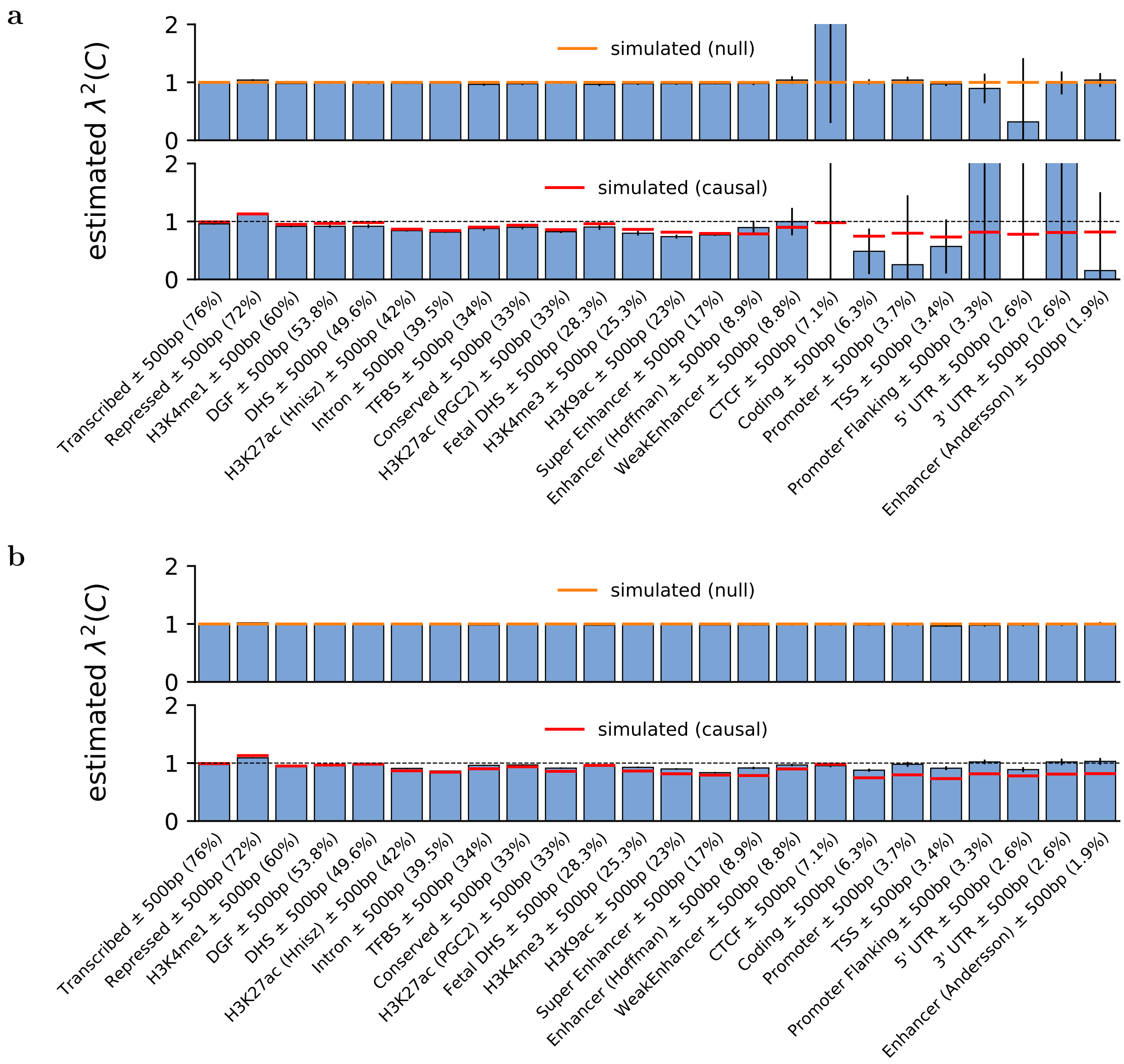


Figure S6: (continued on next page)

**c**

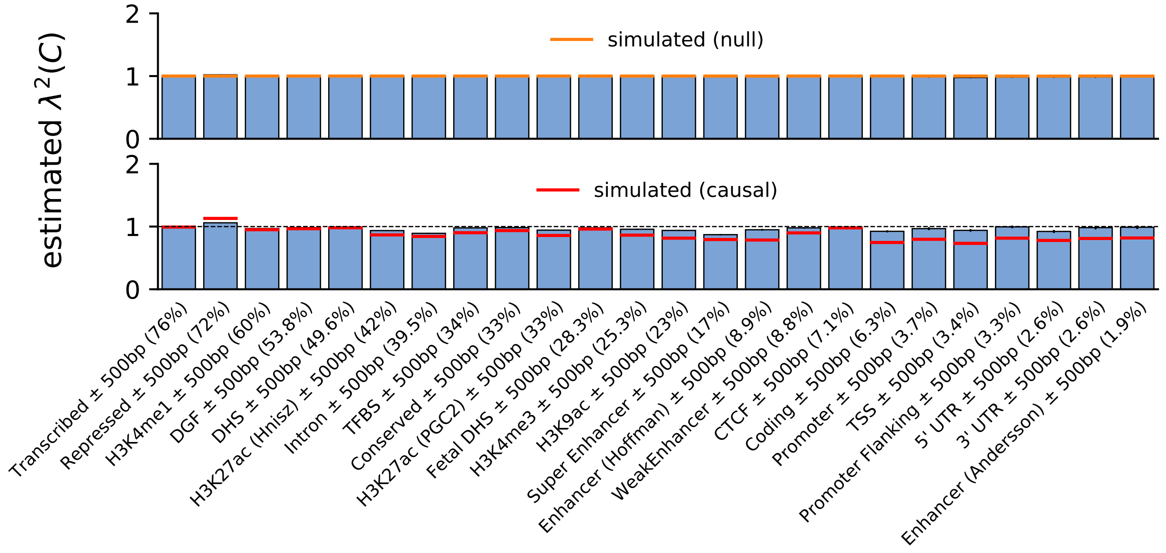


**d**

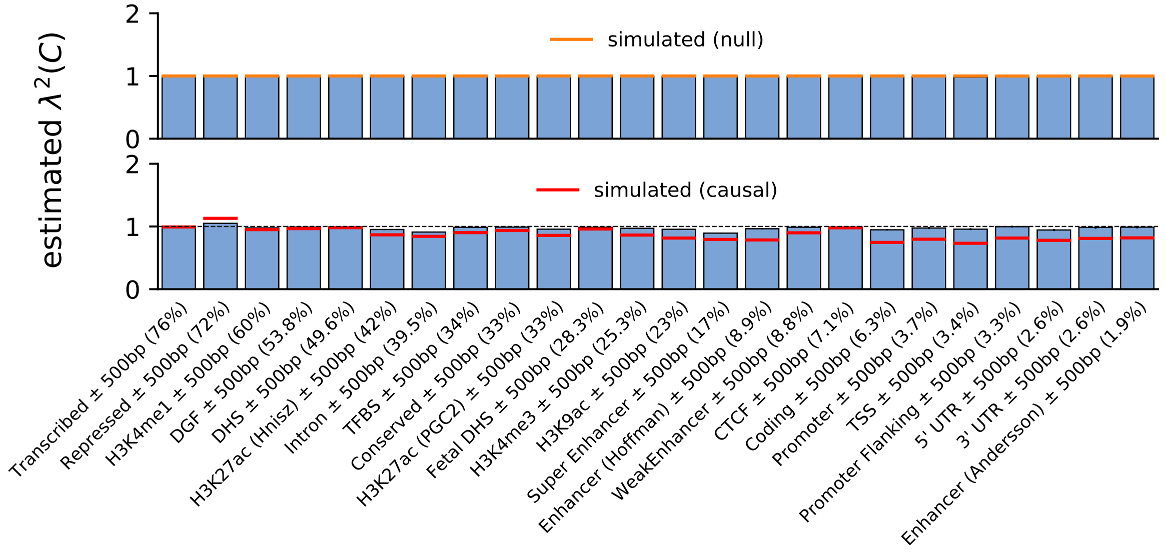


Figure S6: (continued on next page)

**e**

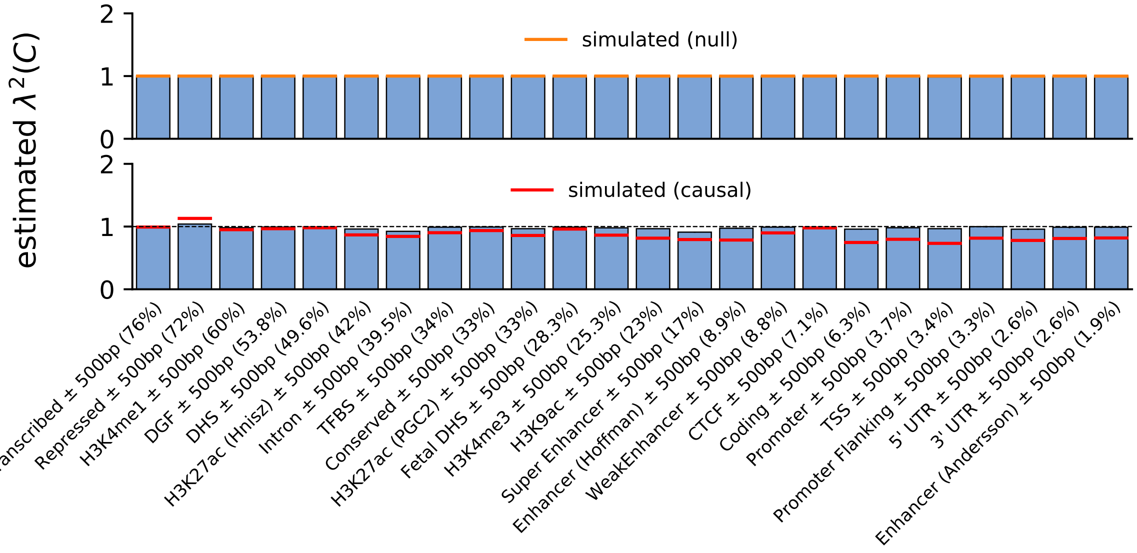


Figure S6: **Accuracy of S-LDXR in estimating enrichment of stratified squared trans-ethnic genetic correlation, $\lambda^2(C)$, of 500-base-pair extended functional annotations.** Here, 10% of SNPs were randomly selected to be causal. Shrinkage level, $\alpha$, was set to 0.0 in **a**, 0.25 in **b**, 0.5 in **c**, 0.75 in **d**, and 1.0 in **e**. Mean and standard errors were obtained across 1,000 simulations. Error bars represent 1.96 times the standard error on both sides.

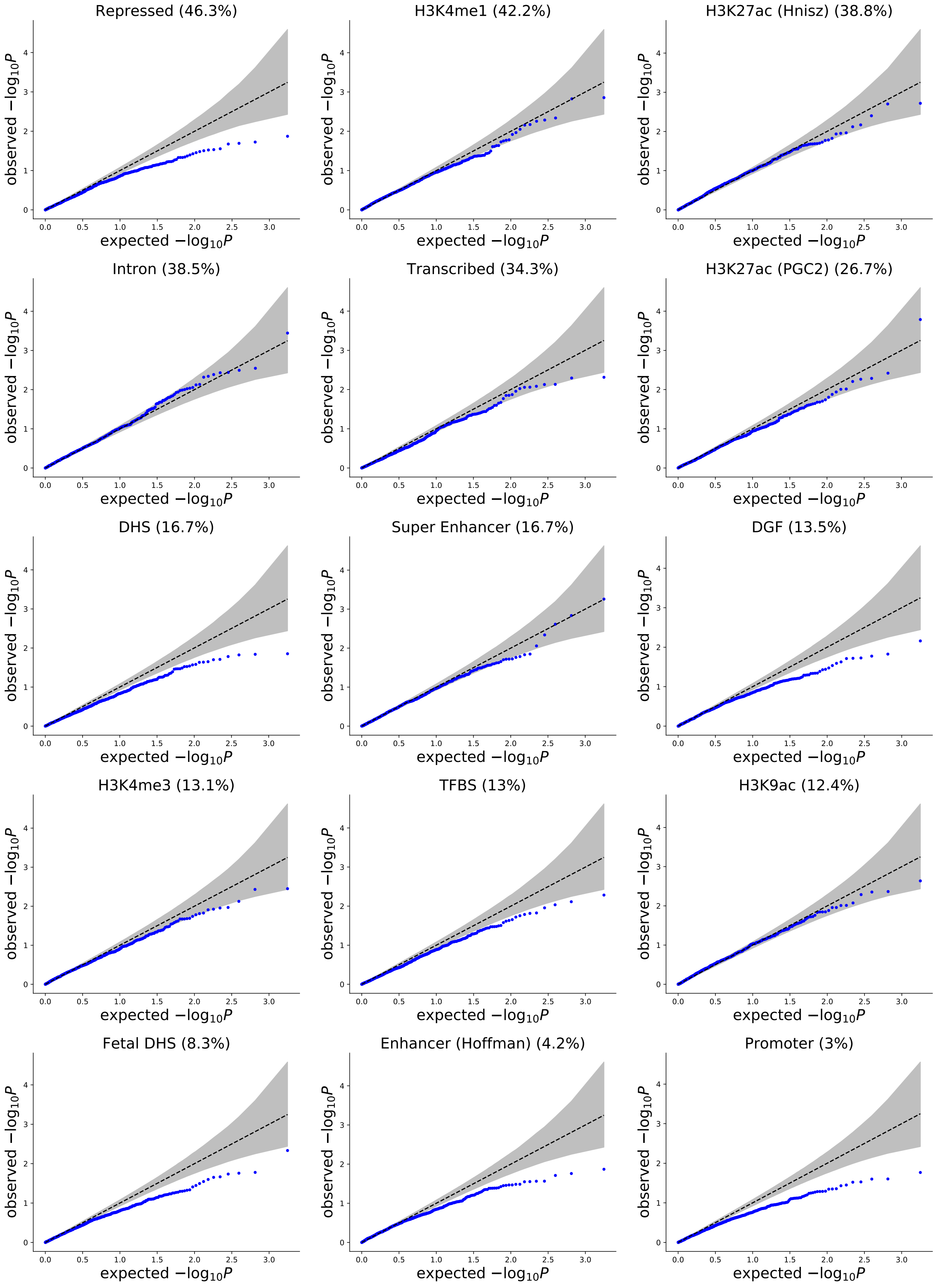


Figure S7: (continued on next page)

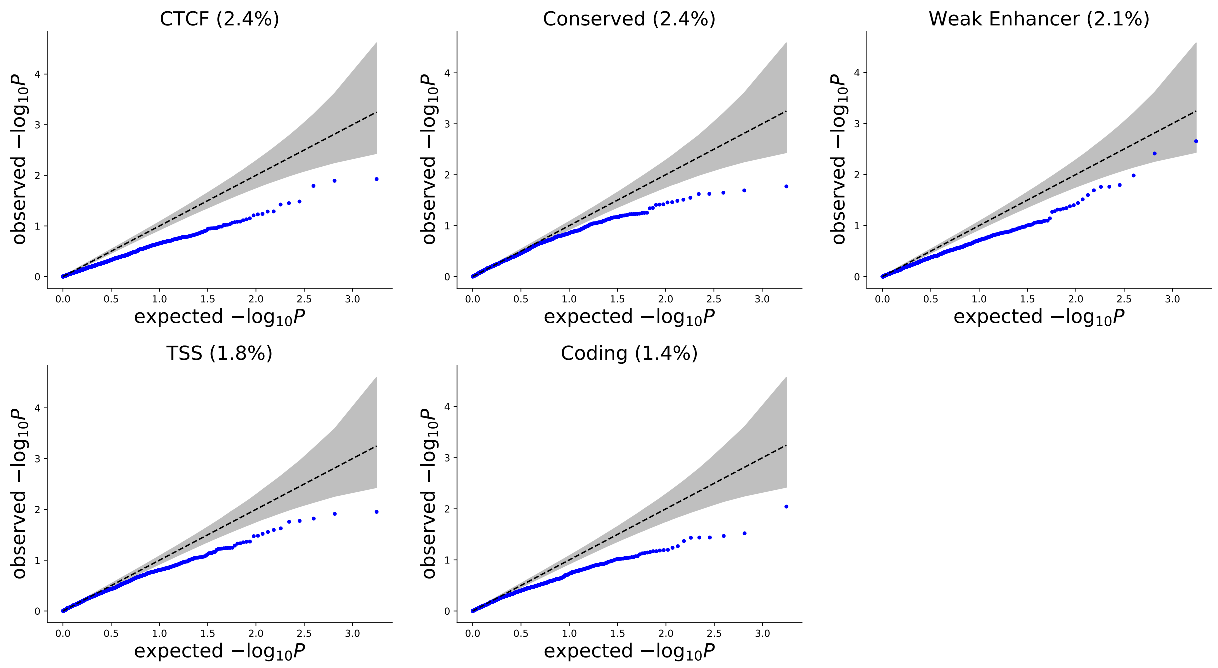


Figure S7: **Q-Q plot for S-LDXR p-values testing enrichment of squared trans-ethnic genetic correlation of 20 main functional annotations in 1,000 null simulations.** 10% of SNPs were randomly selected to be causal. S-LDXR was applied with the baseline-LD-X model annotations. The shrinkage level, $\alpha$, was set to 0.5. Shaded area represent the 95% confidence interval. Size of the annotation (proportion of SNPs) is shown in parentheses.

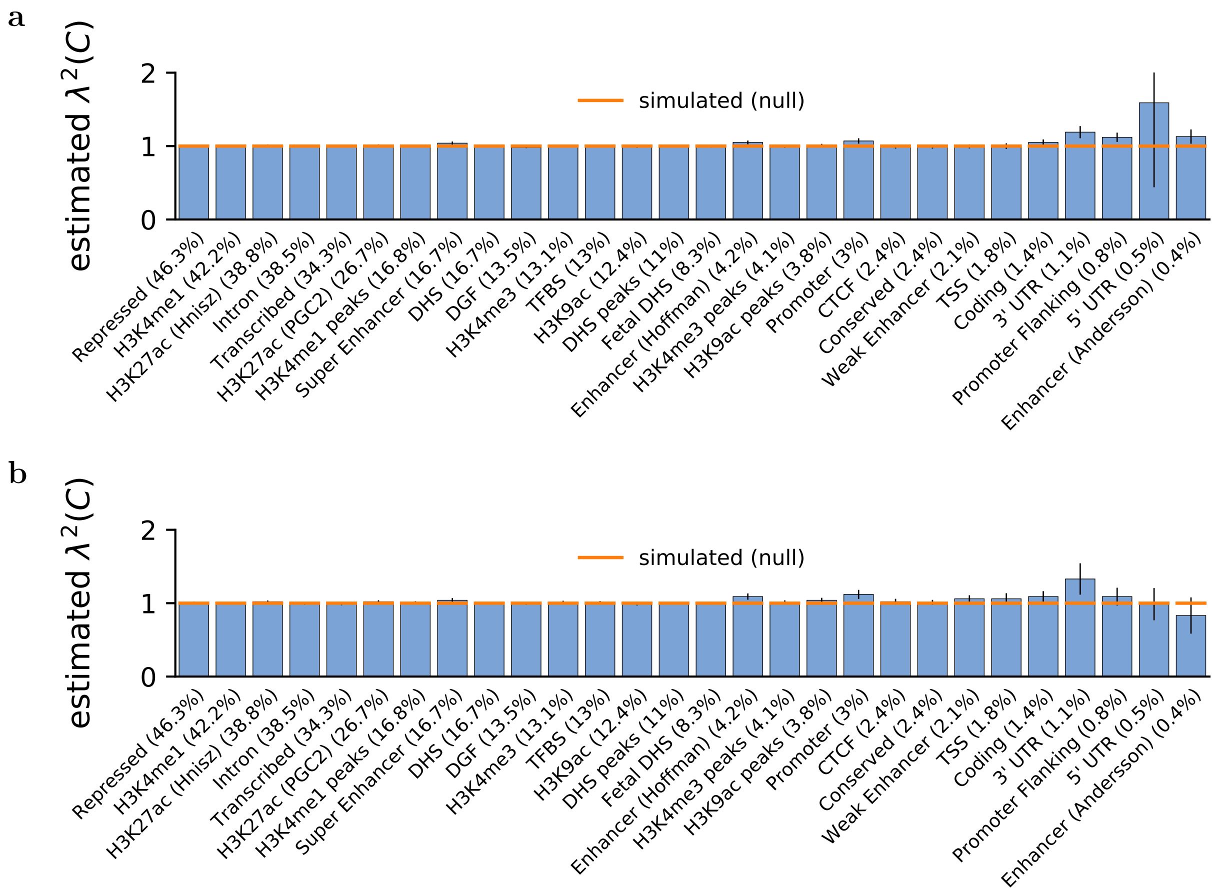


Figure S8: **Accuracy of S-LDXR in estimating enrichment of stratified squared trans-ethnic genetic correlation, $\lambda^2(C)$, of functional annotations in simulations with annotation-dependent MAF-dependent genetic architectures.** 10% of SNPs were randomly selected to be causal. Shrinkage level, $\alpha$, was set to 0.5. **a**) S-LDXR was applied with the baseline-LD-X model annotations. **b**) S-LDXR was applied with the baseline-LD-X model annotations and 5 MAF bin annotations. Mean and standard errors were obtained across 1,000 simulations. Error bars represent 1.96 times the standard error on both sides.

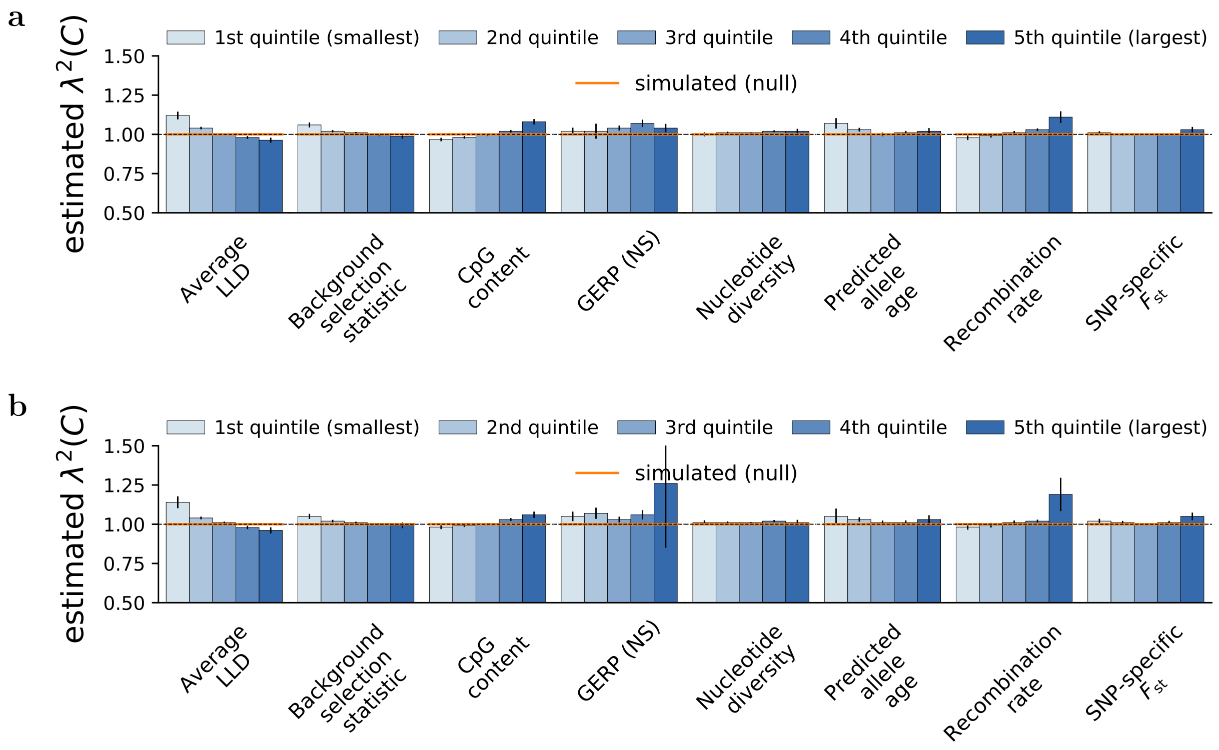


Figure S9: **Accuracy of S-LDXR in estimating enrichment of stratified squared trans-ethnic genetic correlation, $\lambda^2(C)$, of quintiles of continuous-valued annotations in simulations with annotation-dependent MAF-dependent genetic architectures.** 10% of SNPs were randomly selected to be causal. Shrinkage level, $\alpha$, was set to 0.5. **a**) S-LDXR was applied with the baseline-LD-X model annotations. **b**) S-LDXR was applied with the baseline-LD-X model annotations and 5 MAF bin annotations. Mean and standard errors were obtained across 1,000 simulations. Error bars represent 1.96 times the standard error on both sides.

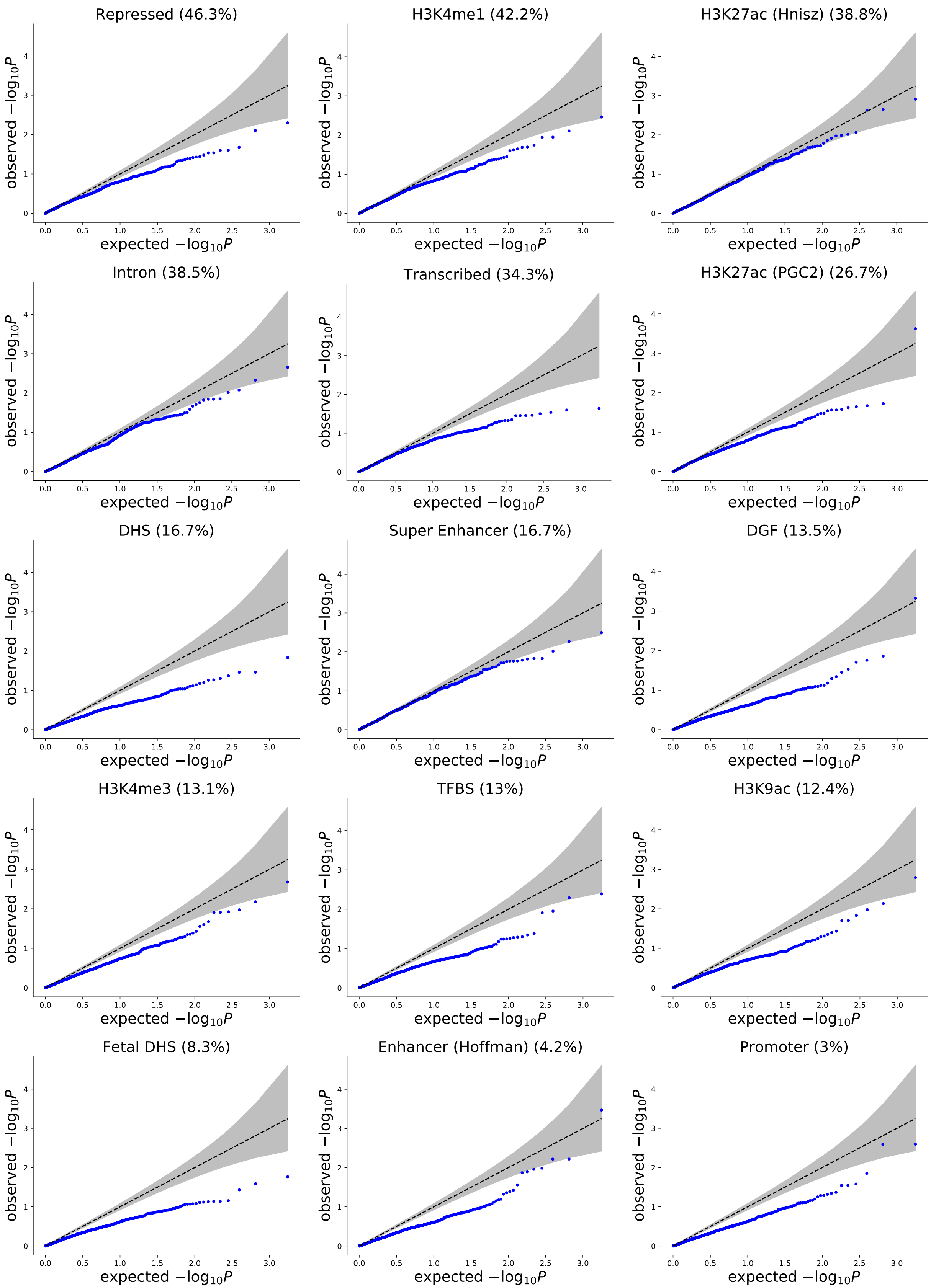


Figure S10: (continued on next page)

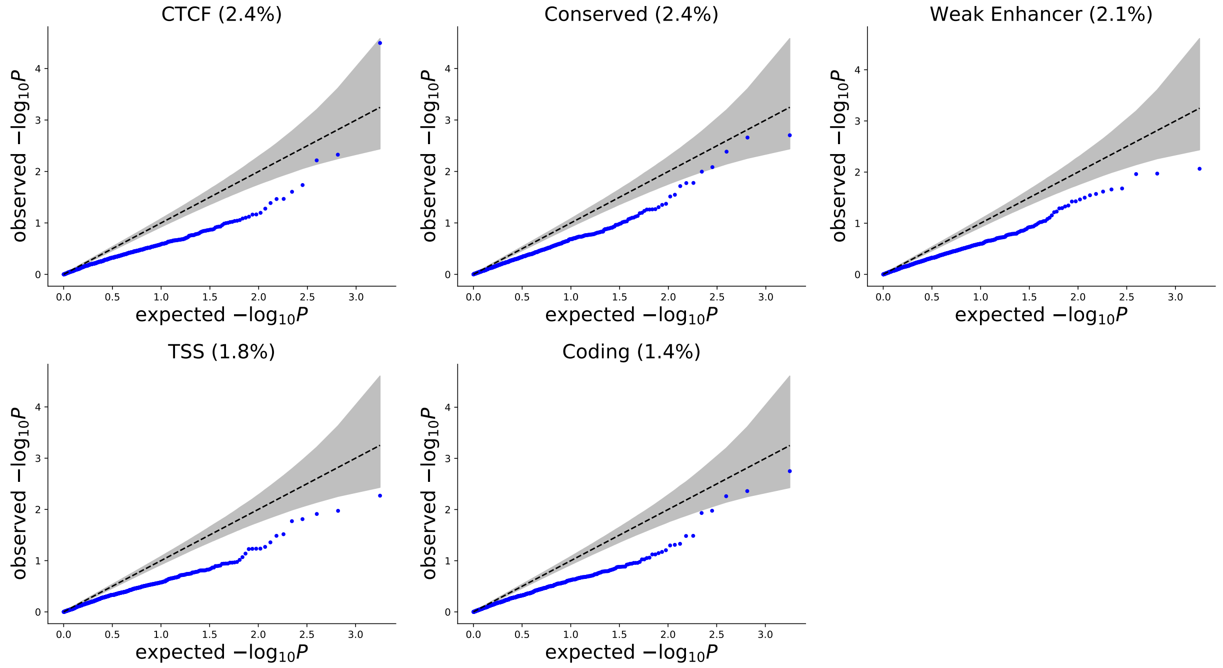


Figure S10: **Q-Q plot for S-LDXR p-values testing enrichment of squared trans-ethnic genetic correlation of 20 main functional annotations in 1,000 null simulations with annotation-dependent MAF-dependent genetic architectures.** 10% of SNPs were randomly selected to be causal. S-LDXR was applied with the baseline-LD-X model annotations. The shrinkage level, $\alpha$, was set to 0.5. Shaded area represent the 95% confidence interval. Size of the annotation (proportion of SNPs) is shown in parenthesis.

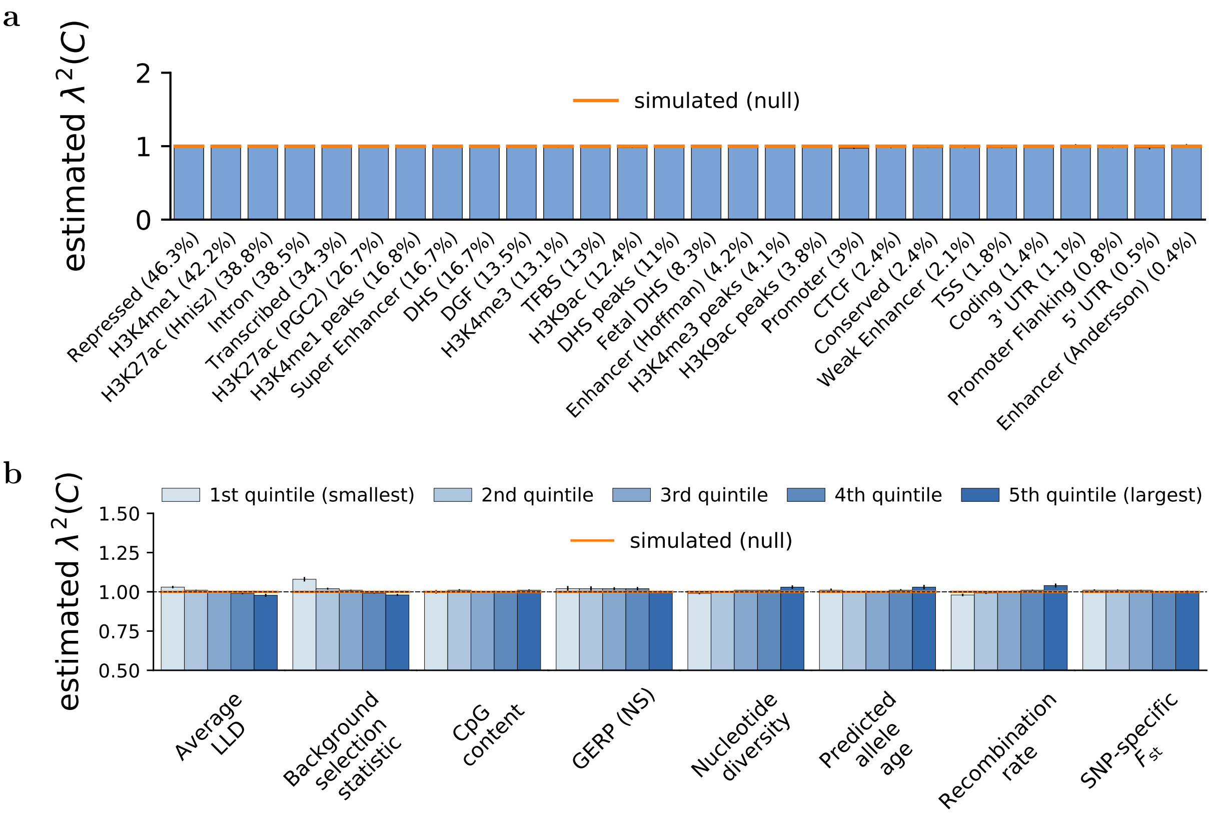


Figure S11: **Accuracy of S-LDXR in estimating enrichment of stratified squared trans-ethnic genetic correlation, $\lambda^2(C)$, when causal variants differ across the two populations.** 10% of SNPs were randomly selected to be causal. Shrinkage level, $\alpha$, was set to 0.5. **a**) Estimates of $\lambda^2(C)$ for binary functional annotations. **b**) Estimates of $\lambda^2(C)$ for quintiles of continuous-valued annotations. Mean and standard errors were obtained across 1,000 simulations. Error bars represent 1.96 times the standard error on both sides.

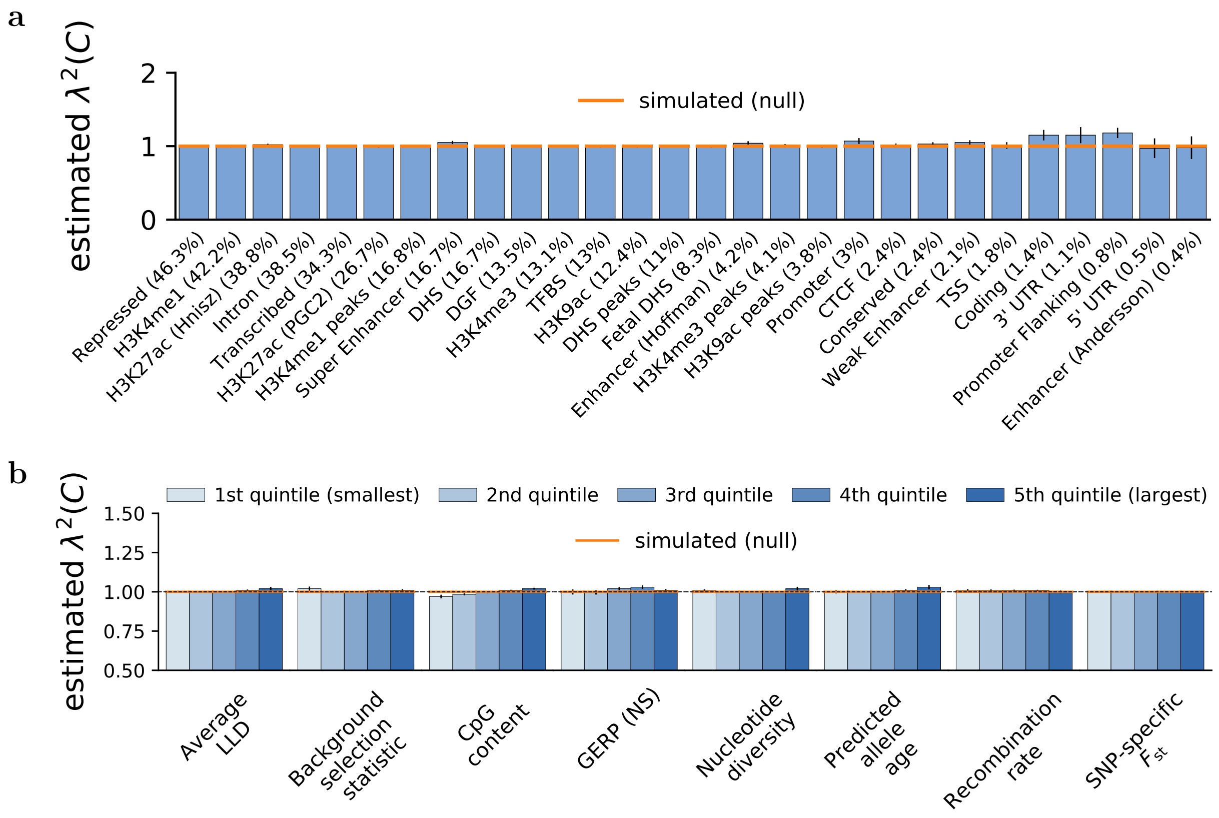


Figure S12: **Accuracy of S-LDXR in estimating enrichment of stratified squared trans-ethnic genetic correlation, $\lambda^2(C)$, in simulations under the baseline-LD-X model using all simulated GWAS samples and half (250) the default reference panel sample size.** 10% of SNPs were randomly selected to be causal. Shrinkage level, $\alpha$, was set to 0.5. **a**) Estimates of $\lambda^2(C)$ for binary functional annotations. **b**) Estimates of $\lambda^2(C)$ for quintiles of continuous-valued annotations. Mean and standard errors were obtained across 1,000 simulations. Error bars represent 1.96 times the standard error on both sides. Numerical results are reported in Table S13a.

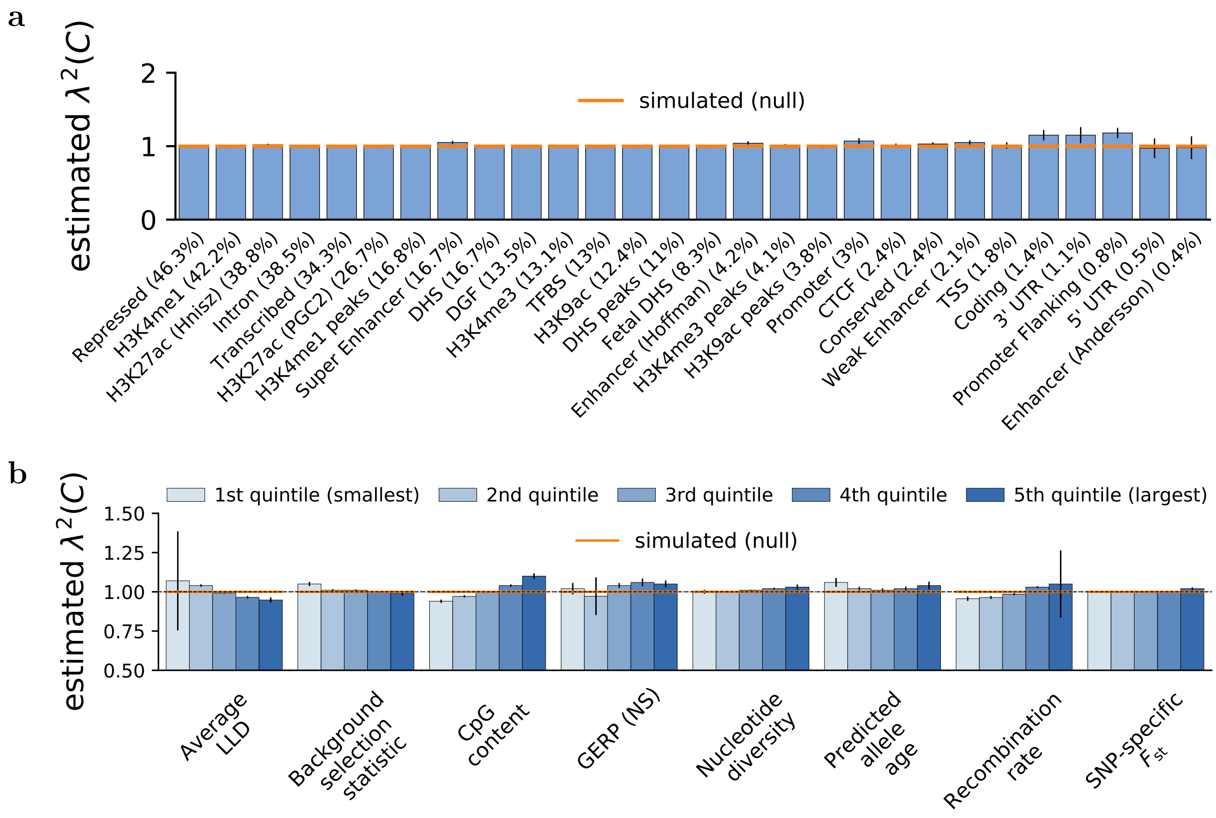


Figure S13: **Accuracy of S-LDXR in estimating enrichment of stratified squared trans-ethnic genetic correlation, $\lambda^2(C)$, in simulations with annotation-dependent MAF-dependent genetic architectures using all simulated GWAS samples and half (250) the default reference panel sample size.** 10% of SNPs were randomly selected to be causal. Shrinkage level, $\alpha$, was set to 0.5. **a**) Estimates of $\lambda^2(C)$ for binary functional annotations. **b**) Estimates of $\lambda^2(C)$ for quintiles of continuous-valued annotations. Mean and standard errors were obtained across 1,000 simulations. Error bars represent 1.96 times the standard error on both sides. Numerical results are reported in Table S13b.

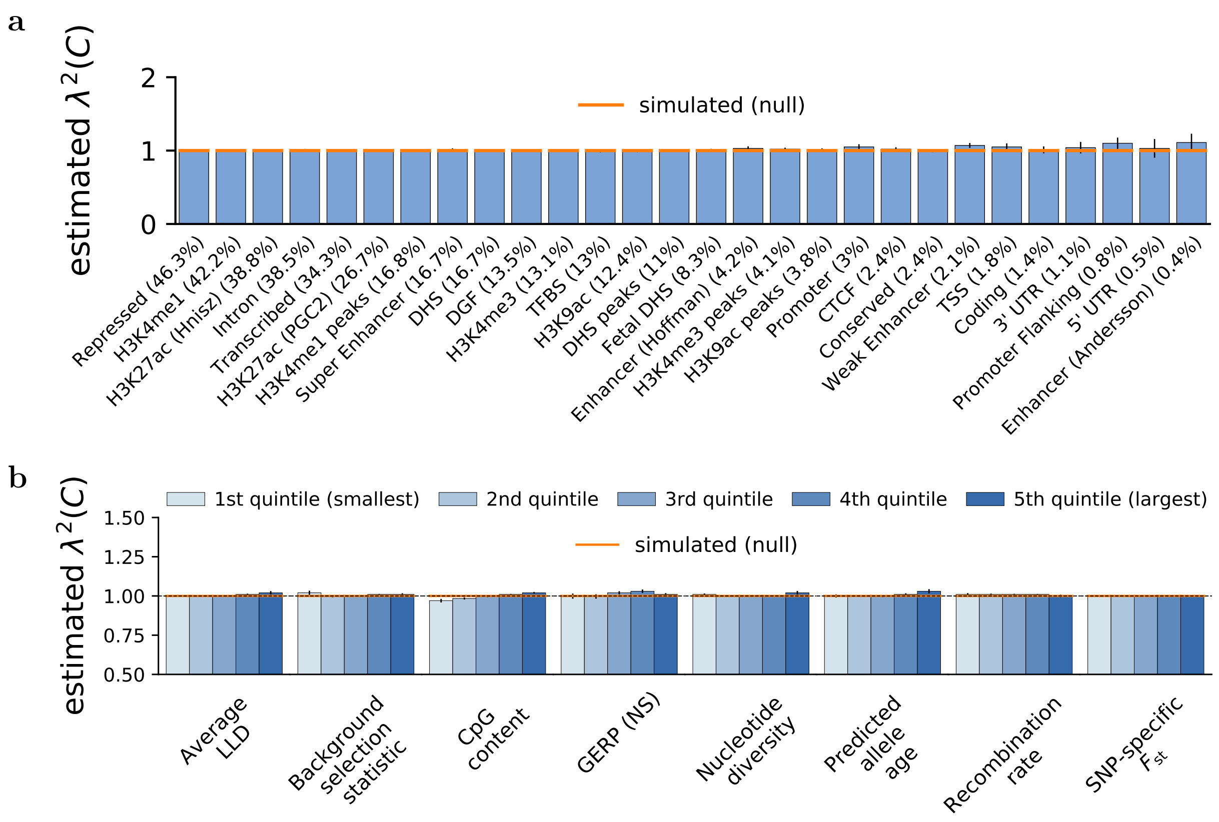


Figure S14: **Accuracy of S-LDXR in estimating enrichment of stratified squared trans-ethnic genetic correlation, $\lambda^2(C)$, in simulations under the baseline-LD-X model using all simulated GWAS samples and twice (1,000) the default reference panel sample size.** 10% of SNPs were randomly selected to be causal. Shrinkage level, $\alpha$, was set to 0.5. **a**) Estimates of $\lambda^2(C)$ for binary functional annotations. **b**) Estimates of $\lambda^2(C)$ for quintiles of continuous-valued annotations. Mean and standard errors were obtained across 1,000 simulations. Error bars represent 1.96 times the standard error on both sides. Numerical results are reported in Table S14a.

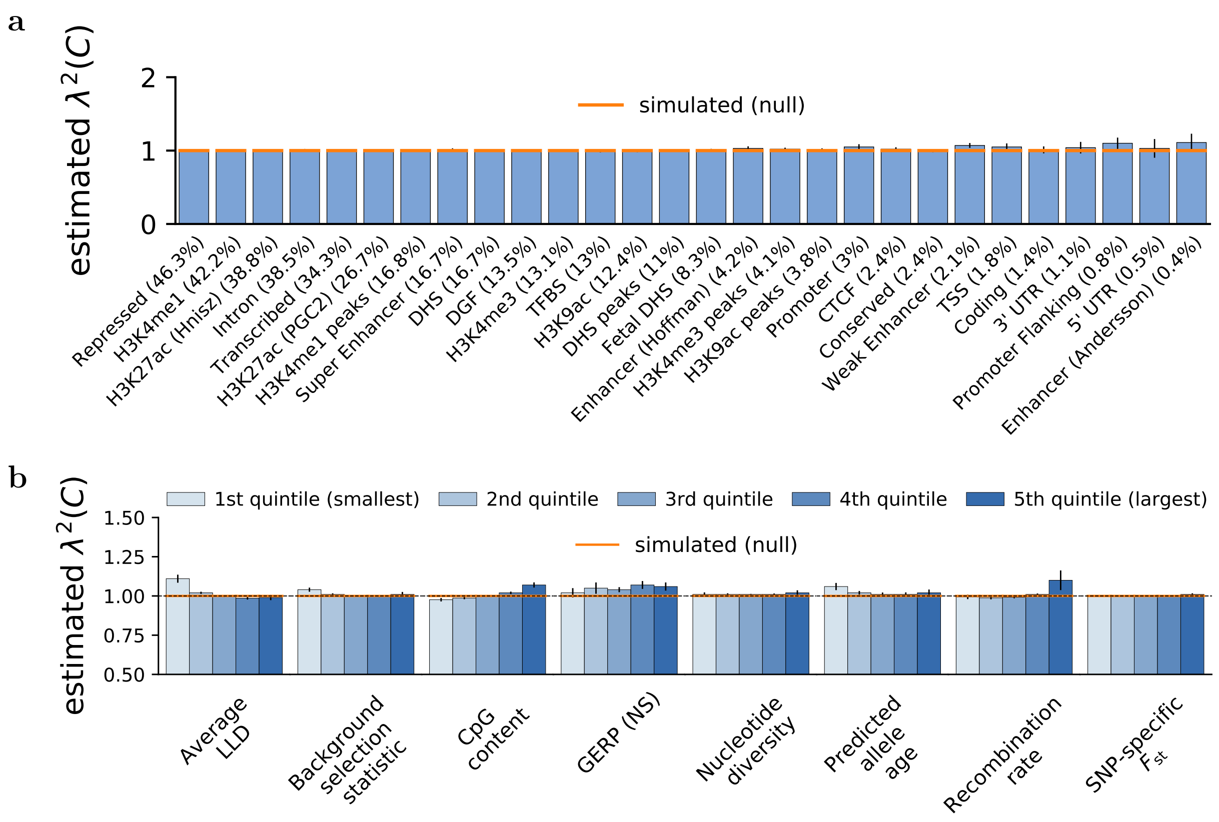


Figure S15: **Accuracy of S-LDXR in estimating enrichment of stratified squared trans-ethnic genetic correlation, $\lambda^2(C)$, in simulations with annotation-dependent MAF-dependent genetic architectures using all simulated GWAS samples and twice (1,000) the default reference panel sample size.** 10% of SNPs were randomly selected to be causal. Shrinkage level, $\alpha$, was set to 0.5. **a**) Estimates of $\lambda^2(C)$ for binary functional annotations. **b**) Estimates of $\lambda^2(C)$ for quintiles of continuous-valued annotations. Mean and standard errors were obtained across 1,000 simulations. Error bars represent 1.96 times the standard error on both sides. Numerical results are reported in Table S14b.

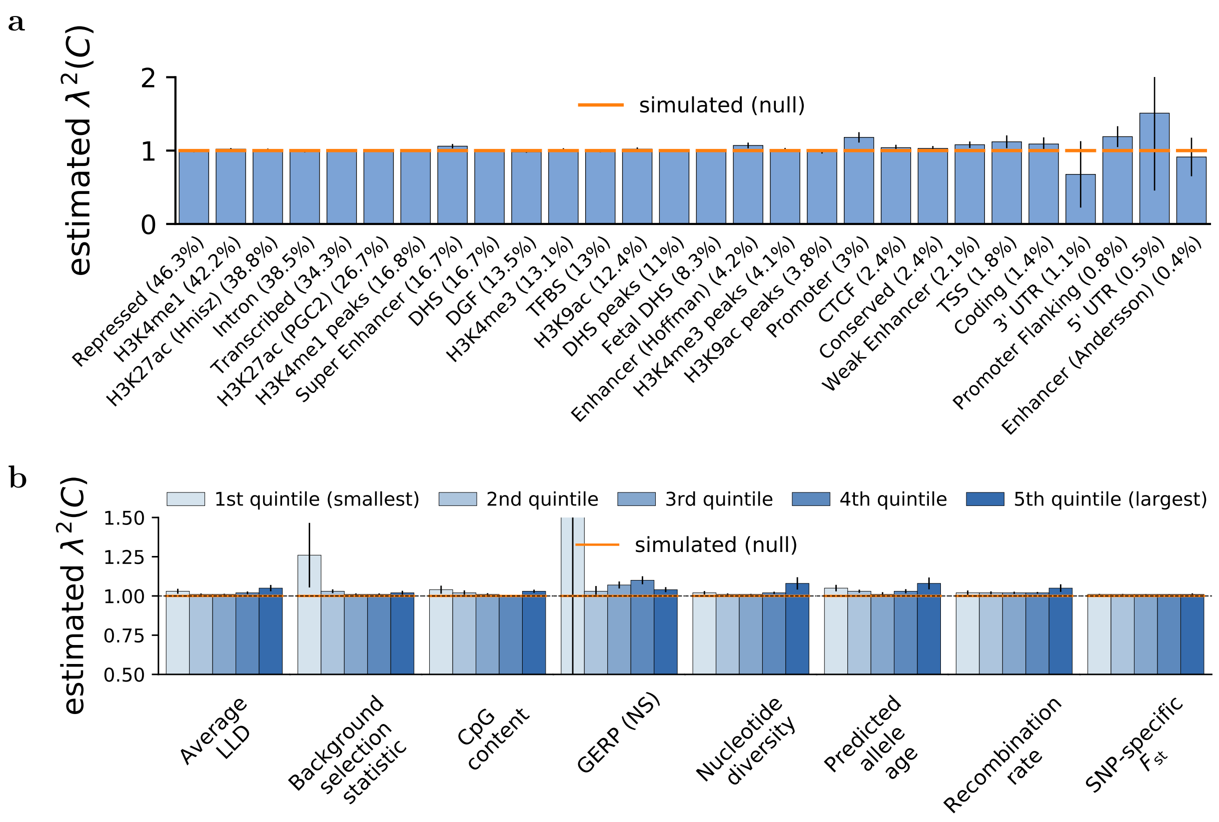


Figure S16: **Accuracy of S-LDXR in estimating enrichment of stratified squared trans-ethnic genetic correlation, $\lambda^2(C)$, in simulations under the baseline-LD-X model using half of the simulated GWAS samples and the default (500) reference panel sample size.** 10% of SNPs were randomly selected to be causal. Shrinkage level, $\alpha$, was set to 0.5. **a**) Estimates of $\lambda^2(C)$ for binary functional annotations. **b**) Estimates of $\lambda^2(C)$ for quintiles of continuous-valued annotations. Mean and standard errors were obtained across 1,000 simulations. Error bars represent 1.96 times the standard error on both sides. Numerical results are reported in Table S15a.

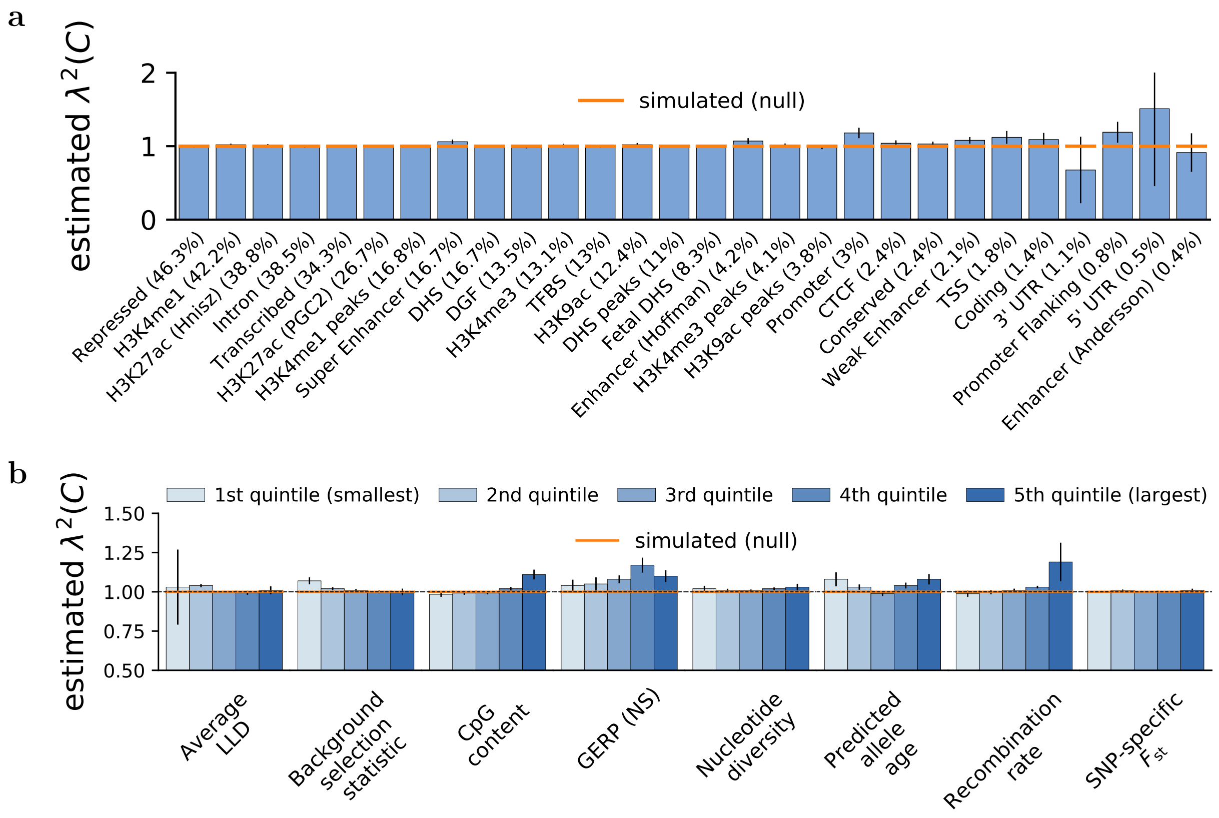


Figure S17: **Accuracy of S-LDXR in estimating enrichment of stratified squared trans-ethnic genetic correlation, $\lambda^2(C)$, in simulations with annotation-dependent MAF-dependent genetic architectures using half of the simulated GWAS samples and the default (500) reference panel sample size.** 10% of SNPs were randomly selected to be causal. Shrinkage level, $\alpha$, was set to 0.5. **a**) Estimates of $\lambda^2(C)$ for binary functional annotations. **b**) Estimates of $\lambda^2(C)$ for quintiles of continuous-valued annotations. Mean and standard errors were obtained across 1,000 simulations. Error bars represent 1.96 times the standard error on both sides. Numerical results are reported in Table S15b.

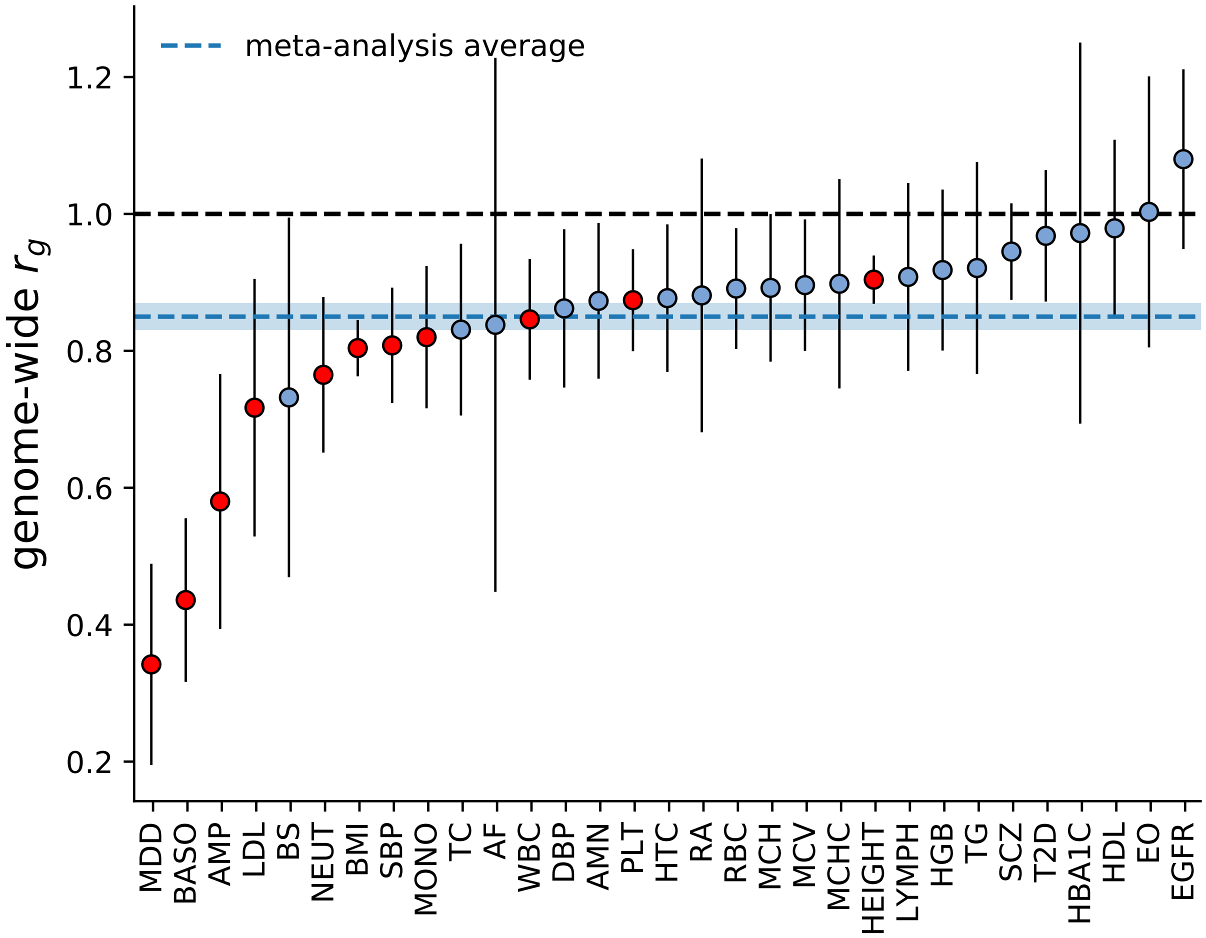


Figure S18: **Genome-wide trans-ethnic genetic correlation for 31 diseases and complex traits.** Diseases and complex traits are sorted by the magnitude of trans-ethnic genetic correlation. Traits with estimated trans-ethnic genetic correlation significantly less than 1 (one-tailed $p < 0.05/31$) are marked by red filled dots. Full name of the traits can be found in Table S16. Error bars represent 1.96 times the jackknife standard error on each side. The blue dashed line represents meta-analyzed $r_g$, and the shaded region covers 1.96 times the meta-analysis standard error on each side.

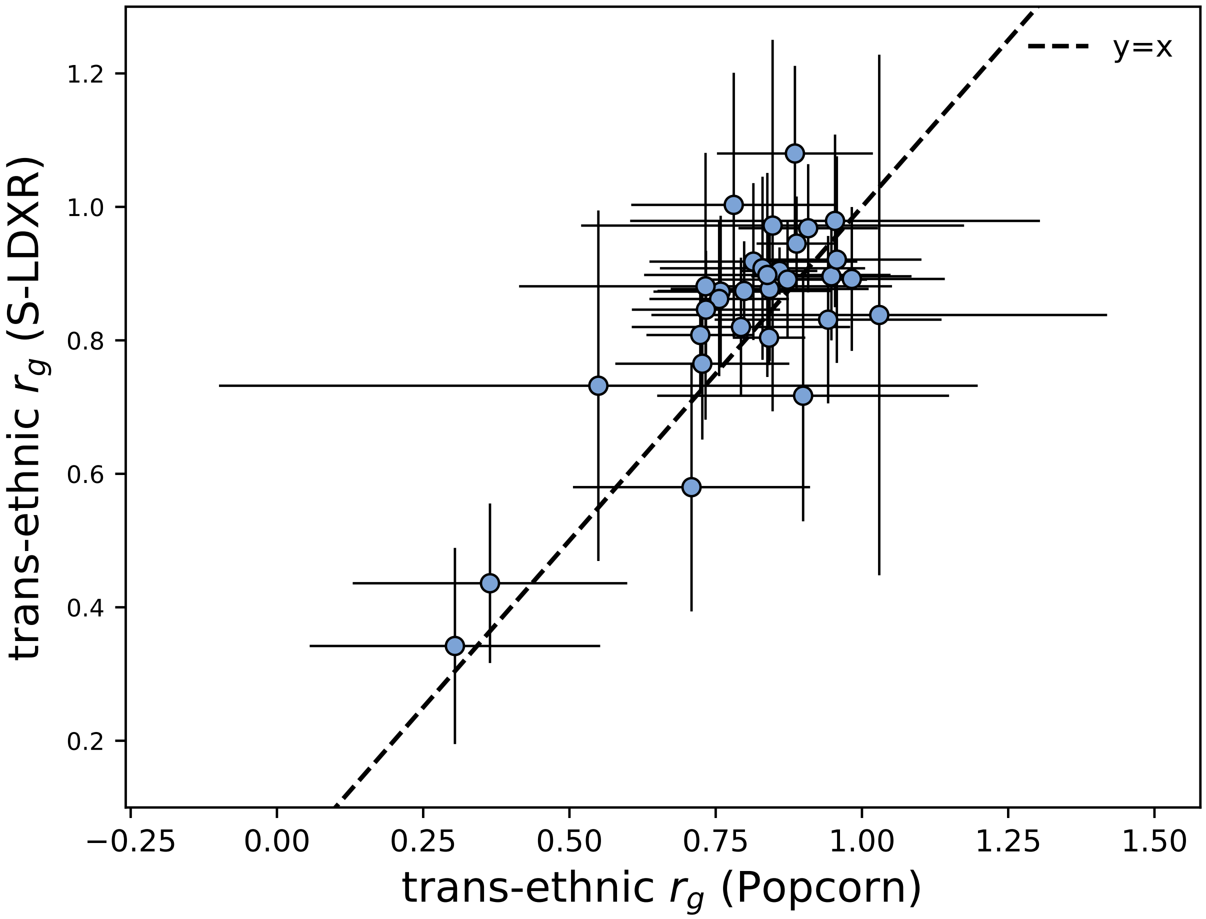


Figure S19: **Comparison of S-LDXR vs. Popcorn[2] estimates of genome-wide trans-ethnic genetic correlations for 31 diseases and complex traits.** Error bars represent 1.96 times the standard error on both sides. The meta-analyze average $r_g$ of S-LDXR and Popcorn are 0.85 (s.e. 0.01) and 0.82 (s.e. 0.01), respectively.

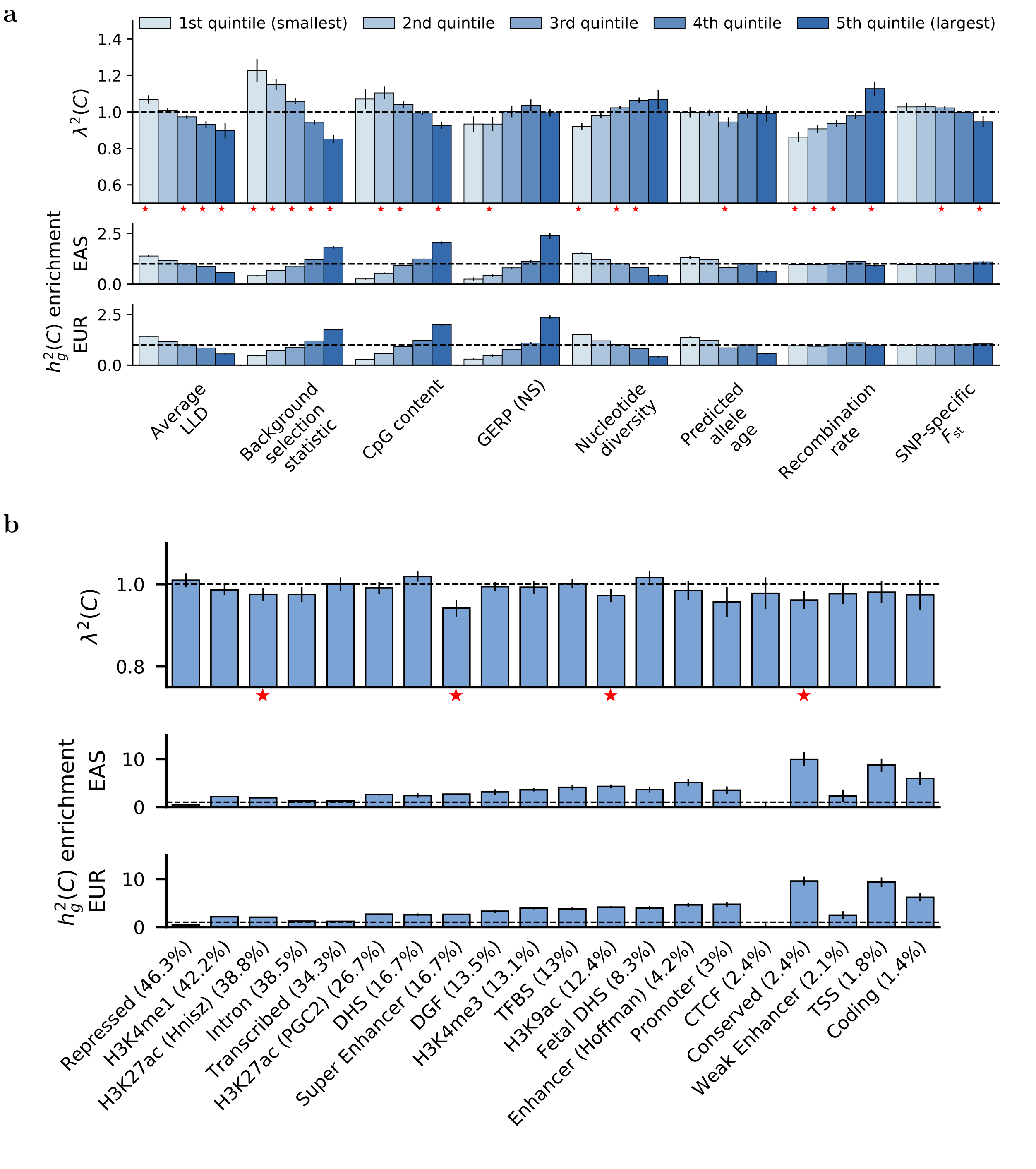


Figure S20: **Enrichment of stratified squared trans-ethnic genetic correlation, $\lambda^2(C)$, across a) quintiles of continuous-valued annotations and b) functional annotations.** The shrinkage level, $\alpha$, was set to 1.0. Error bars represent 1.96 times the standard error on both sides.

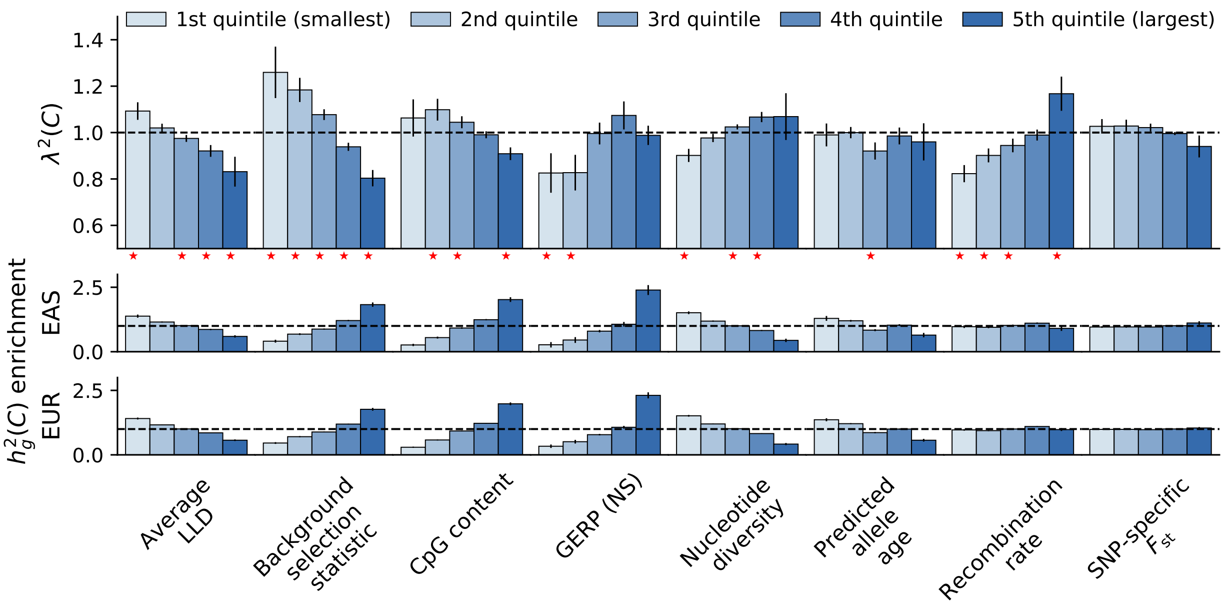


Figure S21: **S-LDXR results for quintiles of 8 continuous-valued annotations across 20 approximately independent diseases and complex traits.** The shrinkage parameter, $\alpha$, was set to 0.5. Error bars represent 1.96 times the standard error on both sides.

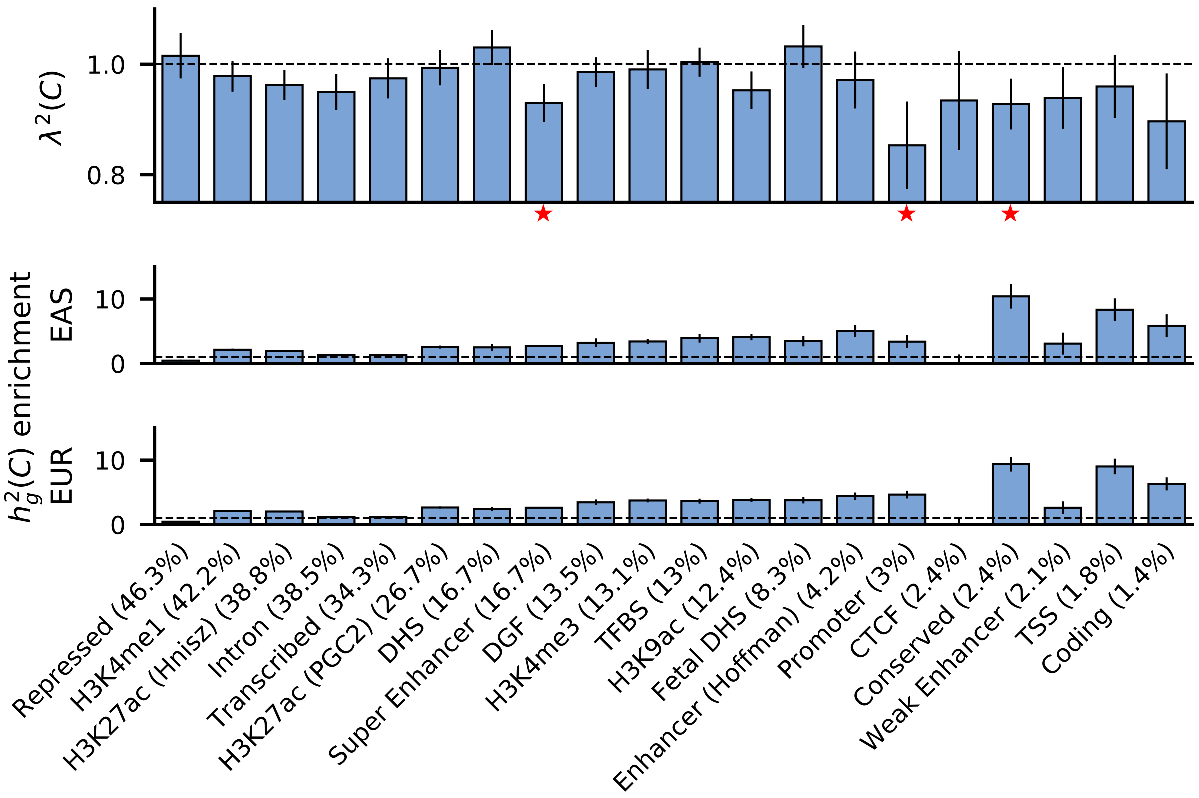


Figure S22: **S-LDXR results for 20 binary functional annotations across 20 approximately independent diseases and complex traits.** The shrinkage parameter, $\alpha$, was set to 0.5. Error bars represent 1.96 times the standard error on both sides.

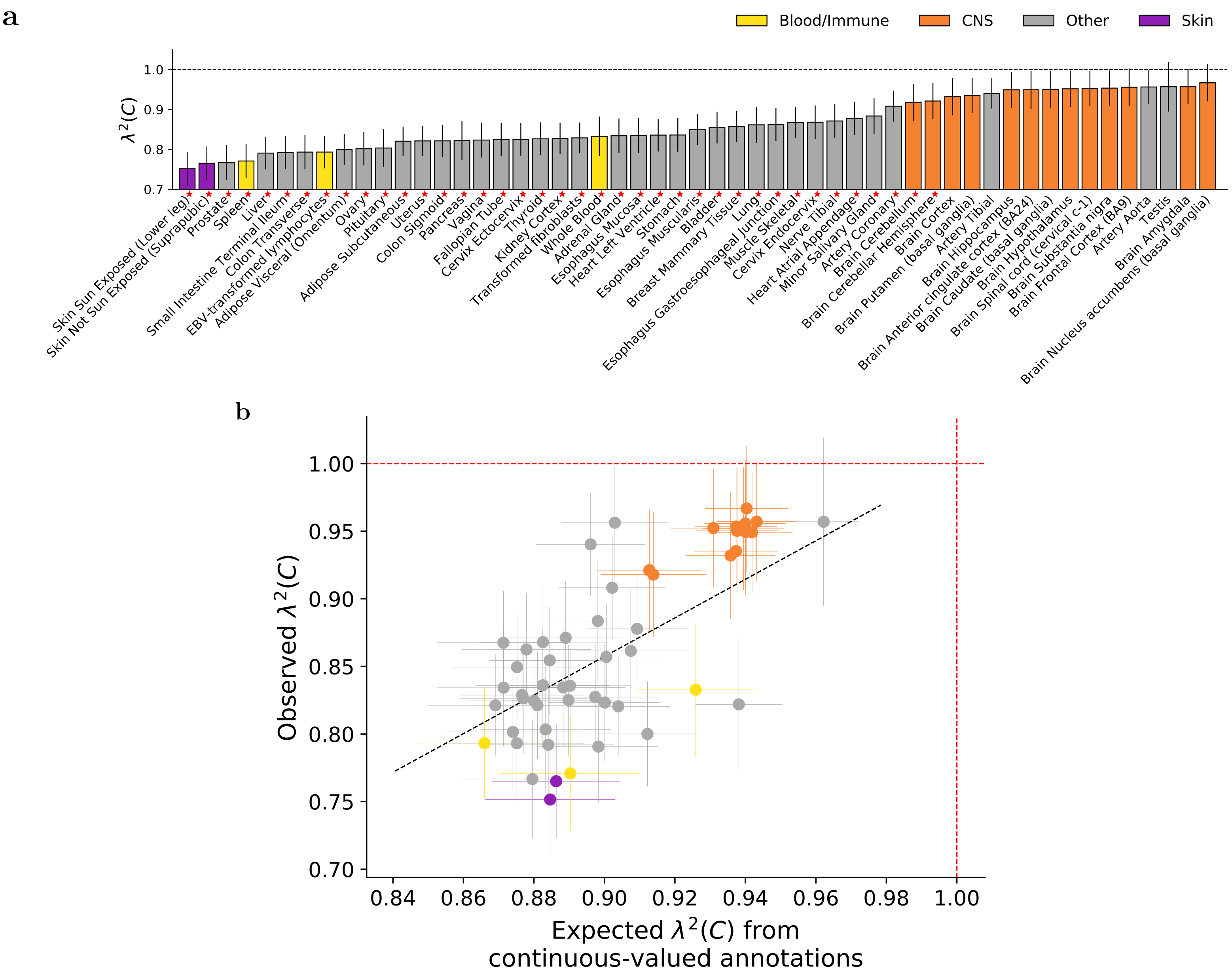


Figure S23: **S-LDXR results for 53 specifically expressed gene (SEG) annotations across 31 diseases and complex traits in analyses with the shrinkage parameter $\alpha$ set to 0.0.** (a) We report estimates of the enrichment/depletion of squared trans-ethnic genetic correlation ($\lambda^2(C)$) for each SEG annotation (sorted by $\lambda^2(C)$). Results are meta-analyzed across 31 diseases and complex traits. Error bars denote $\pm 1.96\times$ standard error. Red stars ($\star$) denote two-tailed p<0.05/53. Numerical results are reported in Table S21. (b) We report observed $\lambda^2(C)$ vs. expected $\lambda^2(C)$ based on 8 continuous-valued annotations, for each SEG annotation. Results are meta-analyzed across 31 diseases and complex traits. Error bars denote $\pm 1.96\times$ standard error. Annotations are color-coded as in (a). The dashed black line (slope=1.40) denotes a regression of observed $\lambda(C) - 1$ vs. expected $\lambda(C) - 1$ with intercept ($R = 0.74$) constrained to 0. Numerical results including population-specific heritability enrichment estimates are reported in Table S23.

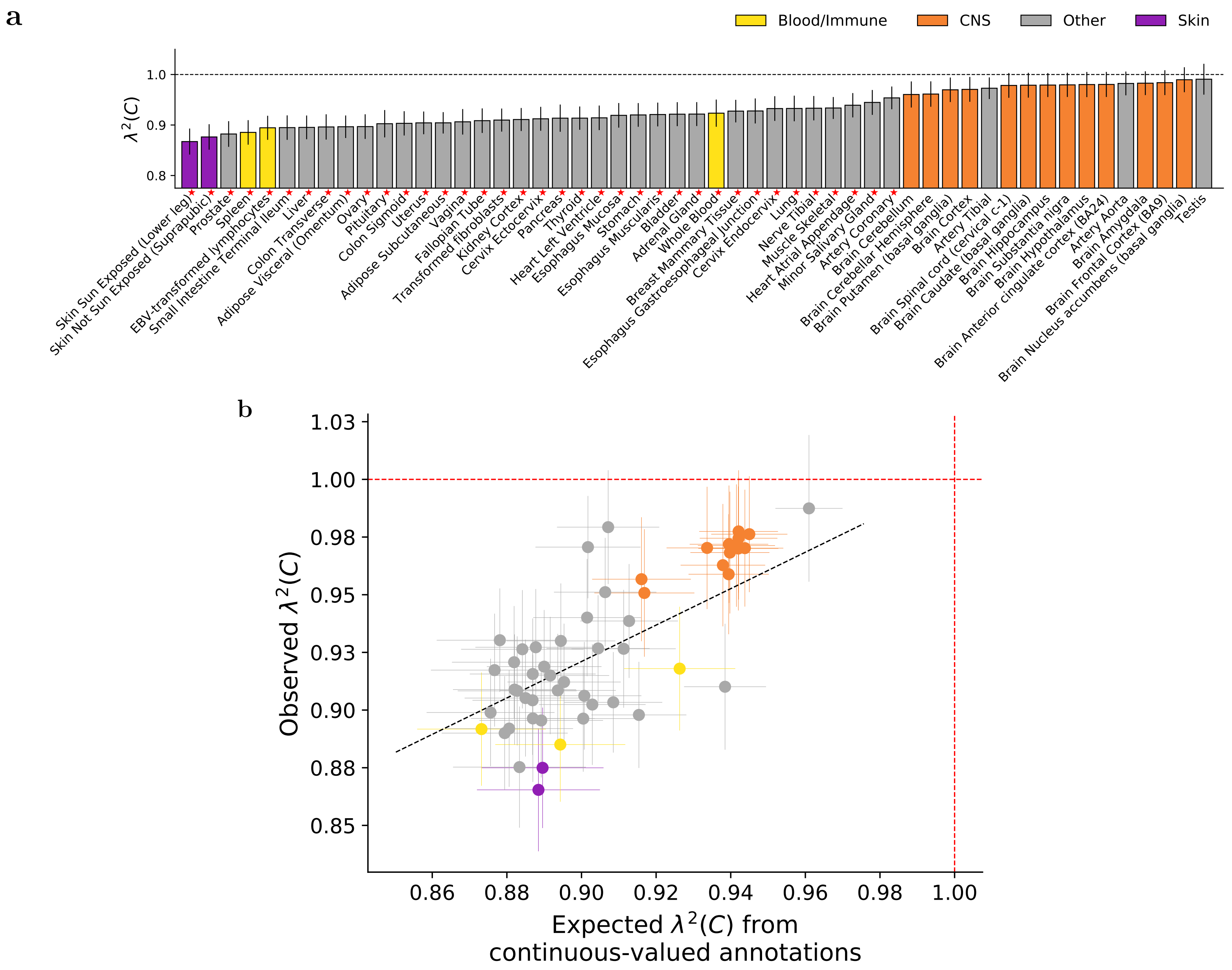


Figure S24: **S-LDXR results for 53 specifically expressed gene (SEG) annotations across 31 diseases and complex traits in analyses with the shrinkage parameter $\alpha$ set to 1.0.** (a) We report estimates of the enrichment/depletion of squared trans-ethnic genetic correlation ($\lambda^2(C)$) for each SEG annotation (sorted by $\lambda^2(C)$). Results are meta-analyzed across 31 diseases and complex traits. Error bars denote $\pm 1.96\times$ standard error. Red stars ($\star$) denote two-tailed p<0.05/53. Numerical results are reported in Table S21. (b) We report observed $\lambda^2(C)$ vs. expected $\lambda^2(C)$ based on 8 continuous-valued annotations, for each SEG annotation. Results are meta-analyzed across 31 diseases and complex traits. Error bars denote $\pm 1.96\times$ standard error. Annotations are color-coded as in (a). The dashed black line (slope=0.76) denotes a regression of observed $\lambda(C) - 1$ vs. expected $\lambda(C) - 1$ ($R = 0.76$) with intercept constrained to 0. Numerical results including population-specific heritability enrichment estimates are reported in Table S23.

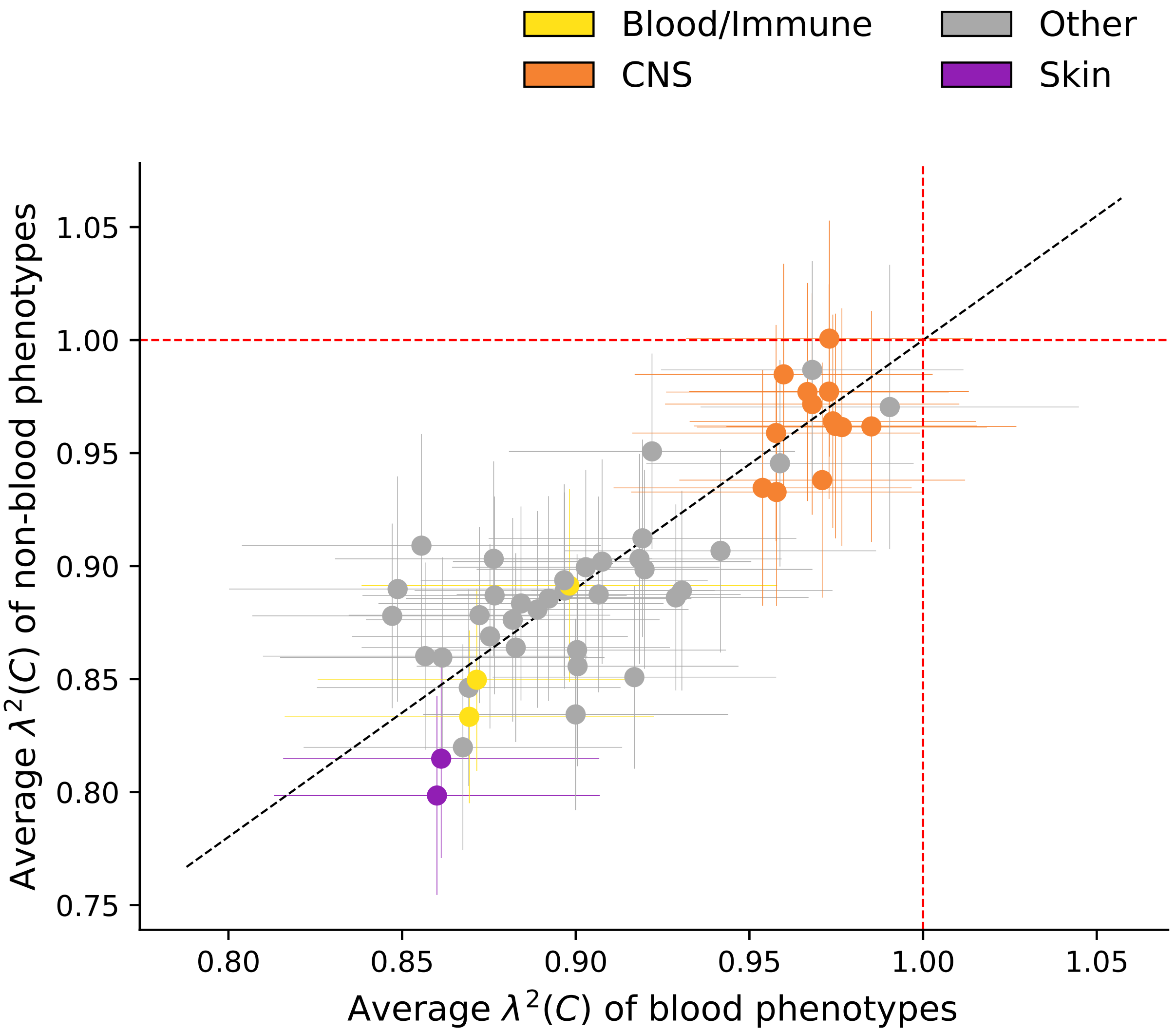


Figure S25: **Comparison of S-LDXR results for 53 specifically expressed gene (SEG) annotations for 14 blood-related traits vs. 17 other traits.** The list of 14 blood phenotypes is: BASO, EO, HBA1C, HGB, HTC, LYMPH, MCH, MCHC, MCV, MONO, NEUT, PLT, RBC, WBC. The list of 16 non-blood phenotype is: AF, AMN, AMP, BMI, BS, DBP, EGFR, HEIGHT, HDL, LDL, MDD, RA, SBP, SCZ, TC, TG, T2D. Full name of the abbreviations can be found in Table S16. Here, the shrinkage parameter was set to 0.5. Error bars represent 1.96 times the standard error on both sides. The black dashed line represent the regression line (slope=1.10) fitting $(\lambda^2(C) - 1)$ of non-blood phenotypes and $(\lambda^2(C) - 1)$ of blood phenotypes, with intercept constrained to 0.

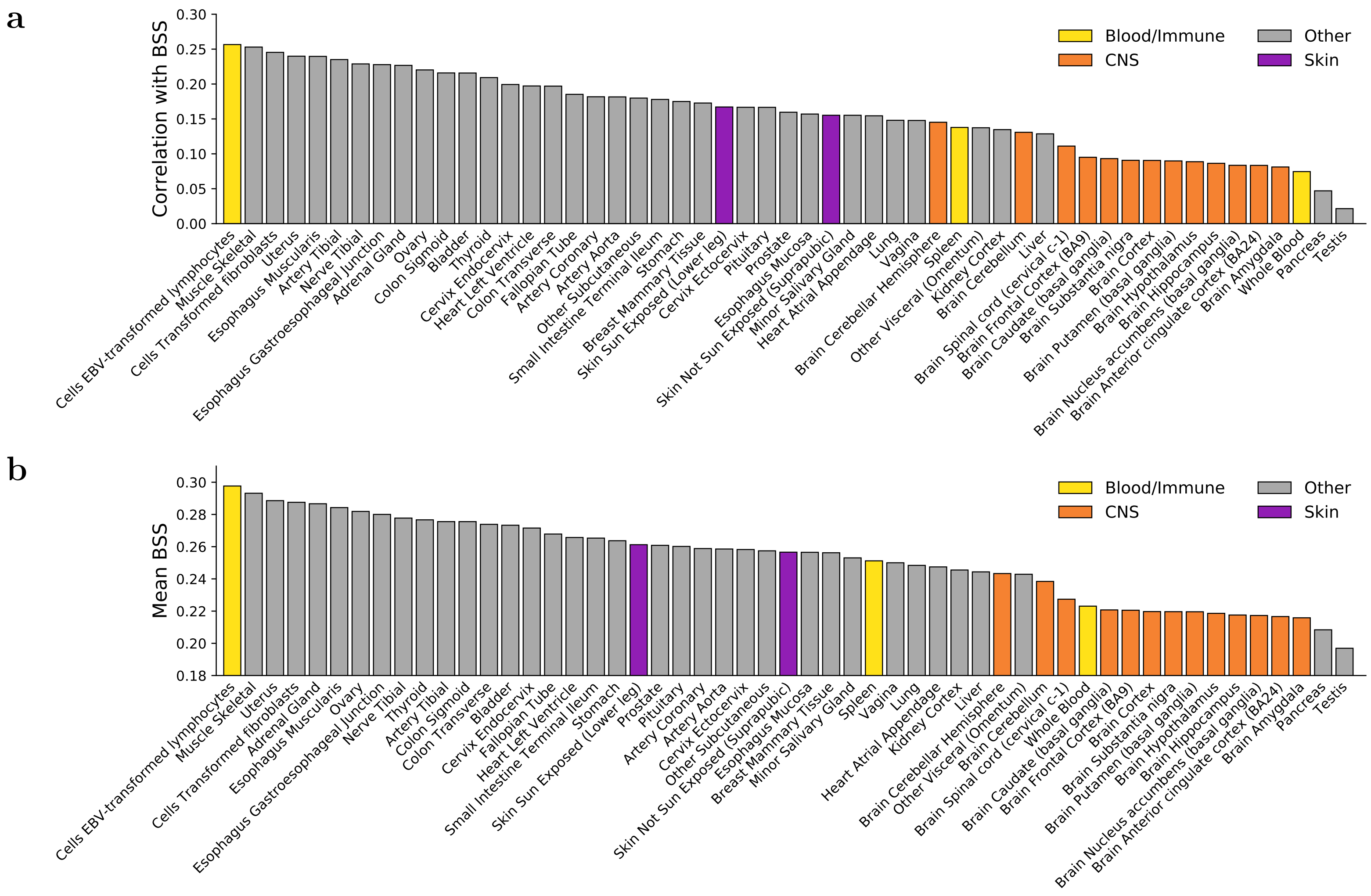


Figure S26: **Relationship between specifically expressed gene (SEG) annotations and background selection statistic (BSS). a)** Correlation between 53 SEG annotations and BSS. Tissues are ranked by magnitude of correlation. **a)** Mean BSS at annotated SNPs for 53 SEG annotations. Tissues are ranked by magnitude of the mean.

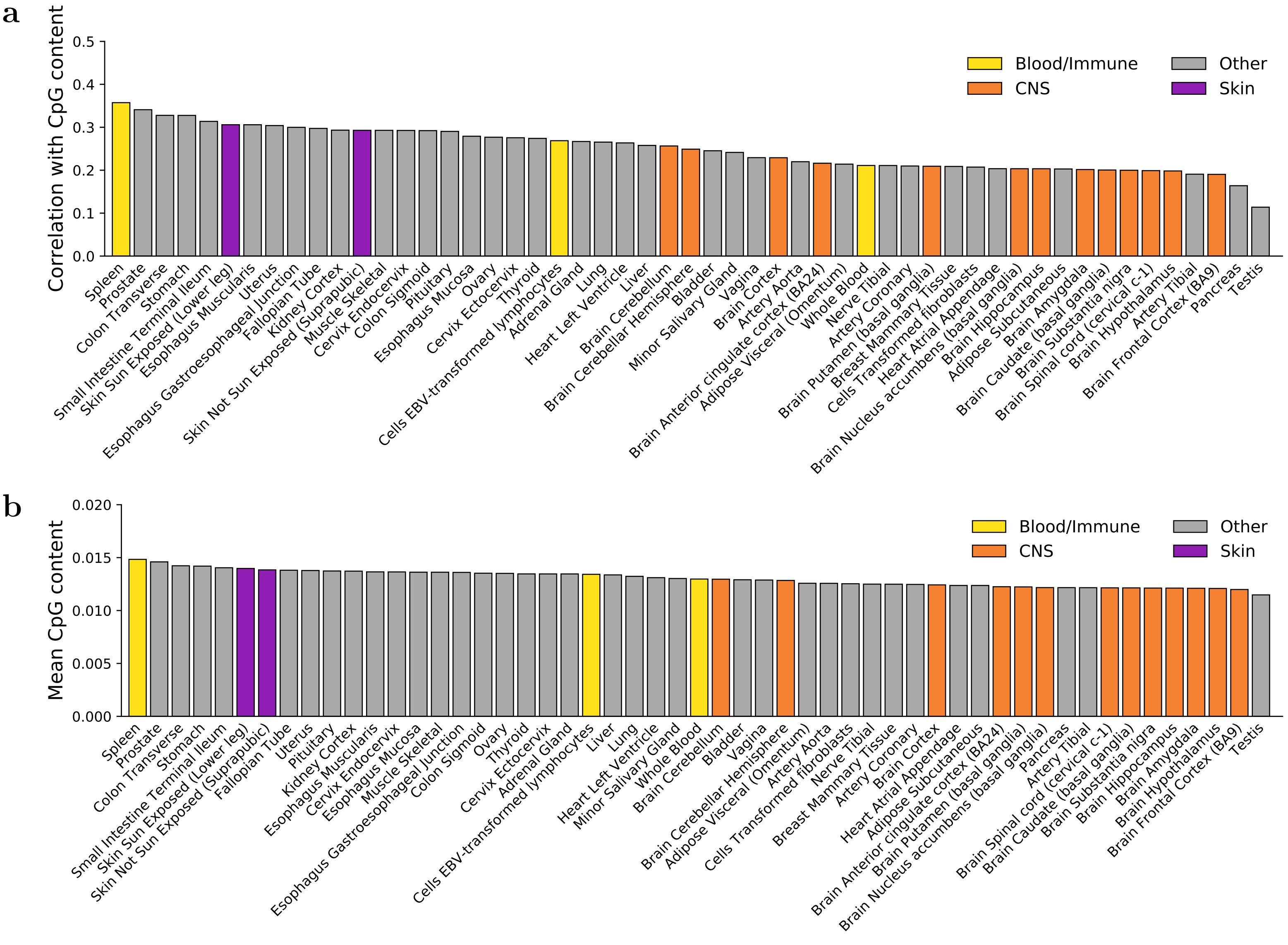


Figure S27: **Relationship between specifically expressed gene (SEG) annotations and CpG content. a)** Correlation between 53 SEG annotations and CpG content. Tissues are ranked by magnitude of correlation. **a)** Mean CpG content at annotated SNPs for 53 SEG annotations. Tissues are ranked by magnitude of the mean.

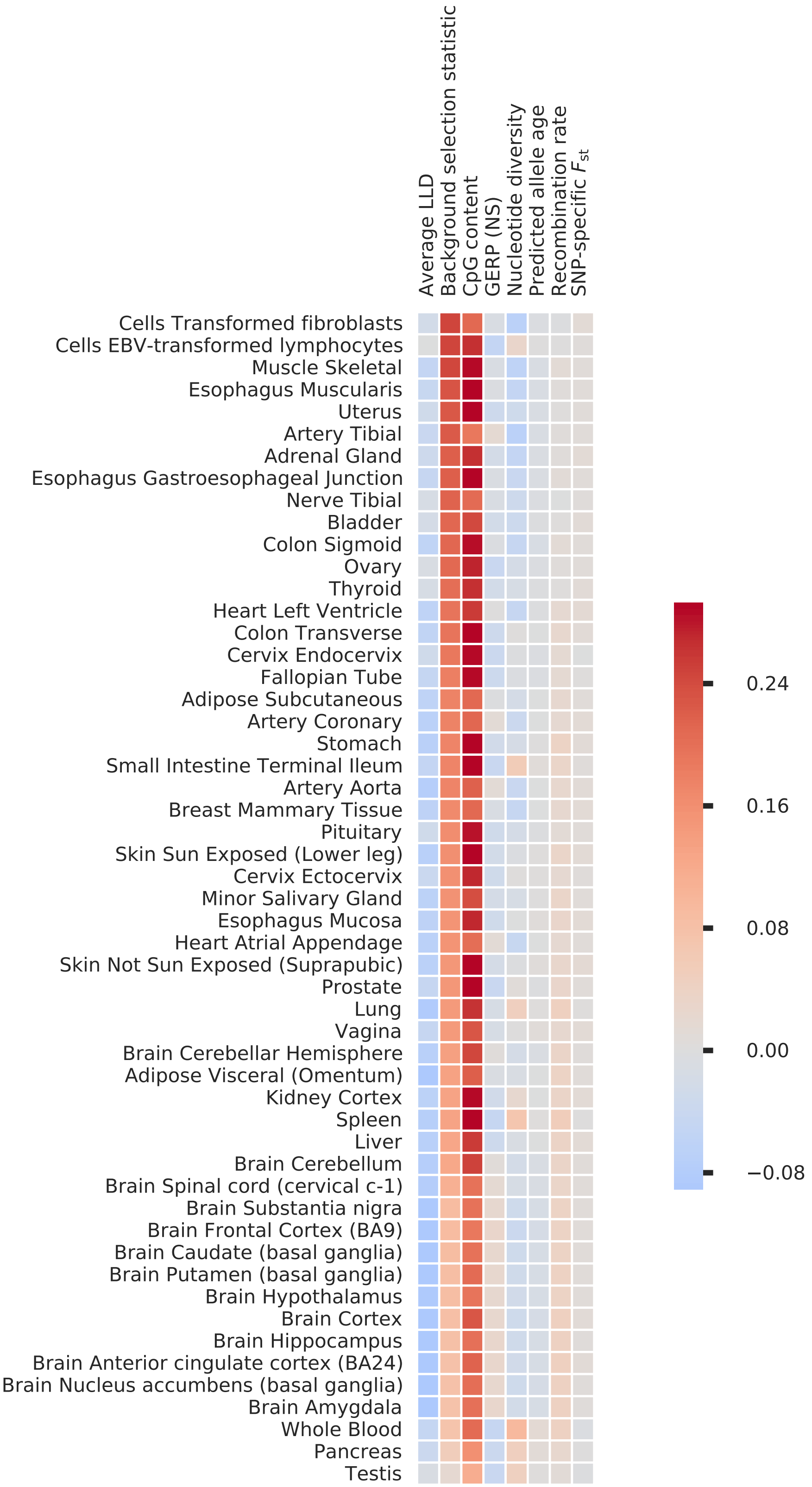


Figure S28: **Correlation between specifically expressed gene annotations and 8 continuous-valued annotations.** Annotations are sorted inversely based on their correlation with background selection statistic.

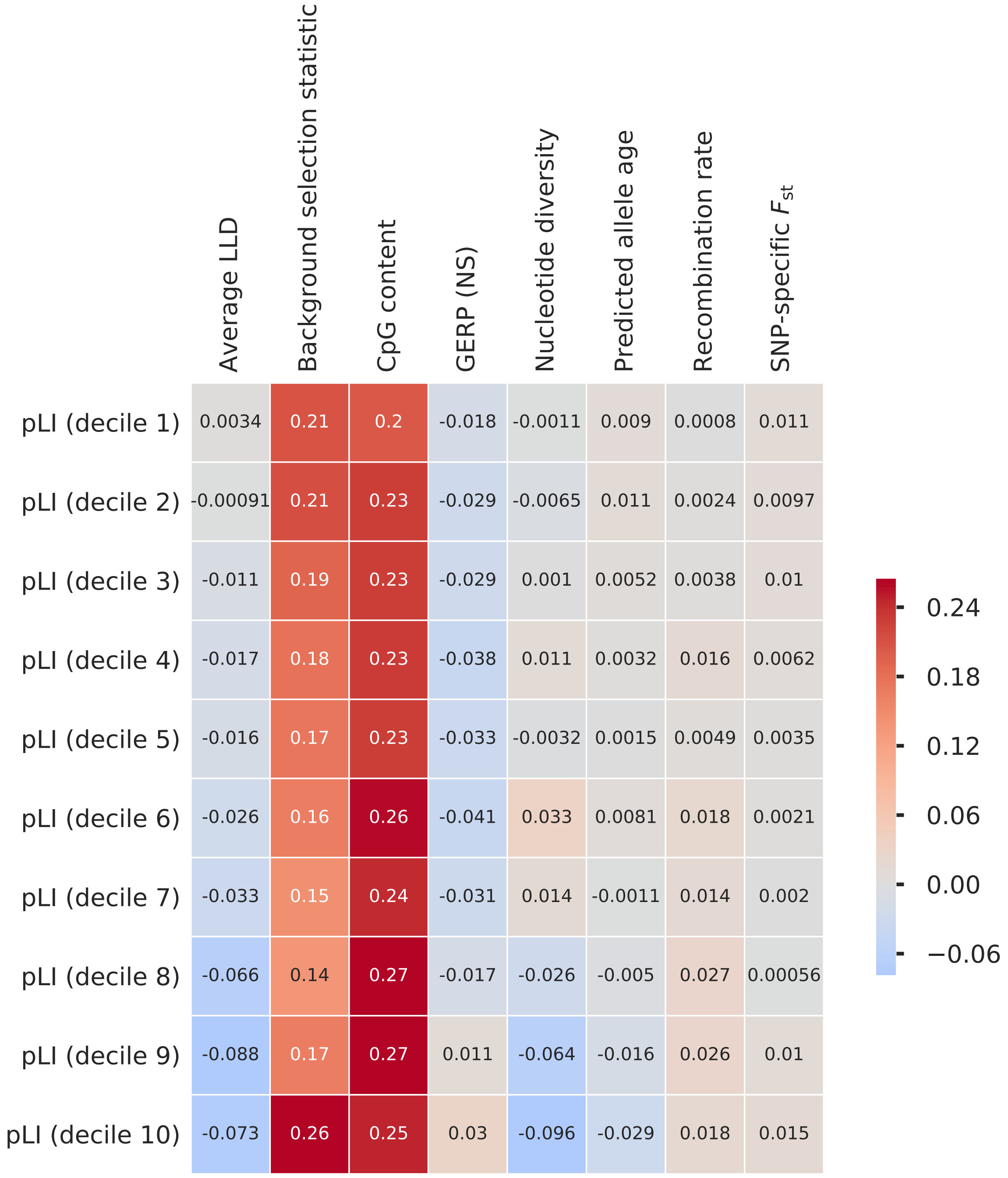


Figure S29: **Correlation between probability of loss-of-function intolerance (pLI) decile gene annotations and 8 continuous-valued annotations.** Here, correlations are calculated across all SNPs with minor allele frequency greater than 5% in both East Asian and European populations.

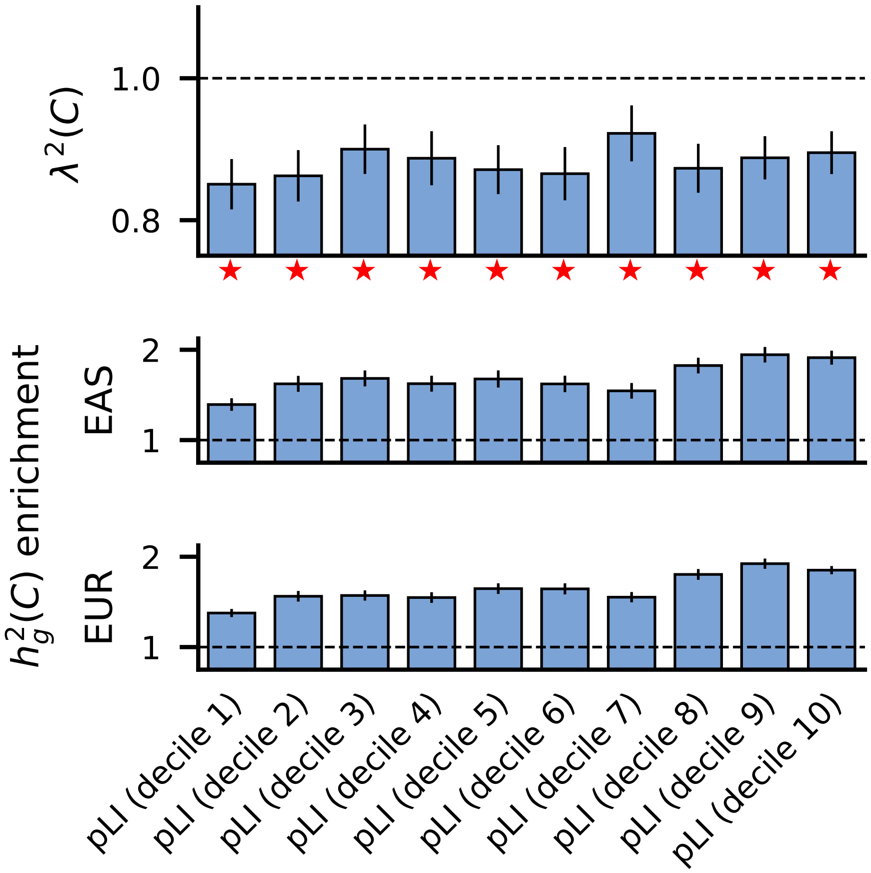


Figure S30: **S-LDXR results for deciles of probability of loss-of-function intolerance (pLI) annotations.** Deciles with $\lambda^2(C)$ significantly less than 1 are marked by red stars. Numerical results can be found in Table S26.

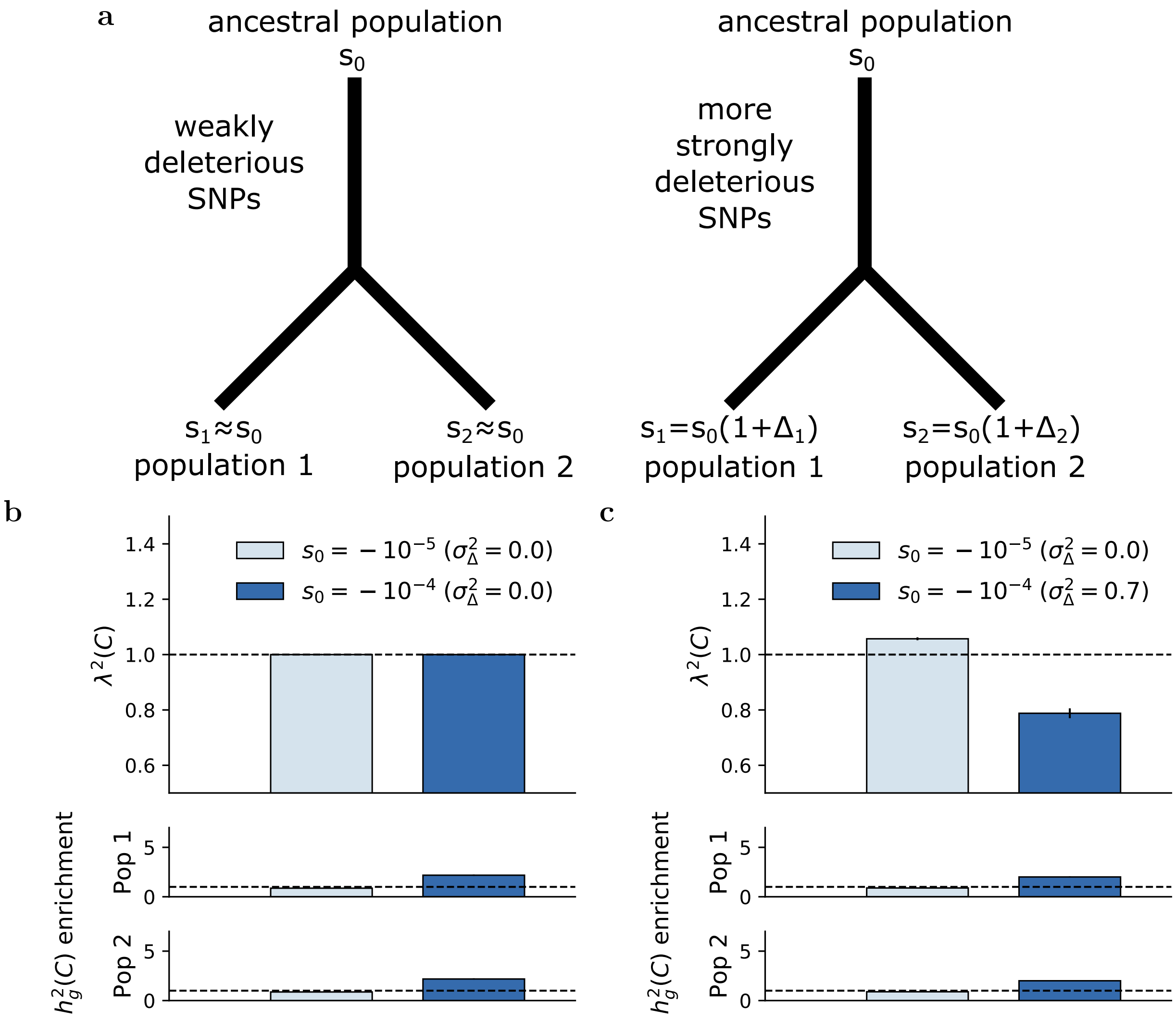


Figure S31: **Evolutionary modeling results using 2-population extension of Eyre-Walker model.** We use negative $s$ to denote deleteriousness, following convention of previous works.[11,12] However, positive $s$ (i.e. beneficial mutations) may also be plausible. (a) Diagrams illustrating fitness effects in population 1 and population 2 ($s_1$ and $s_2$) as a function of the fitness effect in the ancestral population ($s_0$) at weakly deleterious SNPs (left; e.g. corresponding to SNPs in bottom quintile of background selection statistic) and more strongly deleterious SNPs (right; e.g. corresponding to SNPs in top quintile of background selection statistic). (b), (c) We report enrichment/depletion of squared trans-ethnic genetic correlation ($\lambda^2(C)$) for SNPs with different fitness effects, in simulations under a two-population extension of the Eyre-Walker model with (b) $\sigma^2_\Delta = 0$ for both weakly deleterious SNPs ($s_0 = -10^{-5}$) and more strongly deleterious SNPs ($s_0 = -10^{-4}$), (c) $\sigma^2_\Delta = 0.0$ for weakly deleterious SNPs and $\sigma^2_\Delta = 0.7$ for more strongly deleterious SNPs. Results are averaged across 1,000 simulations. Error bars denote $\pm 1.96\times$ standard error. Numerical results are reported in Table S27.

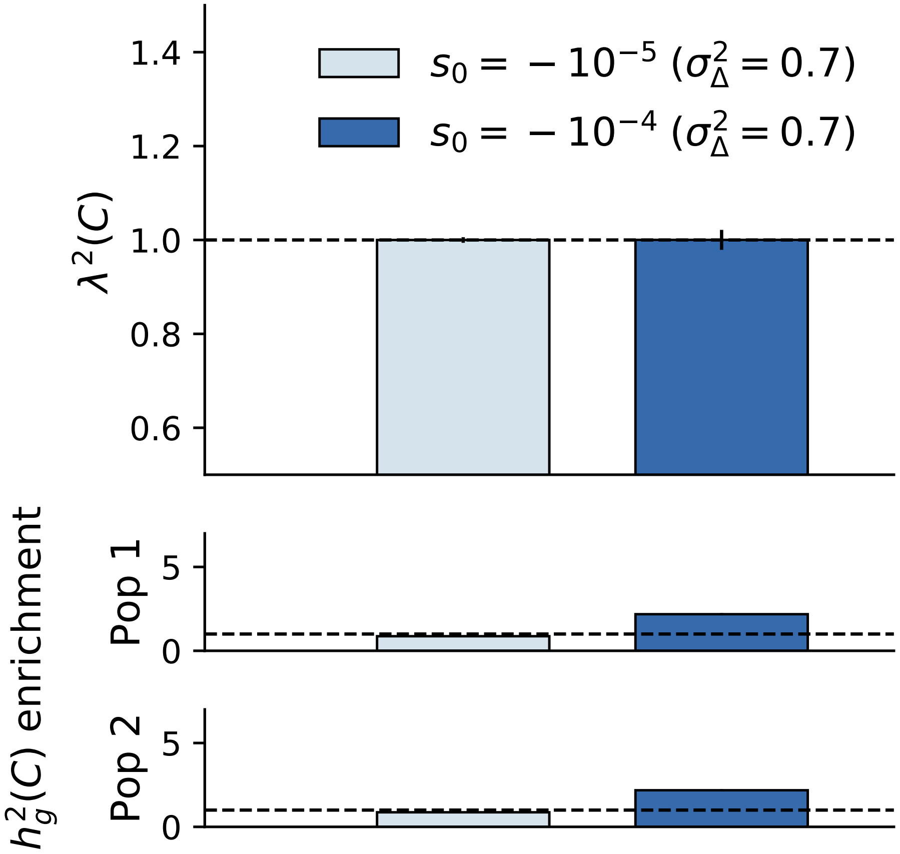


Figure S32: **Evolutionary modeling results using 2-population extension of Eyre-Walker model with $\sigma^2_\Delta = 0.7$ for both weakly deleterious and more strongly deleterious SNPs.** We use negative $s$ to denote deleteriousness, following convention of previous works.[11,12] However, positive $s$ (i.e. beneficial mutations) may also be plausible. We report enrichment/depletion of squared trans-ethnic genetic correlation ($\lambda^2(C)$) for SNPs with different fitness effects, in simulations under a two-population extension of the Eyre-Walker model,[10] with $\sigma^2_\Delta = 0.7$ for both weakly deleterious and more strongly deleterious SNPs. Results are averaged across 1,000 simulations. Error bars denote $\pm 1.96\times$ standard error.
